# Evolutionary origins of epidemic potential among human RNA viruses

**DOI:** 10.1101/771394

**Authors:** Lu Lu, Liam Brierley, Gail Robertson, Feifei Zhang, Samantha Lycett, Donald Smith, Margo Chase-Topping, Peter Simmonds, Mark Woolhouse

## Abstract

To have epidemic potential, a pathogen must be able to spread in human populations, but of human-infective RNA viruses only a minority can do so. We investigated the evolution of human transmissibility through parallel analyses of 1755 virus genome sequences from 39 RNA virus genera. We identified 57 lineages containing human-transmissible species and estimated that at least 74% of these lineages have evolved directly from non-human viruses in other mammals or birds, a public health threat recently designated “Disease X”. Human-transmissible viruses rarely evolve from virus lineages that can infect but not transmit between humans. This result cautions against focussing surveillance and mitigation efforts narrowly on currently known human-infective virus lineages and supports calls for a better understanding of RNA virus diversity in non-human hosts.

## Introduction

Ebolavirus in 2014-15 and SARS coronavirus in 2003 are examples of infectious disease epidemics resulting from the emergence of RNA viruses from non-human reservoirs^1^. However, the majority of spill-overs from non-human reservoirs involve viruses that do not spread in human populations (e.g., the strictly zoonotic orthohantaviruses and bat lyssaviruses)^2^. There have been various studies of ecological predictors of virus infectivity to humans (e.g. ^3, 4^) but less attention has been paid to transmissibility in human populations even though this trait is integral to epidemic potential^5–7^. Here, we aim to identify phylogenetic and ecological characteristics of RNA virus lineages that are associated with the evolution of human transmissibility.

Members of over 200 non-reverse transcribing (non-RT) RNA virus species (i.e. excluding retroviruses) are known to infect humans^8, 9^. Human-infective viruses are found in 20 of the 22 currently recognised families of non-RT RNA viruses of mammals and/or birds (the exceptions being the *Arteriviridae* and *Birnaviridae*). Although members of only 82 human-infective non-RT RNA virus species (40%) are known to transmit within human populations (other than by vertical or iatrogenic routes)^8^, these too have broad taxonomic distributions, being found in 19 families (the *Bornaviridae* having no human-transmissible species).

New human RNA viruses are still discovered regularly^8^ and it is likely that a large number of currently unknown RNA viruses are circulating in non-human mammals and birds, some of which may have the potential to infect and be transmitted by humans^10–12^. All known human RNA viruses form part of purely mammal or mammal/bird virus clades^12^ and there is no evidence of human-infective RNA viruses emerging from other kinds of host^6^.

Human transmissibility (here, broadly defined as spread directly from person to person or via an indirect route, such as environmental contamination or an arthropod vector) is a necessary but not sufficient condition for a virus to have epidemic potential. Pathogens that are transmissible but with a basic reproduction number (*R*0) less than 1 in humans (e.g. Lassa virus) are restricted to self-limiting outbreaks and their long-term persistence requires a non-human reservoir. Previous studies have addressed the ecology^6^ and evolution^13^ of the transition between *R*0<1 and *R*0>1; here, we focus on the evolution of human transmissibility, i.e. *R*0>0. We categorised RNA viruses into three infectivity/transmissibility (IT) levels^6^: Level 1 (L1) viruses are infective to non-human mammals and/or birds but not to humans; L2 viruses are human-infective but not human-transmissible; and L3/4 viruses are human-infective and transmissible (combining viruses with 0<*R*0<1 (L3) and those with *R*0>1 (L4)).

Here, we propose a conceptual model of RNA virus emergence where that latent capacities to infect and transmit between humans are traits inherent to some L1 virus lineages circulating in non-human reservoirs (Figure 1); this has been termed “fortuitous adaptation”^14^. This view emphasizes the role of virus genetic diversity within the reservoir, with humans acting as “sentinels”^10^, revealing the presence of L2 or L3/4 viruses through contact with the reservoir. New L3/4 viruses can emerge by any of three routes: i) directly from viruses in a non-human reservoir (L1) at a rate given by parameter *a*; ii) step-wise via a strictly zoonotic (L2) phase in a non-human reservoir or, less probably, in infected humans at rates given by parameters *b* and *c*; or iii) diversification of human-transmissible (L3/4) virus lineages within humans or, possibly, a non-human reservoir. We note that human infectivity and transmissibility are products of convergent evolution, with their molecular and genetic determinants differing across virus taxa^9^, and that the rates of evolution of these traits represented by parameters *a*, *b* and *c* are likely to vary between lineages.

**Figure 1.**
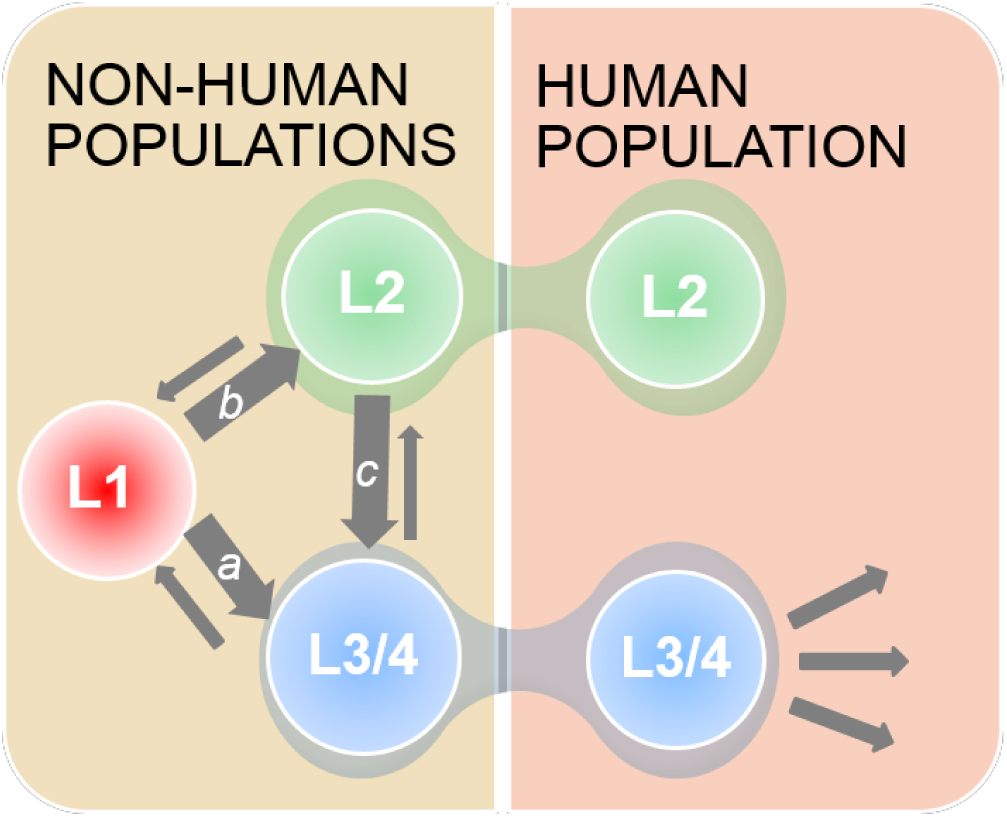
Conceptual model of non-RT RNA virus emergence into human populations. Viruses circulating in the non-human reservoir may not be infective to humans (L1) or have the potential to infect or infect and transmit between humans (L2 and L3/4 respectively). Virus lineages in the reservoir may acquire (thick grey arrows showing rates *a*, *b* and *c*) or lose (thin grey arrows) the capacities to infect and/or transmit in humans through genetic drift – these traits are not under direct selection in the non-human reservoir. L2 and L3/4 viruses in non-human hosts may cross the species barrier and enter human populations. L3/4 (but not L2) viruses may then spread and evolve independently in the human population (multiple grey arrows). Phylogenetic analyses identify past transitions between L1 and L3/4, L1 and L2 and L2 and L3/4 viruses and can be used to estimate the relative values of parameters *a*, *b* and *c*.

## Results

For phylogenetic analysis we used polymerase protein sequences of human and non-human RNA viruses classified into 39 genera (see Supplementary Materials). We only considered genera containing human-infective virus species, noting that not all eligible genera had sufficient sequences available for analysis. We also excluded all retroviruses *a priori* and the *Influenza A virus* genus *a posteriori* from the main analysis (see Supplementary Materials) but, given their importance, we performed separate analyses of these taxa.

Of the 1755 polymerase gene sequences used in our main study, 737 were from L1 viruses, 466 were from L2 viruses (90 from humans, 376 from non-human hosts) and 552 from L3/4 viruses (400 from humans, 152 from non-human hosts). These included virus sequences from natural infections of all available mammal or bird hosts. A plot of the cumulative number of sequences from L1, L2 and L3/4 viruses in our data set over time (Figure S1), reveals the rapid recent increase in numbers of L1 virus sequences that makes our study feasible at this time. The sequences represent 79% and 81% respectively of L2 and L3/4 non-RT RNA virus species currently listed by the International Committee for the Taxonomy of Viruses (ICTV)^9^. Within some virus species, we differentiated between recognised subtypes that differ in infectivity/transmissibility (IT) level (see Data File 4).

We estimated the phylogenies of the mammal/bird virus clade for each of the 39 non-RT RNA virus genera (Figure 2). We then used a Bayesian discrete trait approach (see Supplementary Materials) to estimate IT level for every node within those phylogenies. Our primary interest was tree topology; we did not attempt to date the nodes. The phylogenies indicated that for at least 22/39 genera containing human-infective viruses (L2 and/or L3/4), human infectivity is unlikely (P<0.20) to be an ancestral trait, and for at least 22/31 genera containing human-transmissible (L3/4) viruses, transmissibility is unlikely (P<0.20) to be an ancestral trait (Figures 3a, S2, Data File 1). In total, we identified 77 distinct lineages of human-infective (L2 and/or L3/4) and 57 of human-transmissible (L3/4) viruses (Figure 2, Data File 1).

**Figure 2.**
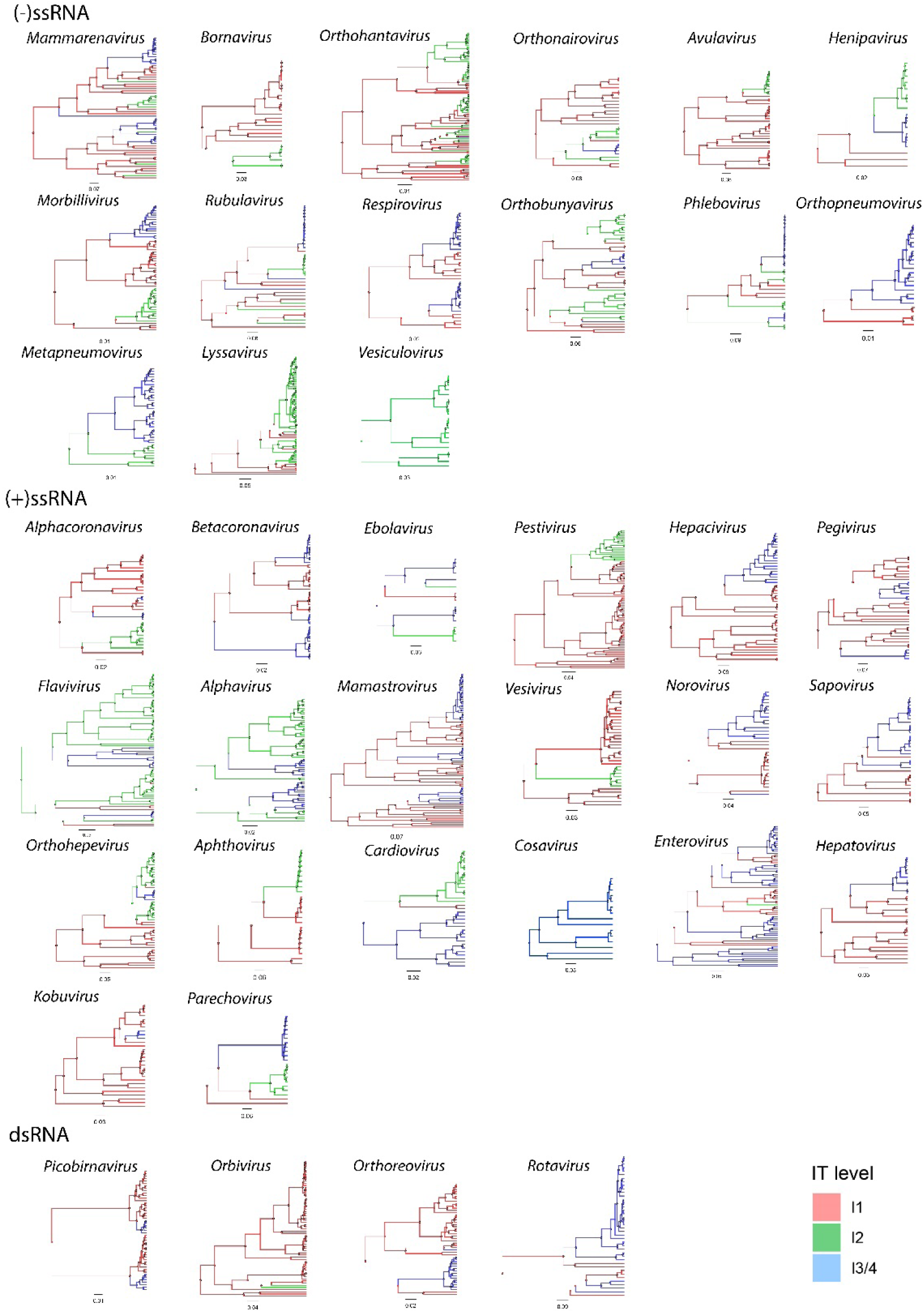
Bayesian maximum clade credibility (MCC) trees for members of 39 virus genera showing the most probable transitions between non-human viruses (red), viruses infective to humans (green) and viruses transmissible in human populations (blue). Phylogenies are arranged by genome type. Raw data are provided in Data File 5.

**Figure 3.**
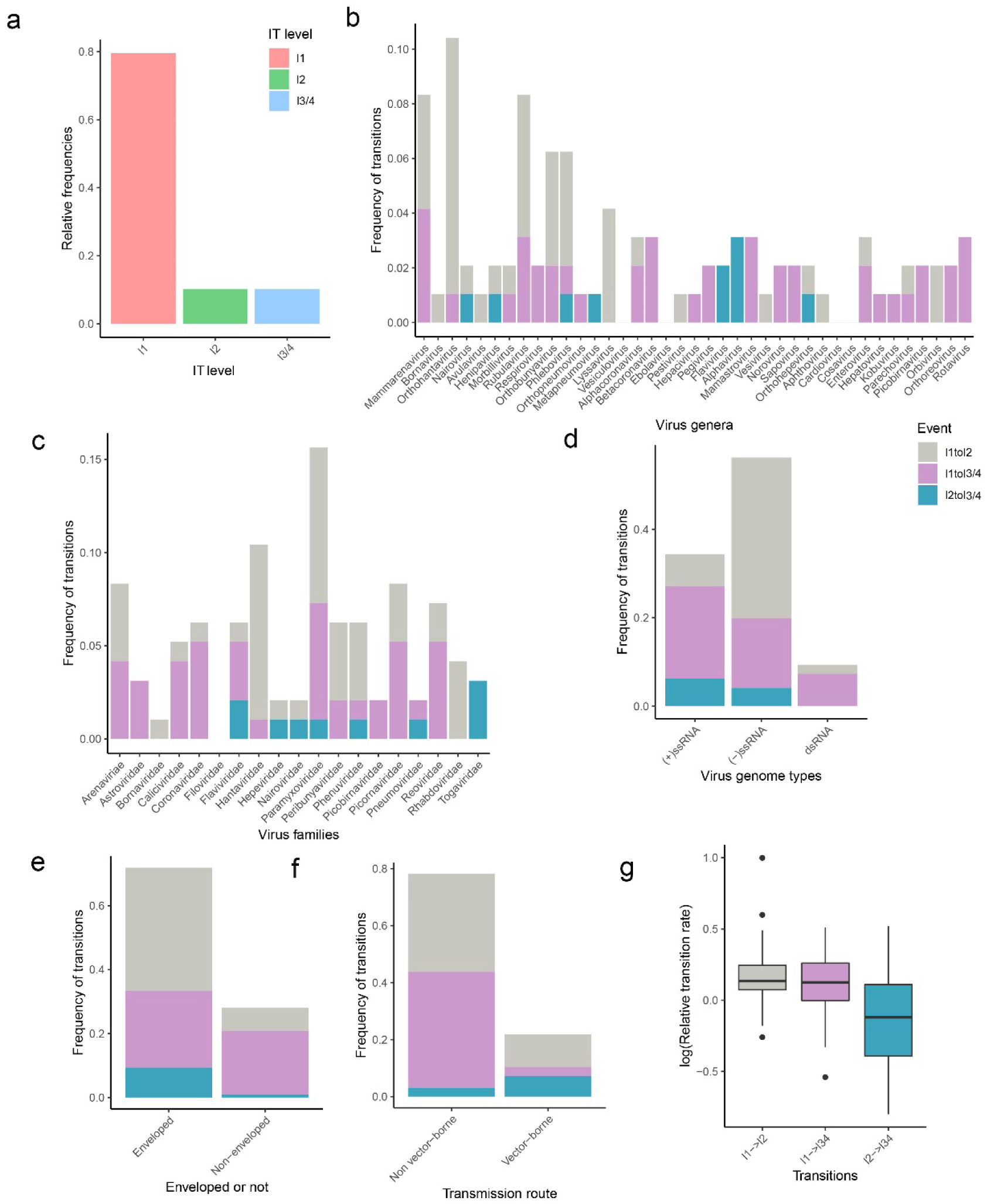
Outputs of discrete trait analyses. **a**) Frequency of estimated infection/transmission (IT) level (L1, L2 or L3/4) for the ancestral nodes of the mammal/bird clade in each genus (see Figure 2). **b**) Frequency of forward transitions by virus genus and whether enveloped. Three transitions are distinguished: L1 to L2 (grey), L1 to L3/4 (purple) and L2 to L3/4 (cyan). N=96. Vector-borne genera indicated with asterisks. **c**) As **b** by virus family. **d**) As **b** by virus genome type. **e**) As **b** by enveloped/non-enveloped. **f**) As **b** by vector-borne/non-vector-borne. **g**) Boxplots showing distribution of genus-level, loge-transformed relative transition rates for level transitions: L1-L2 (N=21), L1-L3/4 (N=26), L2-L3/4 (N=18). The rates used in the plot are shown in Figure S6.

We found evidence of 86 transitions involving gain of human infectivity and 52 involving gain of human transmissibility within the 39 virus genera (Table S1, Figures 2, S3, Data File 2). Human infectivity and transmissibility are likely to have evolved more than once in 20 and 17 genera respectively (Figure 3b). We note that we also found evidence of backward transitions: seven involving loss of human transmissibility and 11 involving loss of human infectivity (Data File 3). We compared our estimated numbers of transitions with outputs from both Markov jumps and a parsimony reconstruction method (see Supplementary Materials). These checks confirm that our estimates of the total numbers of transitions are well supported by the phylogenetic data, even though the precise location of a transition within a phylogeny could not always be estimated with high confidence.

One-third of the genera (13/39) have both L1 and L3/4 but no known, extant L2 viruses. Among genera containing L2 viruses, a L1-L2 transition was followed by a L2-L3/4 transition in just four virus lineages (from the genera *Phlebovirus*, *Orthonairovirus*, *Henipavirus* and *Orthohepevirus*). Overall, we estimate that direct evolution from L1 virus lineages is the most likely origin of 74% of the 57 L3/4 lineages, with just 17% from L2 virus lineages and 9% unknown (Figures 2, S3). Eight genera show species-level diversification within L3/4 lineages, potentially within human populations, but this accounts for only 21/70 (30%) of L3/4 RNA virus species in our data set (Figures 2, S3).

The relative frequencies of L1-L2, L1-L3/4 and L2-L3/4 transitions varied across genera, families and genome types (Figures 3b-d, Table S2). The odds of L1-L3/4 relative to L1-L2 transitions were lower for (-)ssRNA versus (+)ssRNA or dsRNA (Figure 2d), for enveloped versus non-enveloped (Figure 3e) and for vector-borne versus non-vector-borne viruses (Figure 3f) with unadjusted ORs=0.15, 0.23 and 0.23 respectively, all P<0.05. However, these traits are correlated and multivariable analysis suggested that the dominant effect was the low odds of L1-L3/4 transitions among (-)ssRNA viruses relative to dsRNA and (+)ssRNA viruses (Table 1). We note that L2-L3/4 transitions were highly over-represented (unadjusted OR=11.5, P<0.001) among vector-borne viruses (Figure 3f), occurring in only three non-vector-borne virus genera (*Henipavirus*, *Metapneumovirus* and *Orthohepevirus*).

**Table 1.**
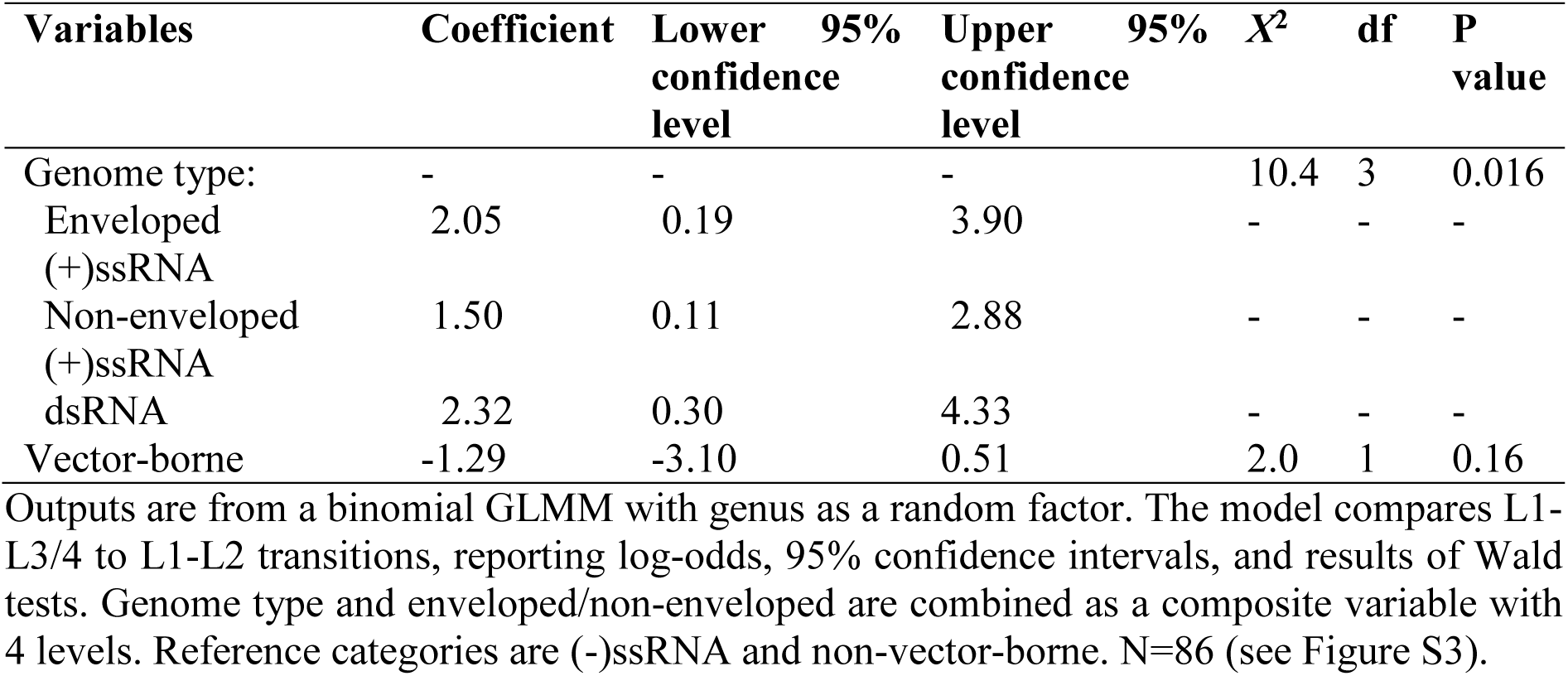
Ancestor node traits as predictors of IT level transitions.

L1-L2, L1-L3/4 and L2-L3/4 transitions had similar distributions of relative node depth (Table S3, Figure S4). There was no indication that L2 viruses tended to arise earlier in lineages than L3/4 viruses. Nor was there a consistent trend towards evolutionary recent (as measured by relative node depth) increases in the number of viruses entering human populations. We also estimated the instantaneous transition rates between IT level (see Supplementary Materials). The rates of L2-L3/4 transitions were not higher than the rates of L1-L3/4 transitions (Figure 3g, Table S4). This implies that the distributions of values of parameters *a* and *c* in our conceptual model (Figure 1) are similar. The estimated rates were highly variable across virus taxa (Figures S5, S6) but this was not associated with genome type/enveloped or being vector borne (Table S4).

In contrast to IT levels, host categories are not mutually exclusive traits of a virus species/subtype, and for some genera (e.g. *Ebolavirus*) it is likely that we are missing sequences from the reservoir host(s). Therefore, we did not attempt a discrete traits analysis for host range but compared the known non-human hosts for L2 and L3/4 viruses at the species/subtype and genus levels (see Supplementary Materials). At the species/subtype level human viruses also found in non-human primates were most likely to be L3/4 rather than L2; those also found in birds were least likely (Figure 4). At the genus level the pattern was similar (Spearman’s rank correlation of coefficient estimates by host category = +0.89, P<0.01) but the confidence intervals are wider due to the smaller sample sizes (Figure S7). These results are consistent with primates being disproportionately frequent ancestral hosts of L3/4 virus lineages, but there are too few virus sequences from wild primates available (74 in our data set) to test that hypothesis more formally through phylogenetic analysis.

**Figure 4.**
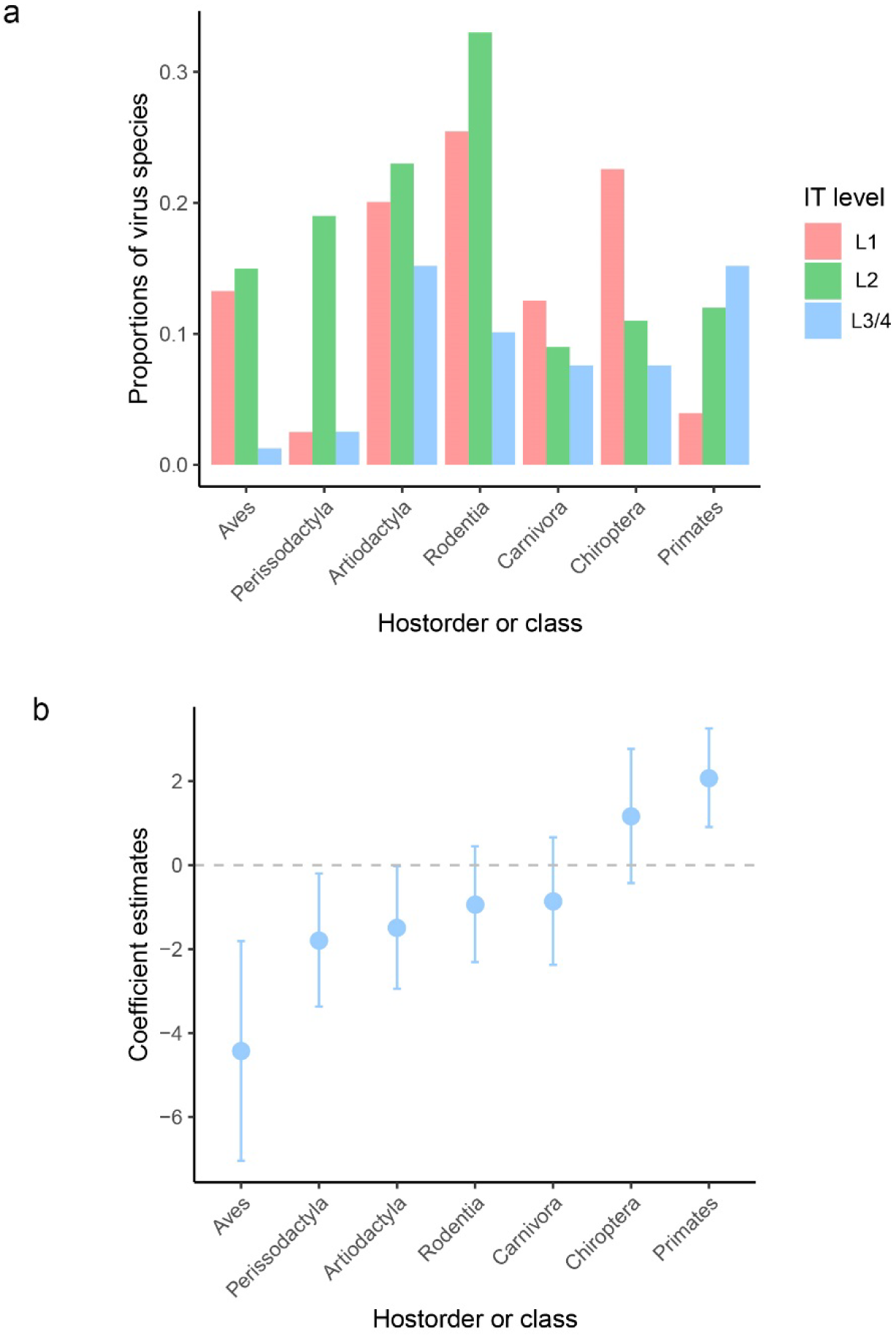
Human-transmissible virus in nonhuman host. a) The proportions of virus species of different IT levels being known to infect a given non-human host category (six orders of mammal and the Class Aves). b) Results for GLMMs with binary responses for association between viruses within a species/subtype being human-transmissible (L3/4) and being known to infect a given non-human host category. Coefficient estimates (with 95% CIs) are compared for six orders of mammal and the Class Aves of L3/4 virus species relative to L2 virus species. Coefficient estimates greater than zero correspond to a positive association. Models include a random term for each observation to account for over-dispersion. Genus and family are included as random effects.

In keeping with earlier work^15^, we used RNA-dependent RNA polymerase protein-based phylogenies in our main analysis. However, we tested the possibility that alternative phylogenies would generate different numbers and distributions of transitions between traits by constructing alternatives trees based on surface protein sequences (Figure S8). We obtained results that differed minimally from our main analysis, most importantly that an estimated 71% of L3/4 lineages have L1 ancestors (see Data File 9).

We recognise that there will be significant gaps in our knowledge of mammal/bird non-RT RNA virus diversity: new species are routinely being identified^8^ and virus genome sequences are accumulating rapidly (Figure S1). Our selection procedure resulted in a data set with only 28% of virus sequences from human hosts, but it is still likely that the phylogenetic diversity of viruses from non-human hosts is greatly underestimated relative to that from humans^3, 14^, noting that for many mammal/bird taxa no RNA viruses at all have yet been reported. There will also be gaps in our knowledge of IT level: some viruses originally classified as L1 have been subsequently found to human infective; and some viruses originally classified as L2 have been subsequently found to be human transmissible as epidemiological data accumulates^6^. As new virus sequences are added and IT levels are assigned or (occasionally) re-assigned it is entirely possible that the estimated ancestors of some L3/4 viruses will change. However, we anticipate that the impact of such changes across the whole data set will be to strengthen our main conclusions. This is because we anticipate that far more L1 viruses will be added than L2, and more L2 than L3/4, continuing the current trend (Figure S1). For this reason, we consider our estimate that 74% of L3/4 viruses arose from L1 lineages to be conservative.

We did not include two important taxa in our main analysis. The *Influenza A virus* genus was excluded *a posteriori* because of a very weak phylogenetic signal for IT level (Figure S9). On inspection of the HA gene phylogeny it is apparent that although human infectivity and transmissibility have evolved in only a few lineages (HA types) there is considerable variability in IT level within those lineages, which would generate inflated counts of IT level change (Figure S10). Nonetheless, the phylogeny is broadly consistent with the pattern that most L3/4 lineages evolve from L1 lineages. We excluded Retroviridae from our main analysis *a priori* because they cause chronic infections that allow far more time for within-host evolution prior to transmission^9^. Consistent with this, we find that transmissible human lentiviruses (lineages of HIV-1 and HIV-2) are most likely have evolved through L1-L2-L3/4 transitions rather than L1-L3/4 (Figure S11).

Our results show that at least 74% of L3/4 non-RT RNA virus lineages (e.g. human coronaviruses 229E and NL63, measles virus, human respiratory syncytial virus, hepatitis C virus and Aichi virus A) are most likely to have emerged directly from reservoirs of L1 viruses. Stepwise emergence to L2 then L3/4 occurs infrequently and mainly in vector-borne viruses. This result is consistent with epidemiological observation: there are no well-supported examples of any of the >100 species of L2 non-RT RNA viruses evolving the capacity to transmit in human populations^6^. We also estimate that diversification within L3/4 lineages has contributed significantly to human RNA virus diversity, accounting for 30% of species (Figure 1). However, a larger fraction, at least 51%, has been generated directly from L1 lineages.

The evolution of a new (and extant) lineage of L3/4 non-RT RNA viruses appears a relatively infrequent event; there are just 57 instances over the entire evolutionary history of the genera considered here, with no suggestion of any recent (on an evolutionary time scale) increase in frequency (Figure S4). Unexpectedly, the same applies to L2 viruses; these are generated from L1 viruses at similar rates as L3/4 viruses and at similar relative node depths (Figures 3g and S4, Table S3). We have previously suggested an alternative model where the L2 trait is easily evolved and easily lost^3^, but this model is not supported by the analysis reported here. Importantly, these findings do not preclude extant pools of L2 and L3/4 viruses that have yet to be recognised and may not yet have had the opportunity to enter human populations^16^, but they do imply that the drivers for the emergence of novel human viruses, with or without epidemic potential, would be ecological rather than evolutionary.

That there was no detectable difference in the relative rates at which L3/4 viruses emerge from L2 and non-human (L1) virus lineages (Figure 3g) implies that the distributions of parameters *a* and *c* in our conceptual model (Figure 1) are similar. Given that, an obvious explanation for the rarity of L2-L3/4 transitions is that the pool of L1 viruses is much larger than that of L2 viruses, as is widely supposed^10, 12^. Moreover, most L2-L3/4 transitions involve vector-borne viruses. Vector-borne pathogens tend to have wide host ranges^17^, possibly due to the transmission route providing direct access to the blood system. There is a substantial deficit of vector-borne L1 viruses in our database (Data File 4), consistent with the L2 pool being relatively larger for vector-borne viruses, thus explaining why we found most L2-L3/4 transitions in this category. In addition, we found that L1 (-)ssRNA virus lineages are relatively more likely to generate L2 than L3/4 viruses (Table 1) and, once this was accounted for, there was no additional effect of a virus having an envelope, as has been suggested previously^5^. Nonetheless, this is a large group and (-)ssRNA virus lineages are an important source of L3/4 viruses (Figure 3d).

The absence of a clear association between human infectivity and human transmissibility may reflect the key role that cell receptors play in determining a virus’s capacity to infect and be transmitted by humans^16, 18^. Cell receptor usage varies between virus genera and sometimes within genera^9^. Host switching (to humans, or to any other new host) is facilitated by a virus using a cell receptor with an amino acid sequence that is conserved between hosts^16^. However, if the receptor has a different tissue distribution in humans then infectivity may not equate with transmissibility^18^. Evolutionary shifts in receptor usage can therefore lead to changes in human infectivity (L1 to L2) or transmissibility (L2 to L3/4) or both (L1 to L3/4).

Our study supports a recent proposal for a large-scale survey of viral diversity in non-human reservoirs, the Global Virome Project (GVP)^11^. Though this idea is controversial and would be costly to implement^19^, its relevance is underlined by our finding that most species of human-transmissible viruses (L3/4) evolve from mammal/bird RNA virus lineages not known to be infective to humans (L1), coupled with the expectation that the great majority of mammal and bird viruses are still unrecognised^12^. We note recent progress in using machine learning to predict host range from sequence data^20^ and the potential of this kind of approach to help identify human-infective and human-transmissible viruses even in the absence of human cases.

Finally, our study also provides empirical support for, and underlines the public health importance of, “Disease X”, the scenario that a future serious international epidemic might be caused by a pathogen taxon not currently known to affect humans^21^. We find that the ability of RNA viruses to transmit between humans most frequently evolves in a manner consistent with the “Disease X” model of emerging infections.

## ACKNOWLEDGEMENTS

We are grateful to Andrew Rambaut and Paul Sharp for helpful discussions, and to Alex Bhattacharya for assistance with data preparation.

## Author contributions

L.L., S.L., P.S. and M.W. designed the study. L.L., L.B. and F.Z. assembled the data sets. L.L., G.R. and M.C-T performed the analyses. D.S. validated some of the phylogenies. L.L. and M.W. wrote the manuscript. All authors reviewed, commented on and approved the manuscript.

## Data availability

The raw data used in these analyses are all freely available in the Data Files. These include links to sequence data in GenBank.

## SUPPLEMENTARY MATERIALS

### Materials and Methods

#### Data

We compiled a data pool of 7488 polymerase (or functional equivalent) gene sequences from non-RT RNA viruses found naturally (i.e. excluding deliberate laboratory exposures) in humans, other mammals or birds and reported in the NCBI GenBank database^1^. Inclusion criteria were that complete or nearly complete polymerase protein sequences were available, together with information of natural infected host (mammal or bird only) species and isolation date (year). No sequence data were available for 30 otherwise eligible human-infective virus species and these could not be included in the analysis – all of these were rare viruses, the majority of which have not been reported in humans for at least 10 years. For some zoonotic virus species, sequences were only available from non-human hosts. To reduce sampling bias, especially of human viruses, we then subsampled the dataset to keep no more than five sequences per host species per virus species/subtype and to avoid clusters of sequences from single outbreaks.

We linked individual sequences to infectivity/transmissibility (IT) level: L1, L2 or L3/4 as defined in the main text. IT level was attributed using an updated version of a recently published epidemiological data set but six viruses that are transmitted between humans only by vertical and/or iatrogenic routes^2^ were categorised here as L2. Four virus species (*Norovirus, Sapovirus, Orthohepevirus A, Aichivirus A*) for which there are recognised subtypes that differ in IT level were included as subtypes rather than single species.

Seven virus genera (*Coltivirus*, *Erbovirus*, *Marburgvirus*, *Seadornavirus*, *Thogotovirus*, *Tibrovirus* and *Torovirus*) were excluded because there were too few sequences (<15) for analysis. Three genera (*Deltavirus*, *Rubivirus* and *Salivirus*) provided no useful information because they each contain a single L3/4 species known only from humans. Two genera (*Influenza B virus* and *Influenza C virus*) had too few non-L3/4 sequences available (one and three respectively). The *Influenza A virus* genus was considered separately from the main analysis as BaTS tests indicated that there is a uniquely poor association between IT level and polymerase phylogeny for this taxon (see below). Retroviridae (three genera with human-infective species) were excluded *a priori* from our main analysis given their markedly different biology^3^. However, we carried out a separate analysis for the one genus – *Lentivirus* – with sufficient *pol* protein sequences available.

Our main data set comprised 1755 sequences from 39 non-RT RNA virus genera (31 with human-transmissible (L3/4) viruses) for analysis. We linked these sequences to the host category from which the virus was isolated, as nine taxonomic orders of the Class Mammalia (distinguishing humans and non-human primates), plus marsupials and collectively the Class Aves (noting that <5 sequences were available from Eulipotyphla, Lagomorpha, Scandentia and marsupials, with the remaining mammalian orders not represented at all). One multi-species genus, *Cosavirus*, was represented only by sequences from human hosts. Links to the sequences and associated metadata are provided in Data File 4.

#### Phylogenetic analysis

We performed parallel phylogenetic analyses using the BEAST software package^4^ (V1.8.2). We were primarily interested in tree topology; branch lengths scale to numbers of amino acid substitutions and we did not attempt to date the nodes – such estimates are likely to be unreliable given the long time scales involved^5^. For every virus genus, polymerase gene sequences were translated to amino acid and aligned with MUSCLE V3.8.4^6^, and we reconstructed Bayesian time-scaled phylogenies for each of the 39 genera and perform discrete traits analysis with respect to IT level. Our methodology is comparable to the approach used in Ref.^7^.

We tested the associations between IT levels and polymerase trees per genus using Phylogeny-trait association test (BaTS)^8^, inspecting both the Association index (AI) and Parsimony score (PS) between genera. We observed that the *Influenza A virus* genus showed evidence of a uniquely weak phylogenetic signal (Fig. S9a) so this genus was subsequently considered separately from the main analysis. Similar results were obtained using phylogenies based on surface proteins as described below (Fig. S9b).

We estimated the phylogenies of the sequences using a Bayesian Markov chain Monte Carlo (MCMC) method that was implemented using the Bayesian evolutionary analysis in BEAST. Different combinations of substitution models, clock models and population size models were evaluated by using the path sampling (PS) to estimate marginal likelihoods^9^. The best fitting model was a WAG model with a gamma distribution (G) across sites as the substitution model, with an uncorrelated log-normal relaxed molecular clock model and with a constant size coalescent process or Yule process prior over the phylogenies. Here, we allowed the branch length to be scaled by substitution per site rather than by time (with ucld.mean equal to 1). The MCMC chains were run for 100 million iterations with sub-sampling every 10,000 iterations and 10% burn-in and at least two replicates were performed per genus. MCMC convergence and effective sample size of parameter estimates were evaluated using Tracer 1.5 (http://beast.bio.ed.ac.uk). Summary Maximum Clade Credibility phylogenies were created from the posterior samples of trees using TreeAnnotator, and a further sample of 1000 trees was extracted from each genus-level posterior sample for additional processing. We validated our phylogenies of the mammal/bird virus clade by comparing their topologies with the representative phylogenies on the family/genus level published by ICTV^3^, noting that we are missing sequences/species that did not meet our inclusion criteria.

We applied asymmetric discrete trait models using BEAST to estimate IT level over each genus-level posterior sample of 1000 trees^10^. The instantaneous transition probabilities between IT levels (equivalent to the relative rates matrix) were estimated simultaneously. All the Bayesian phylogenies mapped by IT level traits are provided in Data File 5.

For each genus-level phylogeny we obtained the estimated the probability distributions for IT level at the root ancestral node. Transitions were identified as a change in the most probable IT level estimated for any two adjacent nodes on the phylogeny (Fig. 2). We counted all occurrences within each tree of the three possible types of ‘forward’ transition: L1 to L2 (non-human virus acquiring human infectivity); L1 to L3/4 (non-human virus acquiring human infectivity and transmissibility); and L2 to L3/4 (infective but non-transmissible virus acquiring human transmissibility). Instances of the possible ‘backward’ transitions (L2 to L1, L3/4 to L1 and L3/4 to L2) were also recorded.

We also applied Markov Jumps ^11^ with discrete trait model to estimate and validate the expected number of level transitions in the phylogenetic trees (an example XML file shown in Data File 6). We obtained excellent agreement between median estimates of numbers of forward transitions using Markov jumps and counts estimated from our discrete traits analysis (Table S1, Fig. S12a). We were also able to reproduce the patterns of variation between virus types and taxa with minimal discrepancies (compare Figs 2b-f and S13a-e). Outputs are compared in full in Data File 7.

In addition, we applied parsimony reconstruction models to trace trait evolution, using Mesquite V 3.5.1 (http://www.mesquiteproject.org), with the input phylogenies being generated with the protein sequences of each virus genus via maximum likelihood method, using RaxML V 8 (WAG+G, bootstraps n=1000)^12^. We obtained excellent agreement between mean estimates of numbers of forward transitions using parsimony analysis and counts estimated from our discrete traits analysis (Table S1, Fig. S12b), noting that the parsimony approach does not identify individual nodes involved in transitions. Outputs are compared in full in Data File 7.

To test the robustness of our results to the choice of RNA-dependent RNA polymerase protein sequences to construct our phylogenies we compared results using different proteins. We used 1560 surface protein amino acid sequences (Data File 8) to generate alternative phylogenies for 35 genera (Fig. S8), by mapping the correspondent surface protein with polymerase protein from the same virus strain. We found only a small number of discrepancies between numbers of L1-L2, L1-L3/4 and L2-L3/4 transitions for the two sets of phylogenies (Fig. S14), with total numbers of the three transition types differing by -5, -1 and +1 respectively (Data File 9).

#### Statistical analysis

We used Fisher’s exact tests to examine whether relative frequencies of transitions between different IT levels varied among families, genera, genome types, and between vector-borne/non-vector-borne viruses. We then compared frequencies of L1-L2 and L1-L3/4 transitions using a binomial generalised linear mixed model (GLMM) with transition type as the response variable, genome/enveloped type and vector-borne/non-vector-borne as explanatory variables and genus as a random factor (using the R package ‘lme4’). We constructed a composite variable representing the genome type and enveloped structure of viruses involved in transition events. This had four levels: (-)ssRNA (all enveloped), (+)ssRNA enveloped, (+)ssRNA non-enveloped, and dsRNA (all non-enveloped). Models with genus as the only random factor and models with both genus and family as random factors had similar AIC values (ΔAIC<2), so the former were used in all analyses.

We calculated the depth of the node of each transition on the genus-level phylogenetic tree relative to the ancestral node (Fig. S4 and Data Files 2 and 3). We used a beta distributed GLMM (using the R package ‘glmmTMB’^13^) to compare node depth (response variable) across level transition types (explanatory variable) with genus as a random factor. We used a linear mixed model (LMM) with a normal distribution to compare log_e_-transformed transition rates (response variable) by level transition groups (explanatory variable) with genus as a random factor. We also included genome type/enveloped and vector-borne in the model as potential confounding variables.

We tested for an association between host range and IT level in two ways. First, we considered known non-human hosts of the 179 human virus species/subtypes in our data set, using availability of virus sequence as a gold standard indicator of host range. Second, we considered known non-human hosts for all viruses from each of 39 genera using the same criteria. For these analyses, we classified species/subtypes and genera as non-human-transmissible if they occurred in a genus or family respectively that contained only viruses incapable of epidemic spread in human populations (i.e. L2 or L3). To compare host range distributions at species/subtype and genus levels we used GLMMs with binary responses to generate odds ratios for containing human-transmissible viruses, with 95% confidence limits, by host category. We included genus and family plus genus respectively as random effects to allow for taxonomic relatedness.

---

**Fig. S1.**
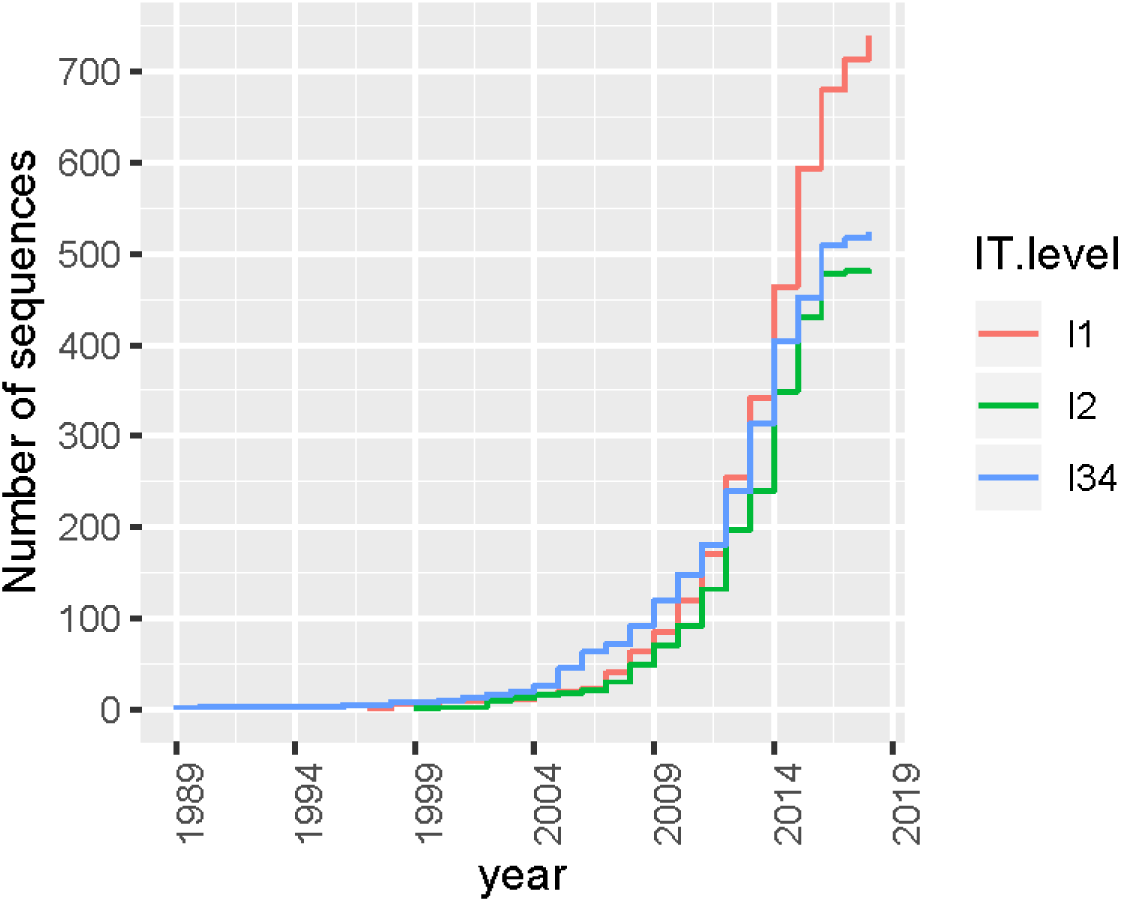
Numbers of sequences used in this study cumulated by year of submission to Genbank. L1, L2 and L3/4 viruses are distinguished.

**Fig. S2.**
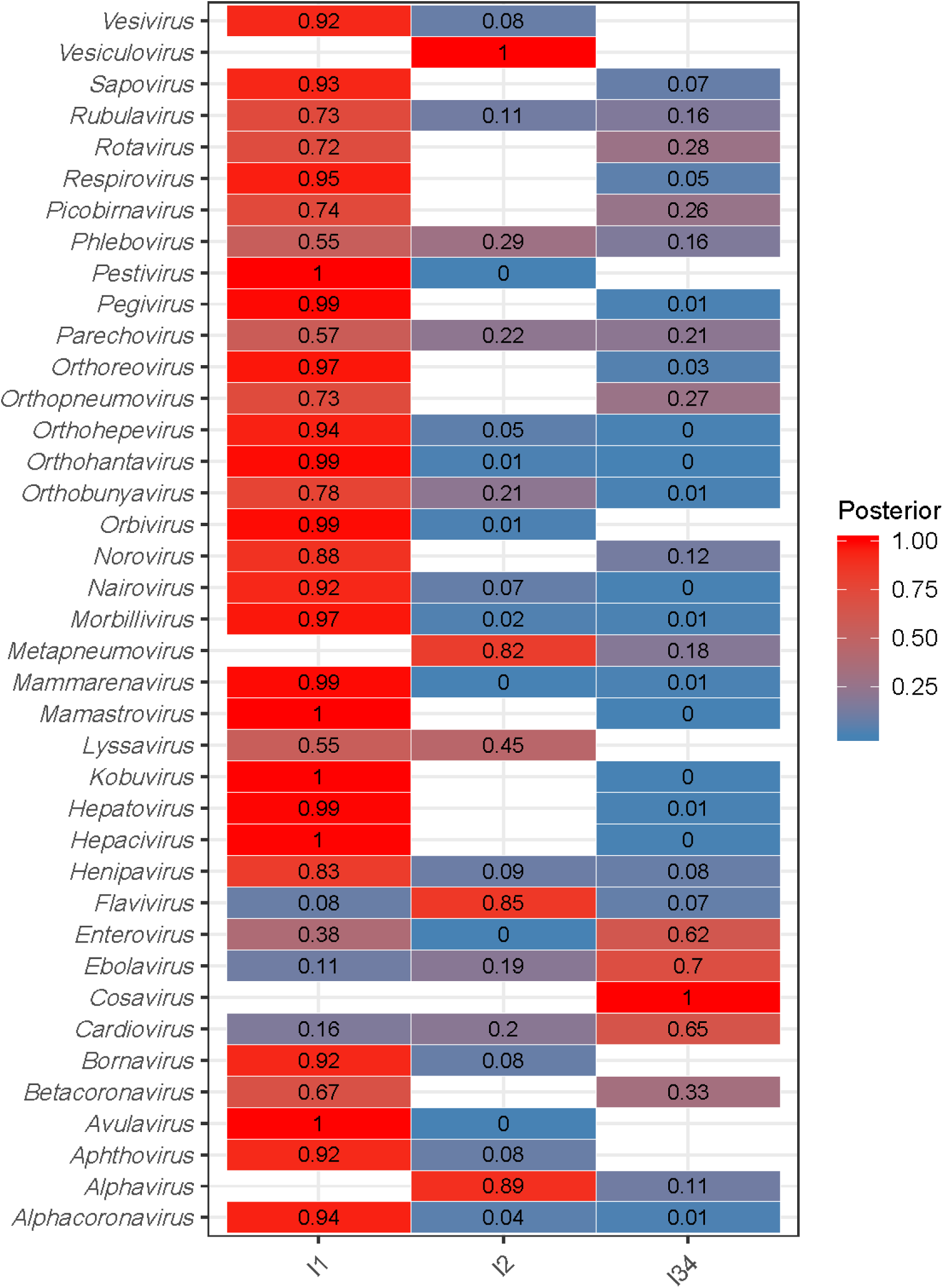
Heat map showing the estimated probabilities that the ancestor for each genus (N=39) was a L1, L2 or L3/4 virus. Blank entries indicate that no sequences from viruses at this IT level were available.

**Fig. S3.**
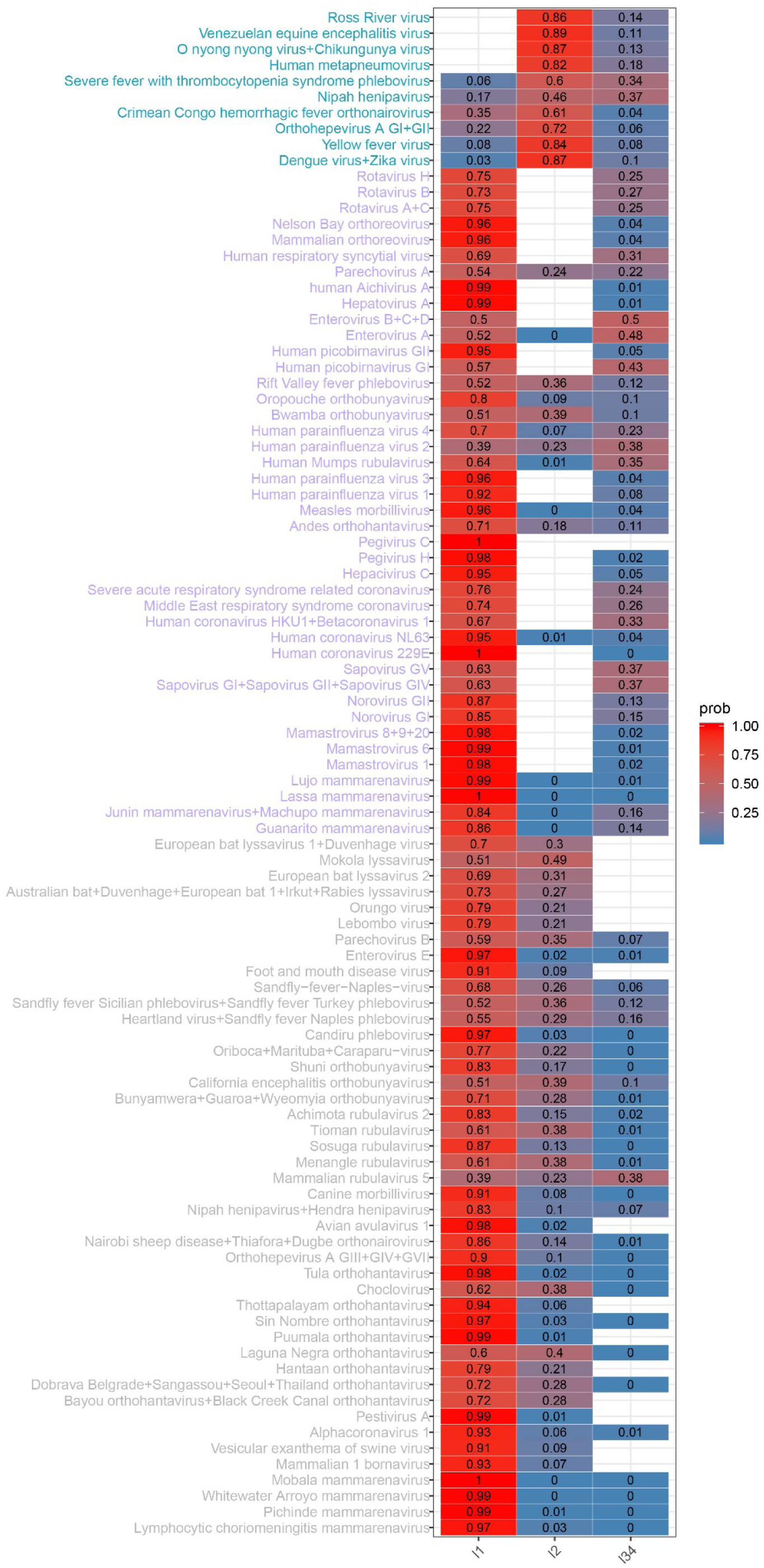
Heat map showing the total estimated probabilities that the ancestral node for each forward transition (N=96) was a L1, L2 or L3/4 virus. Three transitions are distinguished: L1 to L2 (greylabels), L1 to L3/4 (purple) and L2 to L3/4 (cyan). Blank entries indicate that no sequences from viruses at this level were present in the sequence database.

**Fig. S4.**
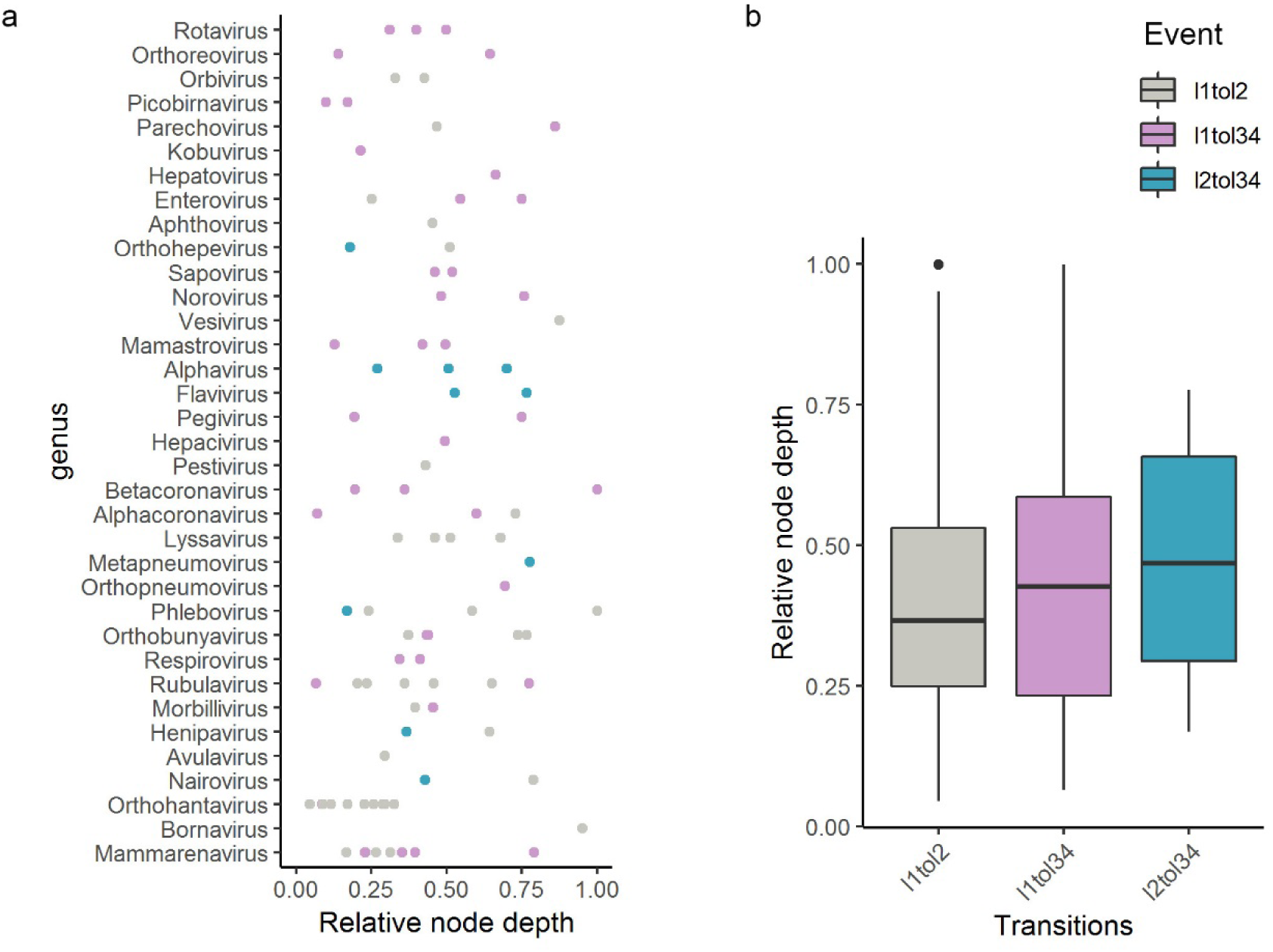
Relative node depths of transitions by genus. Three transitions are distinguished: L1 to L2 (grey), L1 to L3/4 (purple) and L2 to L3/4 (cyan), shown in a) dotplot per genus, b) boxplot per transition. Relative node depths are given in Data File 2.

**Fig. S5.**
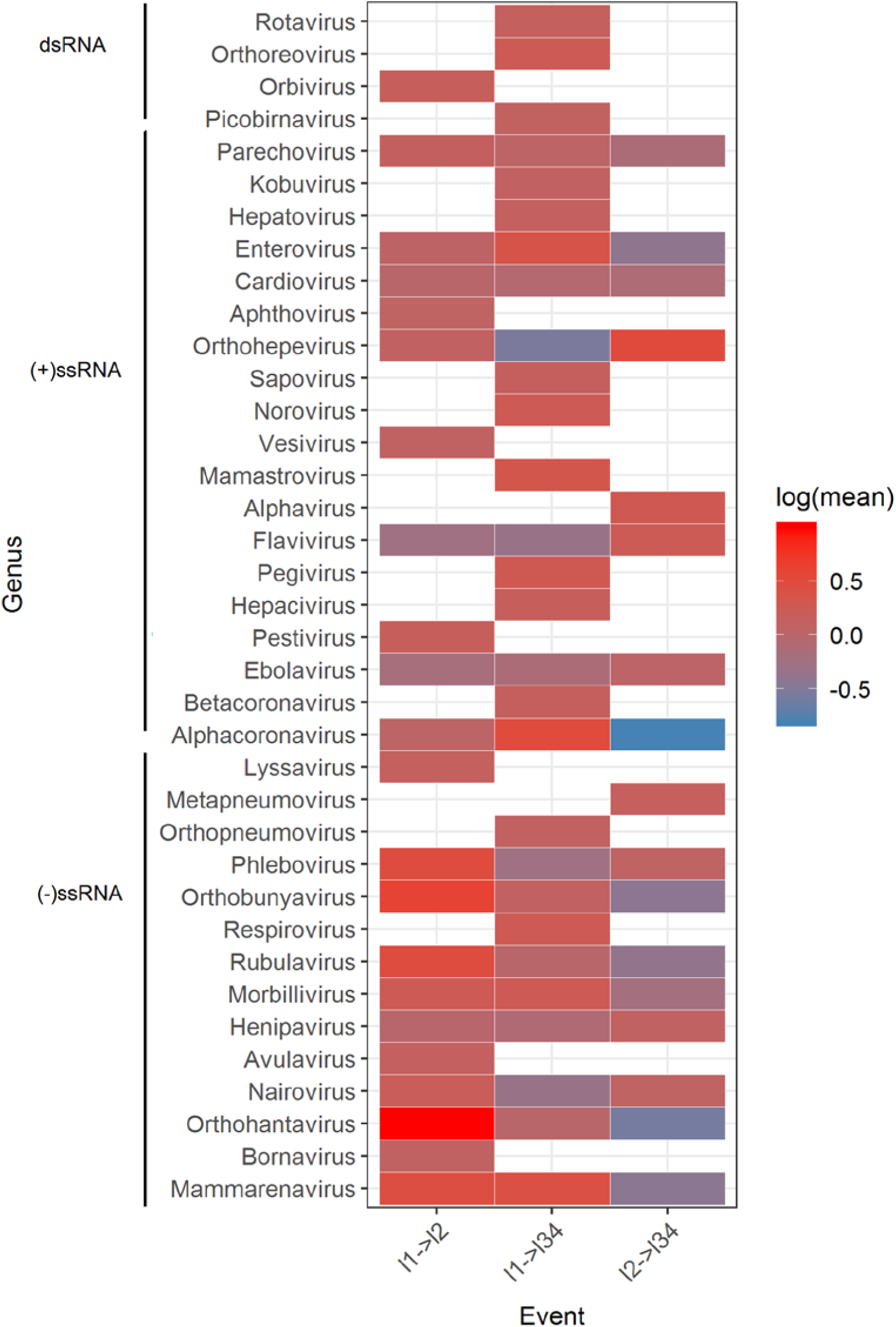
Comparison of instantaneous rates of L1-L2, L1-L3/4 and L2-L3/4 transitions for each genus, using mean relative transition rates on a log scale (see Fig. S6). Blank entries indicate that a transition was not possible in this genus given the information available.

**Fig. S6.**
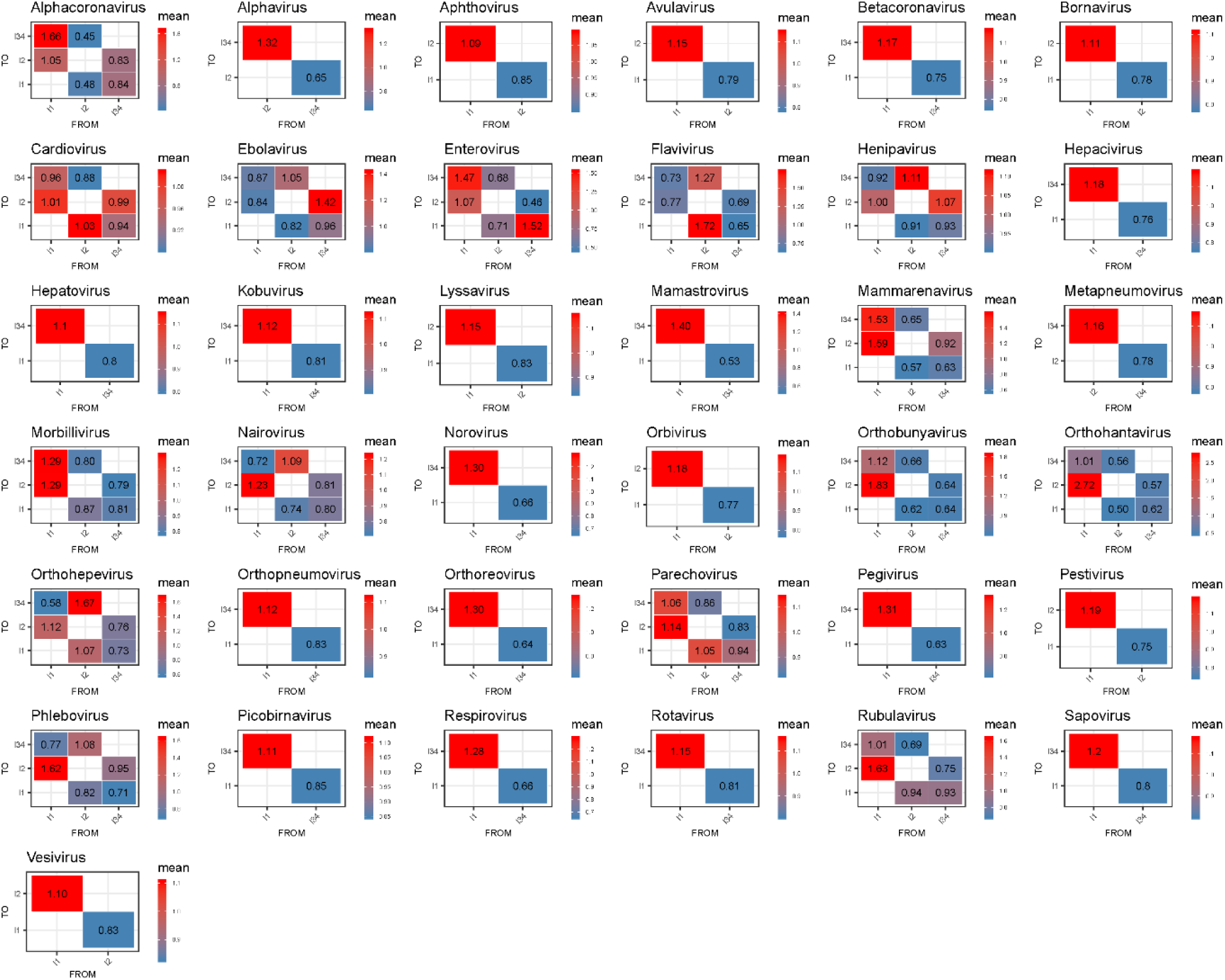
Transmission matrices of relative transition rates between IT levels in each genus with ≥1 transition (N=37). The mean estimates of instantaneous transition rates are shown.

**Fig. S7.**
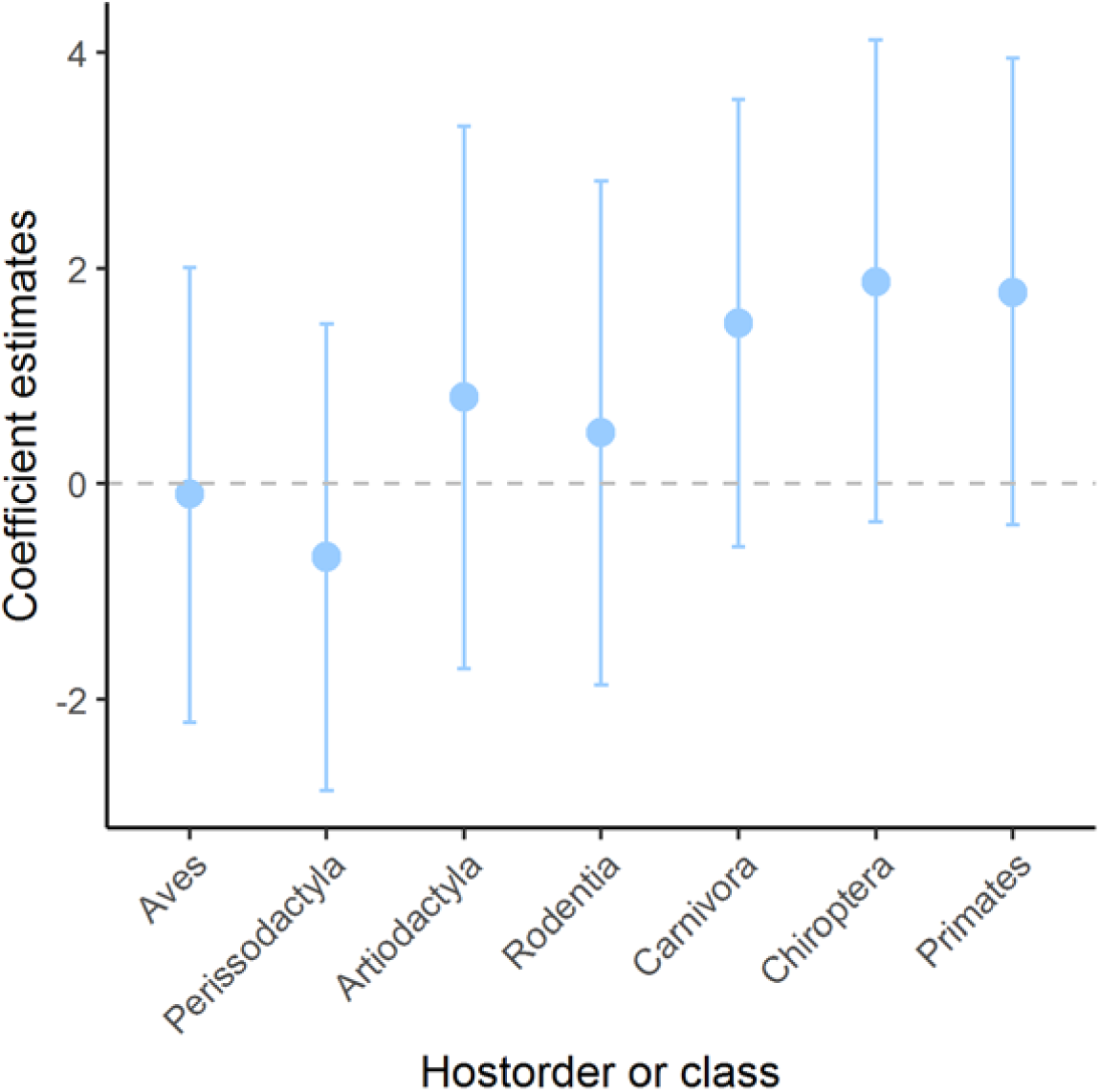
Results for GLMMs with binary responses of the association between viruses within a genus being human-transmissible (L3/4) and being known to infect a given non-human host category. Coefficient estimates (with 95% CIs) are compared for six orders of mammal and the Class Aves. Coefficient estimates greater than zero correspond to a positive association. Models include a random term for each observation to account for over-dispersion. Family is included as a random effect. See also Fig. 3b.

**Fig. S8.**
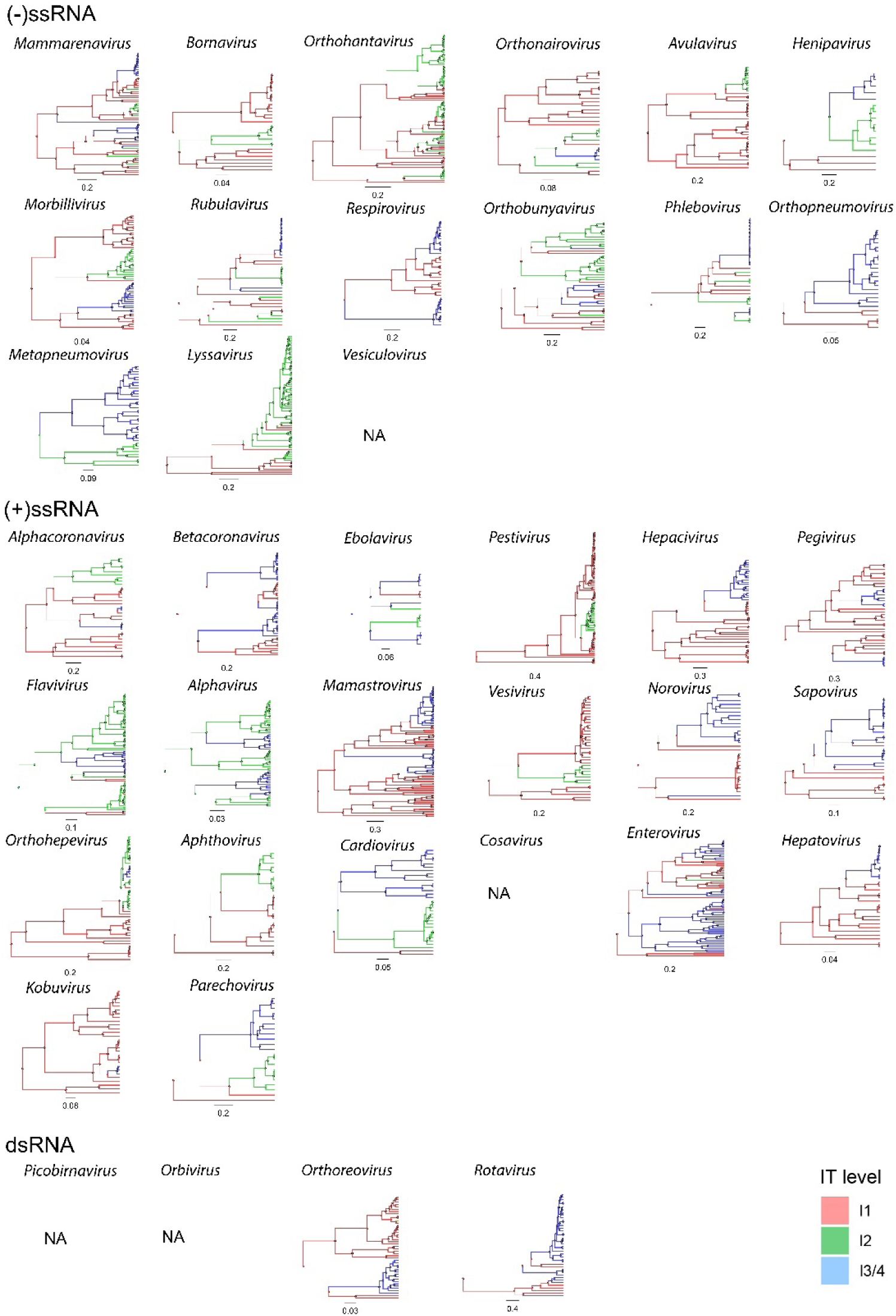
Bayesian maximum clade credibility (MCC) trees for members of 35 virus genera using surface protein sequences (listed in Data File 8). Phylogenies show the most probable transitions between non-human viruses (red), viruses infective to humans (green) and viruses transmissible in human populations (blue). Phylogenies are arranged by genome type.

**Fig. S9.**
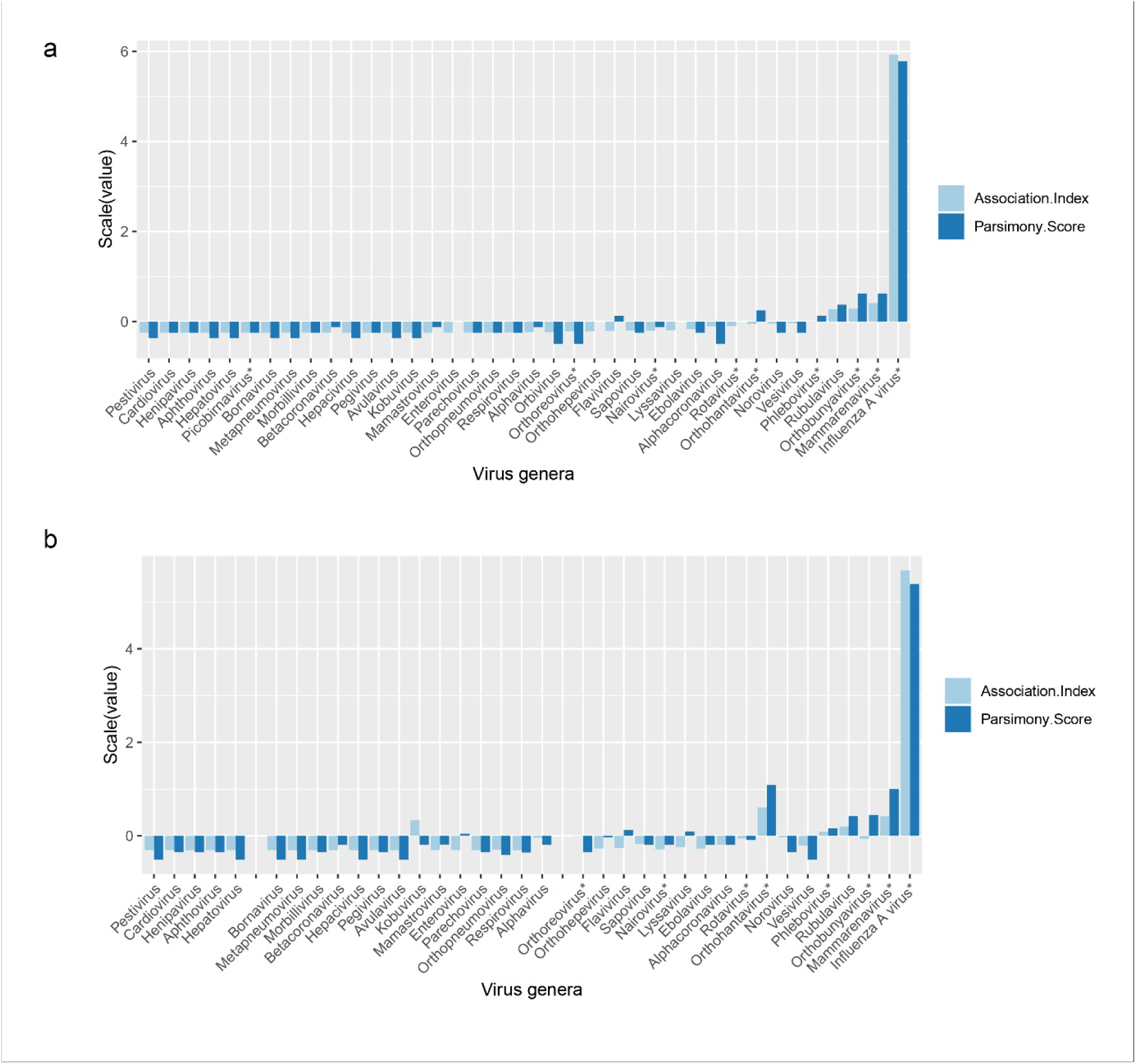
Outputs of Phylogeny-trait association (BaTS)analysis as estimated IT level trait and phylogenies associations per genus. Scaled (i.e. normalised as (x - mean)/standard deviation) values of Association Index (AI) in light blue; scaled Parsimony Score (PS) in dark blue. Higher AI and PS indicate lower association between trait and phylogeny. **a**) RNA-dependent RNA polymerase proteins; and **b**) surface protein proteins. N= 38 and 36 respectively (see Data File 8). Genera with segmented genomes are indicated with asterisk.

**Fig. S10.**
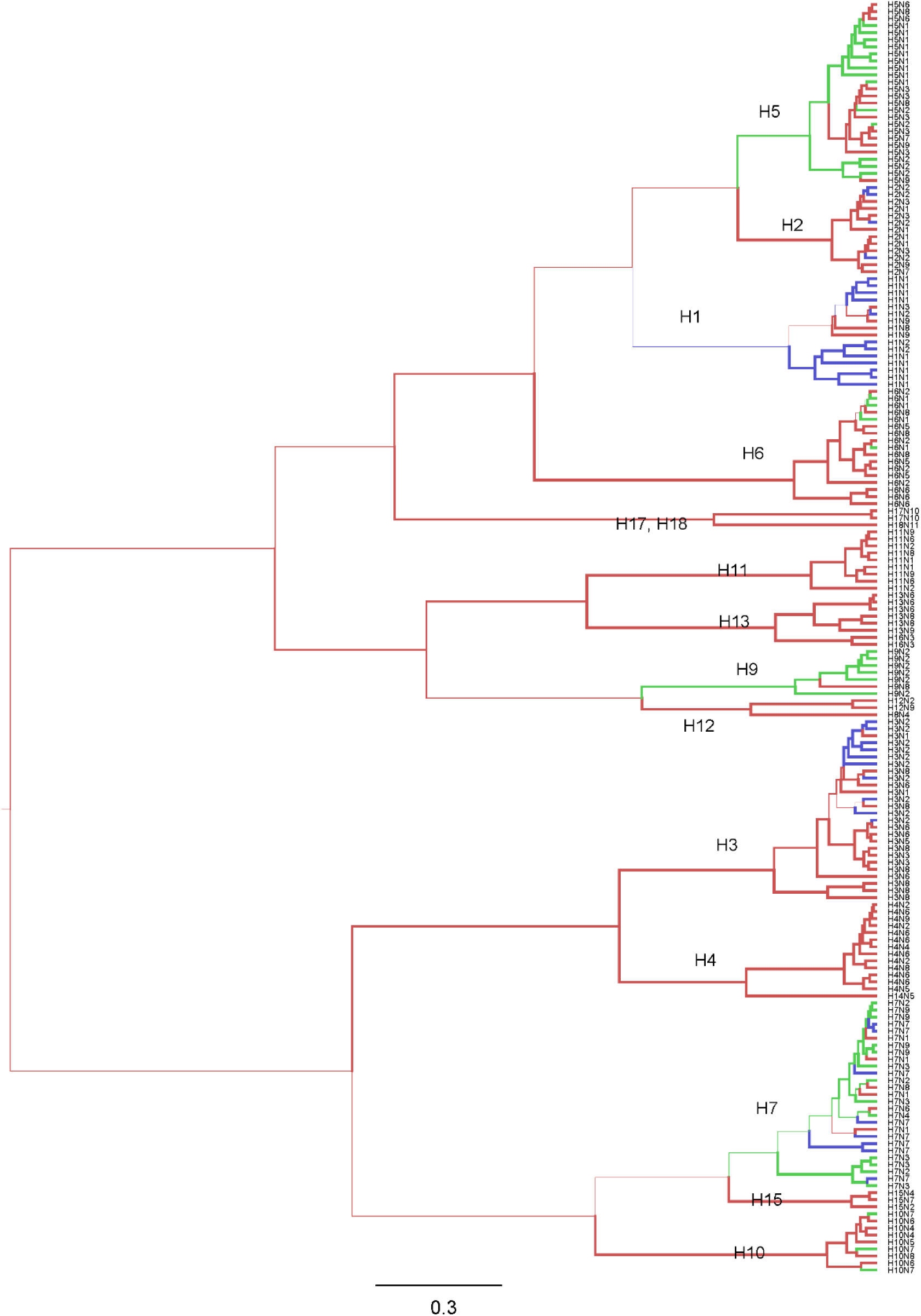
Bayesian maximum clade credibility (MCC) trees for *Influenza A virus* genus. Phylogeny generated using full length hemagglutinin (HA) protein sequences (N=181, representing 67 subtypes), mapped with IT levels using discrete trait models with Markov jumps. Nine subtypes (H10N7, H5N1, H5N2, H6N1, H7N2, H7N3, H7N4, H7N9, H9N2) are IT level 2; 5 subtypes (H7N7, H1N1, H1N2, H2N2, H3N2) are L3/4; other subtypes are L1. Colour coding as Fig. 1. The scale bar represents amino acid substitutions per site.

**Fig. S11.**
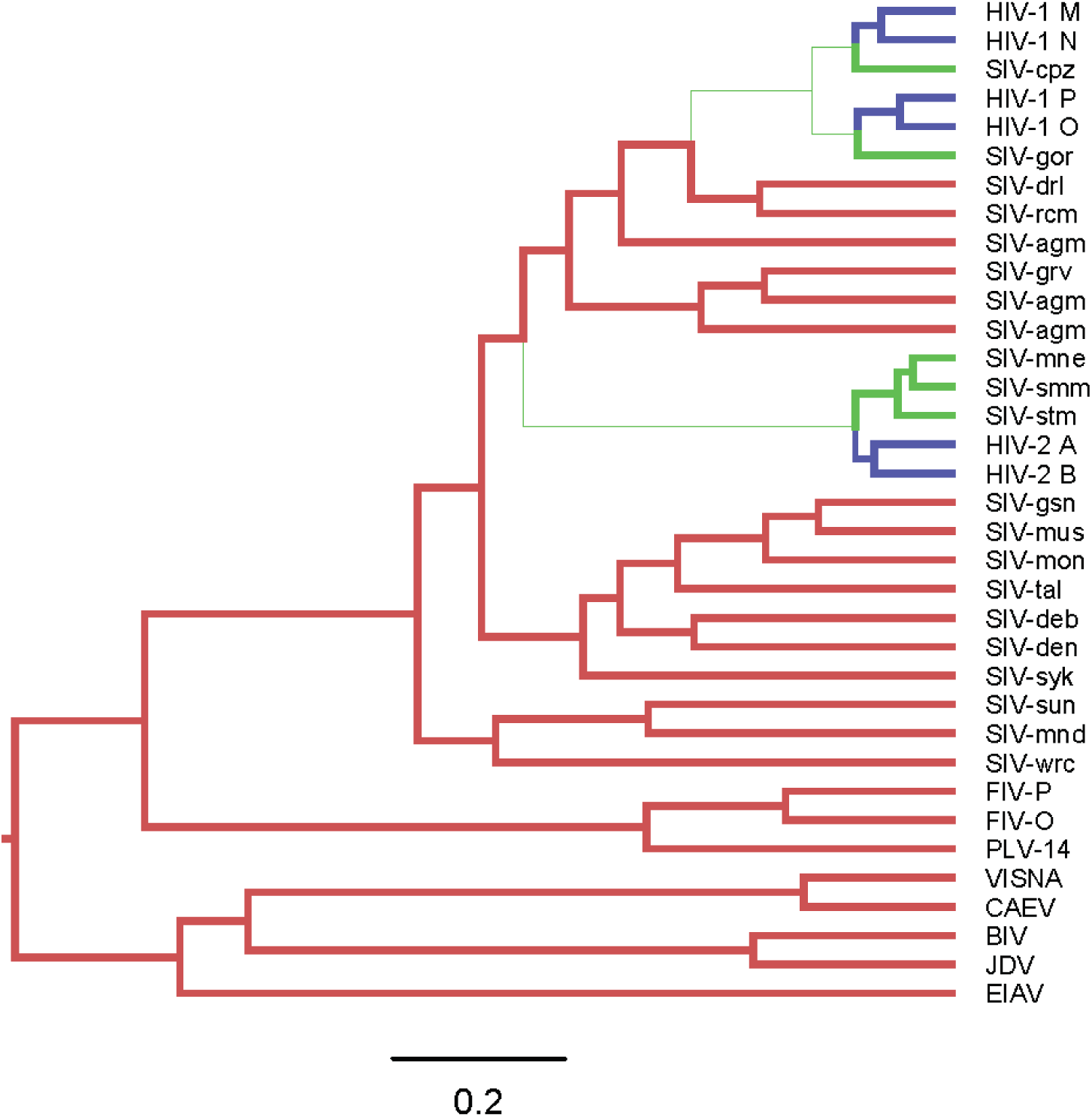
Bayesian maximum clade credibility (MCC) trees for the *Lentivirus* genus. Phylogeny generated using *pol* protein sequences (N=35, representing 10 species), mapped with IT levels using discrete trait models with Markov jumps. HIV-1 (group O, P, N, M) and HIV-2 (group A and B) are IT level 3/4; SIV (in chimpanzees, gorillas and sooty mangabeys) are L2, other species are L1 ^14^. Colour coding as Fig. 1. The scale bar represents amino acid substitutions per site.

**Fig. S12.**
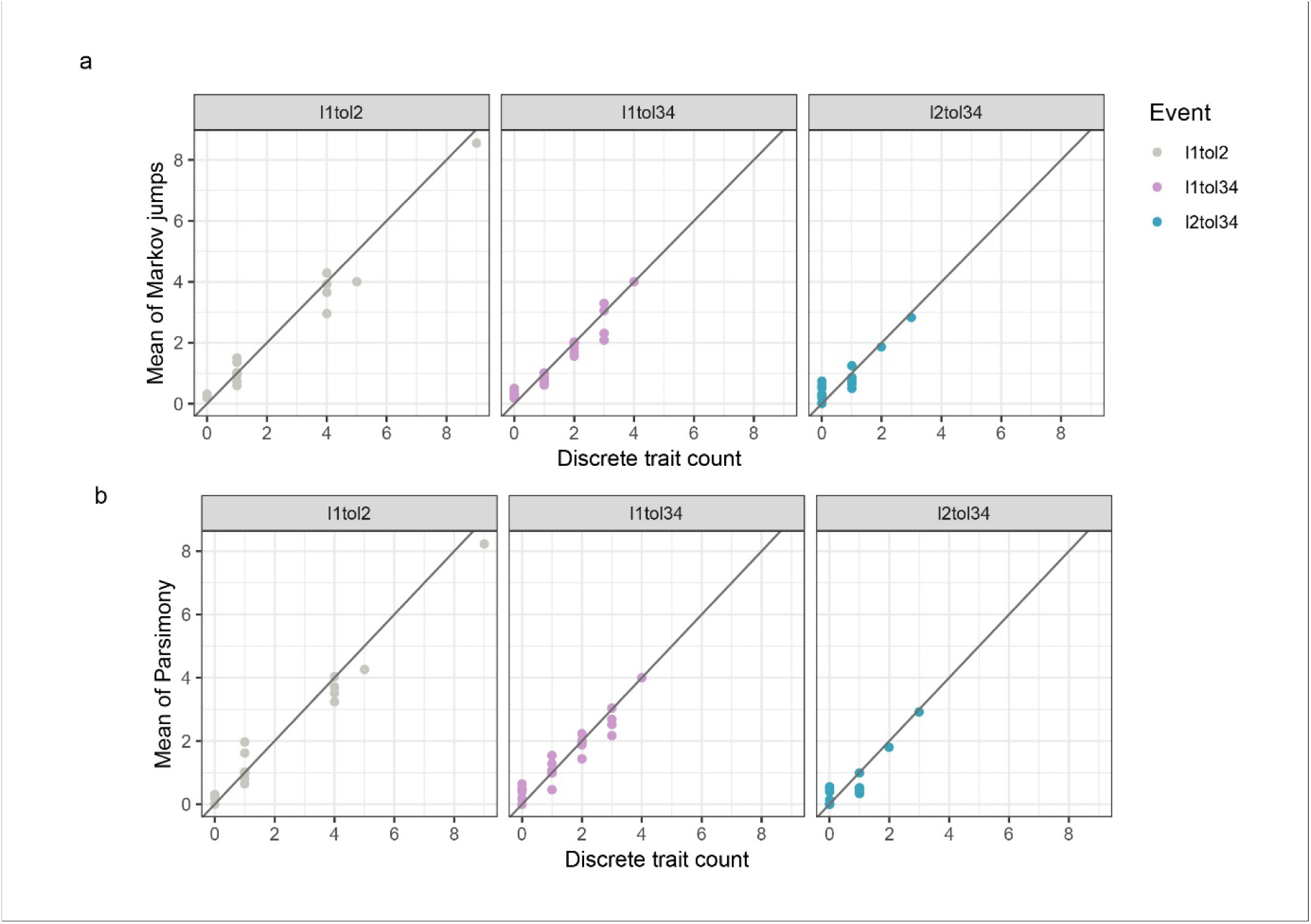
Comparison of number of transitions (mean) in different genera (N=37) using different methods (see Table S1). Comparing **a**) discrete trait count versus number of transitions (mean) estimated by Markov jumps and **b**) observed node changes on discrete trait phylogenies (discrete trait count) versus number of transitions (mean) estimated by parsimony. Lines of equivalence are shown.

**Fig. S13.**
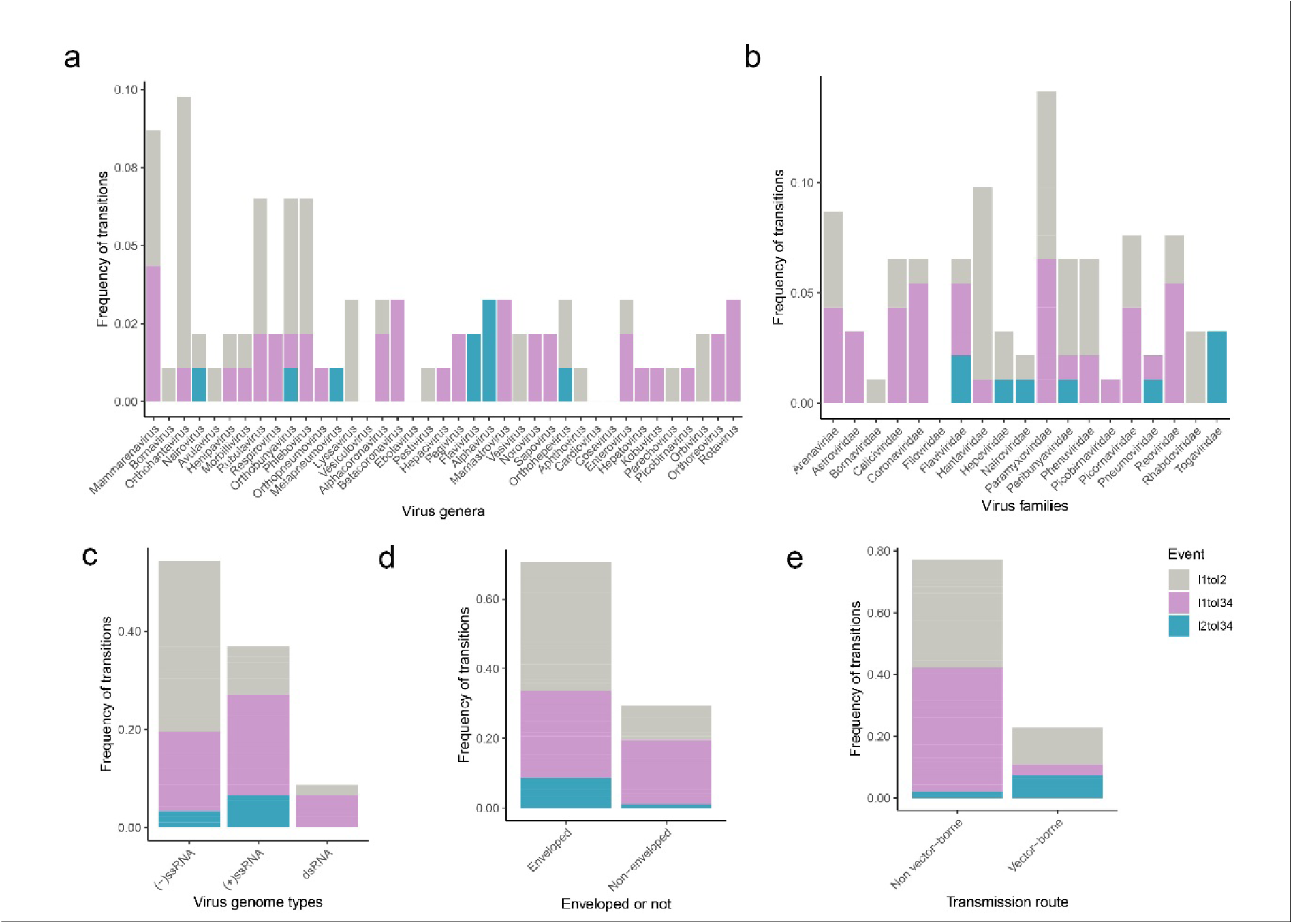
As Fig. 2b-f respectively but based on the results of parsimony analysis.

**Fig. S14.**
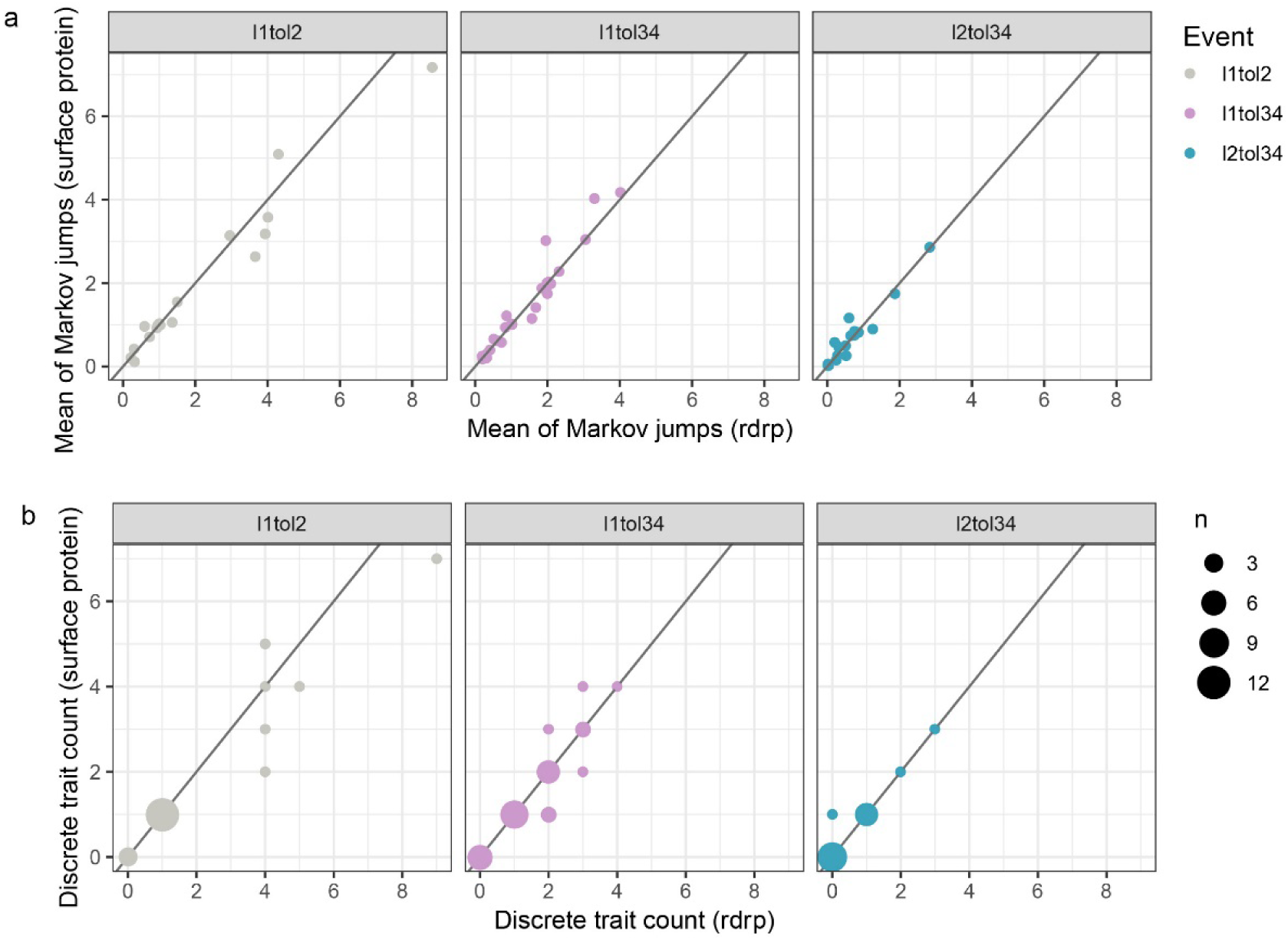
Comparison of number of transitions in different genera (N=35) estimated using phylogenies based on polymerase (rdrp) vs surface protein genes. Graphs compare: **a)** number of transitions (mean) estimated by Markov jumps and **b**) observed node changes on discrete trait phylogenies (discrete trait count). Lines of equivalence are shown.

**Table S1.** Numbers of estimated transitions between human-infective/transmissible (IT) levels 1, 2 and 3/4 across all genera (N=39) compared for three methodologies: 1) counts of the number of internal node changes, transitions, observed from discrete traits analysis* and 2) number of expected transitions (mean) from Markov jumps, both with input trees generated by Bayesian interference; 3) number of expected transitions (mean) from the parsimony reconstruction method with input trees generated using maximum likelihood methods.

| Transition | Counts | Markov jumps | Parsimony |
| --- | --- | --- | --- |
| L1 to L2 | 42 | 43 | 42 |
| L1 to L3/4 | 44 | 38 | 42 |
| L2 to L3/4 | 10 | 10 | 9 |

\*Transitions are identified as changes in the most probable IT level between adjacent nodes (see Fig. 1). Details of level transitions with posterior supports are given in Data Files 2 and 3.

**Table S2.**
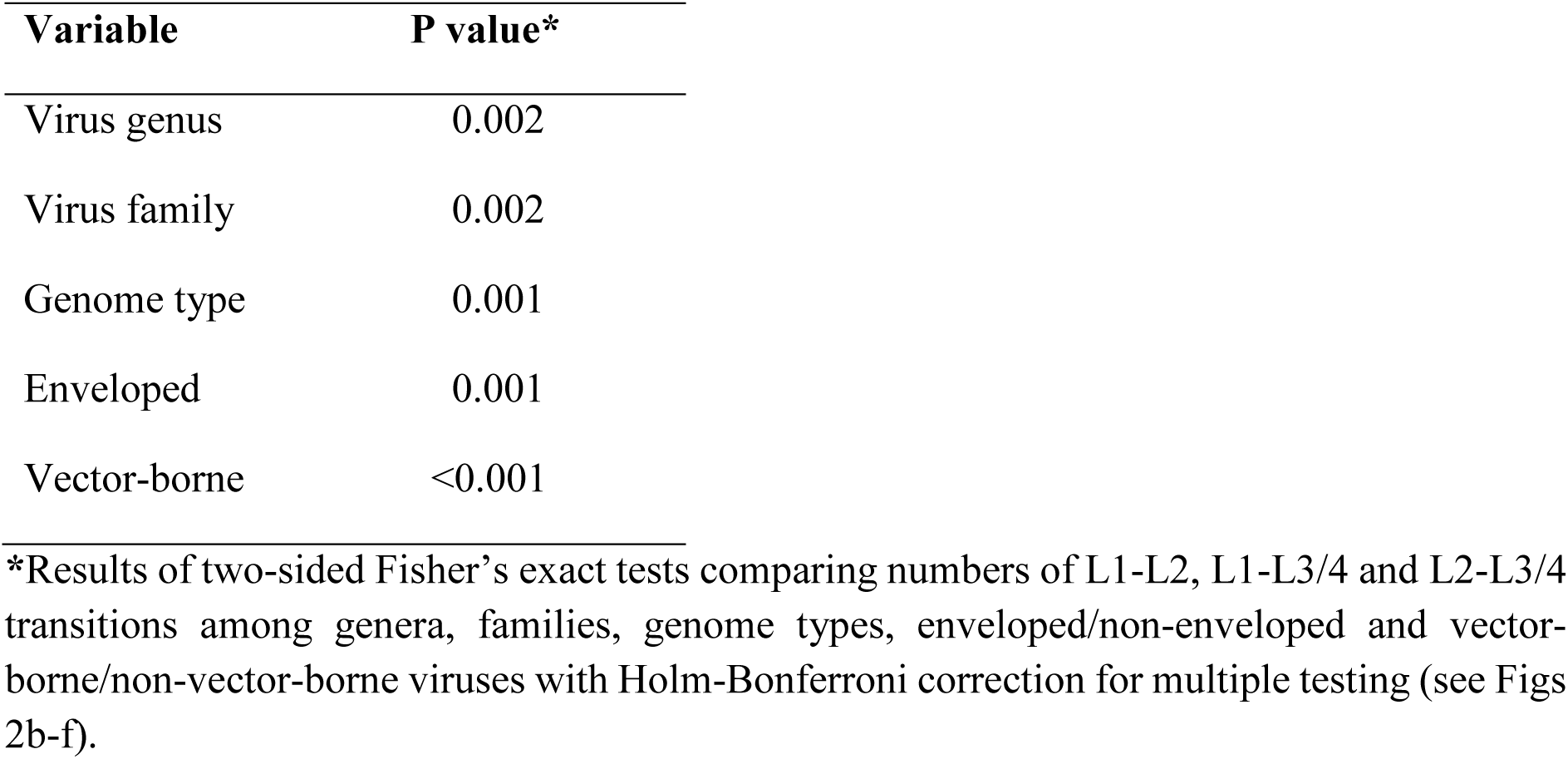
Variation in numbers of types of transitions – univariate analyses.

| **Variable** | **P value\*** |
| --- | --- |
| Virus genus | 0.002 |
| Virus family | 0.002 |
| Genome type | 0.001 |
| Enveloped | 0.001 |
| Vector-borne | <0.001 |

\*Results of two-sided Fisher's exact tests comparing numbers of L1-L2, L1-L3/4 and L2-L3/4 transitions among genera, families, genome types, enveloped/non-enveloped and vector-borne/non-vector-borne viruses with Holm-Bonferroni correction for multiple testing (see Figs 2b-f).

**Table S3.** Results of GLMM with beta distribution and genus as a random factor comparing relative node depth among transition types.

| **Variables** | **Coefficient\*** | **Lower 95% confidence interval** | **Upper 95% confidence interval** | **$X^2$** | **df** | **P value** |
| --- | --- | --- | --- | --- | --- | --- |
| Level change | - | - | - | 0.04 | 2 | 0.98 |
| L1 to L3/4 | 0.02 | -0.41 | 0.45 | - | - | - |
| L2 to L3/4 | 0.08 | -0.64 | 0.79 | - | - | - |

\*Reporting results of Wald test and log-odds and 95% confidence intervals of global model. Reference category is L1 to L2 transitions. N=96.

**Table S4.**
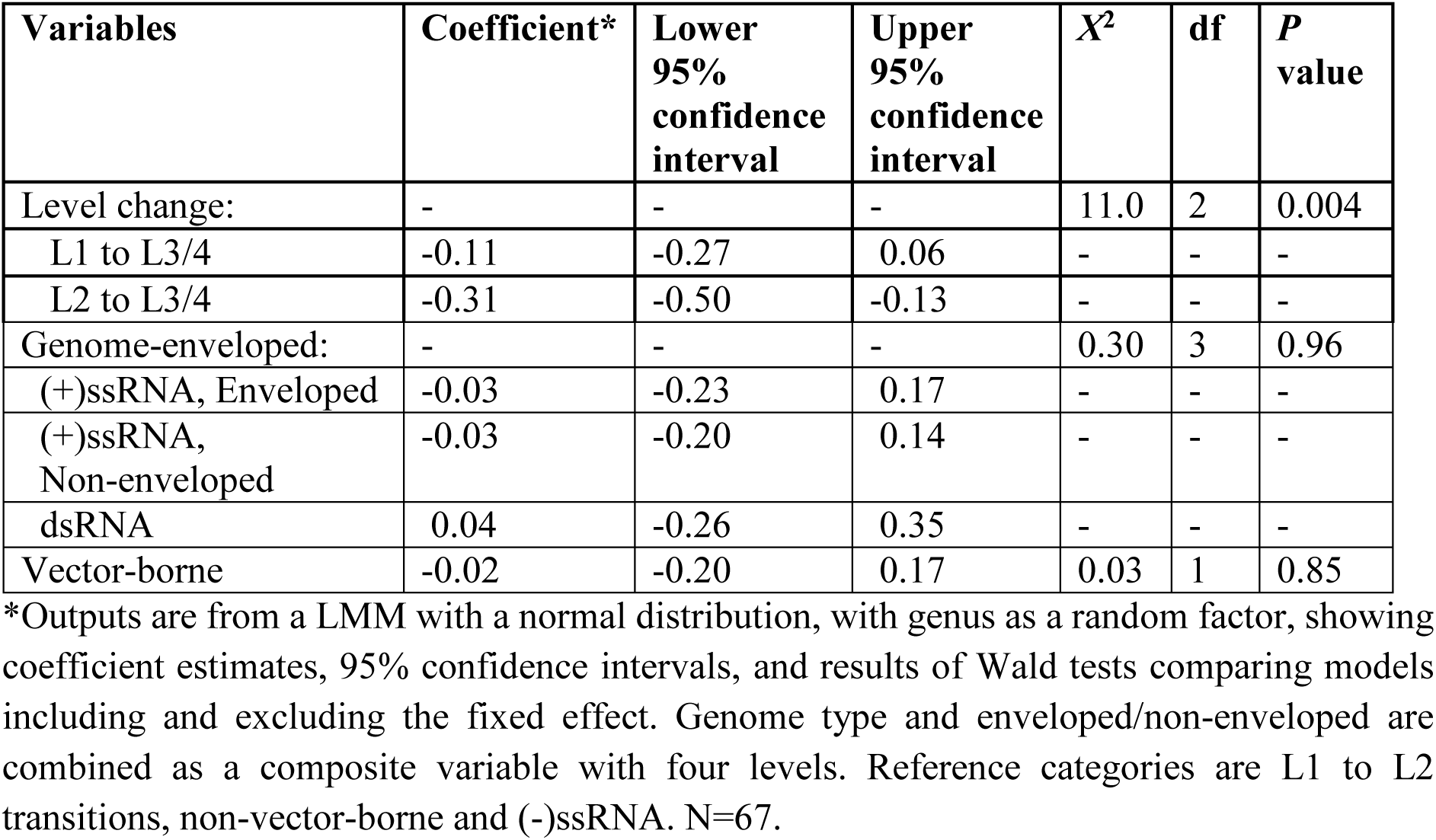
Predictors of loge-transformed genus-level relative transition rates distinguishing L1 to L2, L1 to L3/4 and L2 to L3/4 transitions.

Data File 1

The ancestral level at the root of all genus trees. The posterior support of the ancestral node as well as the posterior probability of each discrete trait state (level) are listed.

Data File 2

Summary of forward transition events (L1-L2, L1-L3/4 and L2-L3/4) and relative node depth. The posterior support of the ancestral node as well as the posterior probability of each discrete trait state (level) are listed.

Data File 3

Summary of reverse transition events (L2-L1, L3/4-L1 and L3/4-L1) and relative node depth. The posterior support of the ancestral node as well as the posterior probability of each discrete trait state (level) are listed.

Data File 4

Sequence data used in this analysis. For each sequence being used in this study, the information of accession number, virus family, genus, species (whether ICTV recognized), transmission level and host type are given.

Data File 5

Maximum clade credibility (MCC) trees of 39 genera mapped with level traits (as shown in Fig. 1). The branches and nodes are coloured according to the most probable ancestral trait inferred by an asymmetric discrete trait model upon the posterior tree set, with correlated trait posterior probabilities indicated by branch widths. Virus species are labelled on tips of the trees. Trees are scaled by number of substitutions per site, with scale bars labelled underneath each genus tree.

Data File 6

Example XML file (*Mamastrovirus*) used in the discrete trait analysis.

Data File 7

Comparison of IT level trait reconstructions from discrete trait models and Parsimony methods.

Data File 8

Surface proteins selected for phylogenetic analysis by genus (N=35).

Data File 9

Comparison of number of L3/4 lineages and number of IT changes found in each genus phylogeny using polymerase and surface protein sequences.

