## Supplementary material for "Evolutionary origins of epidemic potential among human RNA viruses": Data File 5

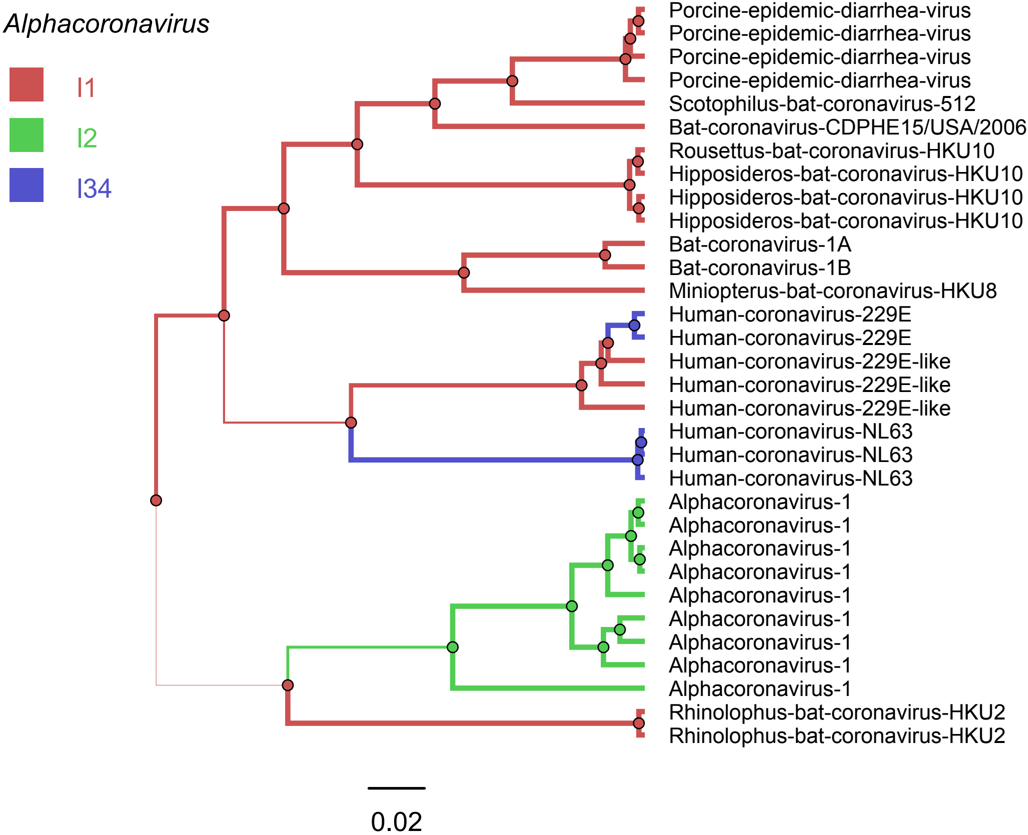
Alphacoronavirus
I1
I2
I34
Porcine-epidemic-diarrhea-virus
Porcine-epidemic-diarrhea-virus
Porcine-epidemic-diarrhea-virus
Porcine-epidemic-diarrhea-virus
Scotophilus-bat-coronavirus-512
Bat-coronavirus-CDPHE15/USA/2006
Rousettus-bat-coronavirus-HKU10
Hipposideros-bat-coronavirus-HKU10
Hipposideros-bat-coronavirus-HKU10
Hipposideros-bat-coronavirus-HKU10
Bat-coronavirus-1A
Bat-coronavirus-1B
Miniopterus-bat-coronavirus-HKU8
Human-coronavirus-229E
Human-coronavirus-229E
Human-coronavirus-229E-like
Human-coronavirus-229E-like
Human-coronavirus-229E-like
Human-coronavirus-NL63
Human-coronavirus-NL63
Human-coronavirus-NL63
Alphacoronavirus-1
Alphacoronavirus-1
Alphacoronavirus-1
Alphacoronavirus-1
Alphacoronavirus-1
Alphacoronavirus-1
Alphacoronavirus-1
Alphacoronavirus-1
Alphacoronavirus-1
Rhinolophus-bat-coronavirus-HKU2
Rhinolophus-bat-coronavirus-HKU2
0.02

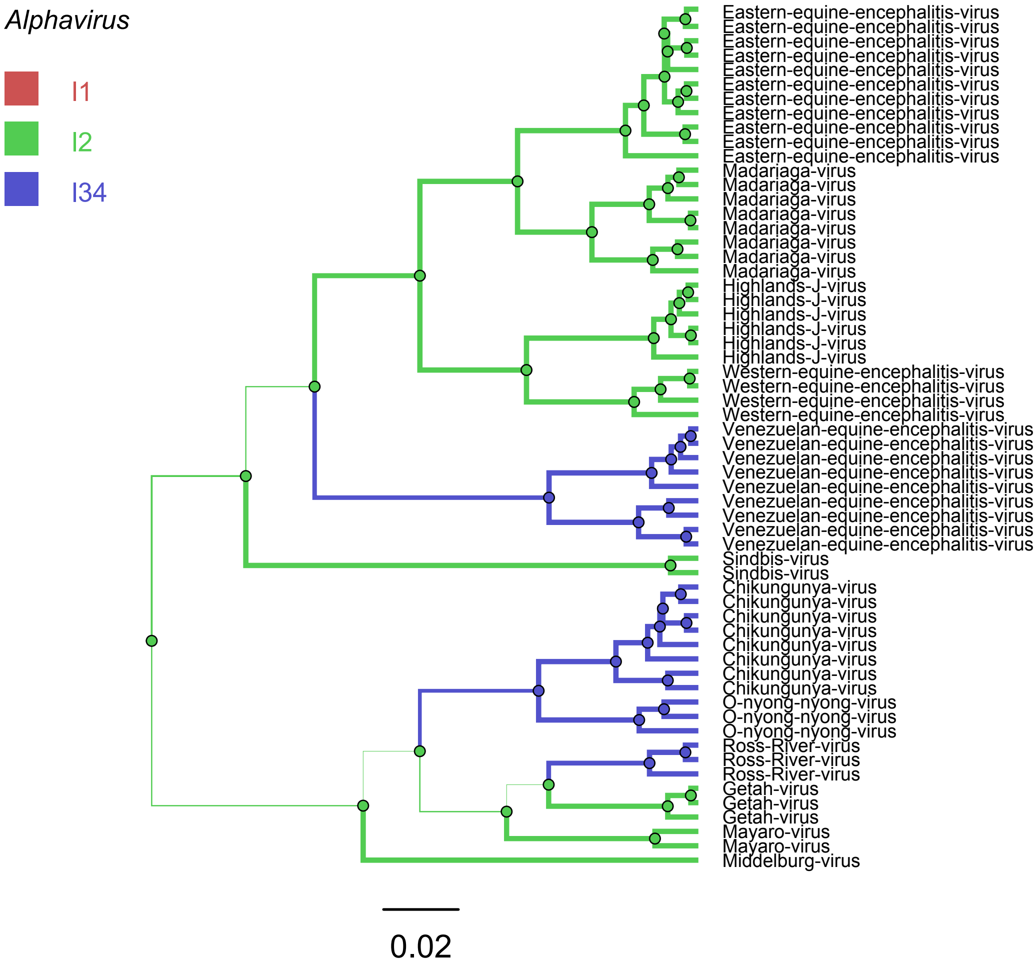

Alphavirus
I1
I2
I34
Eastern-equine-encephalitis-virus
Eastern-equine-encephalitis-virus
Eastern-equine-encephalitis-virus
Eastern-equine-encephalitis-virus
Eastern-equine-encephalitis-virus
Eastern-equine-encephalitis-virus
Eastern-equine-encephalitis-virus
Eastern-equine-encephalitis-virus
Eastern-equine-encephalitis-virus
Eastern-equine-encephalitis-virus
Eastern-equine-encephalitis-virus
Madariaga-virus
Madariaga-virus
Madariaga-virus
Madariaga-virus
Madariaga-virus
Madariaga-virus
Madariaga-virus
Madariaga-virus
Highlands-J-virus
Highlands-J-virus
Highlands-J-virus
Highlands-J-virus
Highlands-J-virus
Highlands-J-virus
Western-equine-encephalitis-virus
Western-equine-encephalitis-virus
Western-equine-encephalitis-virus
Western-equine-encephalitis-virus
Venezuelan-equine-encephalitis-virus
Venezuelan-equine-encephalitis-virus
Venezuelan-equine-encephalitis-virus
Venezuelan-equine-encephalitis-virus
Venezuelan-equine-encephalitis-virus
Venezuelan-equine-encephalitis-virus
Venezuelan-equine-encephalitis-virus
Venezuelan-equine-encephalitis-virus
Venezuelan-equine-encephalitis-virus
Sindbis-virus
Sindbis-virus
Chikungunya-virus
Chikungunya-virus
Chikungunya-virus
Chikungunya-virus
Chikungunya-virus
Chikungunya-virus
Chikungunya-virus
Chikungunya-virus
O-nyong-nyong-virus
O-nyong-nyong-virus
O-nyong-nyong-virus
Ross-River-virus
Ross-River-virus
Ross-River-virus
Getah-virus
Getah-virus
Getah-virus
Mayaro-virus
Mayaro-virus
Middelburg-virus
0.02

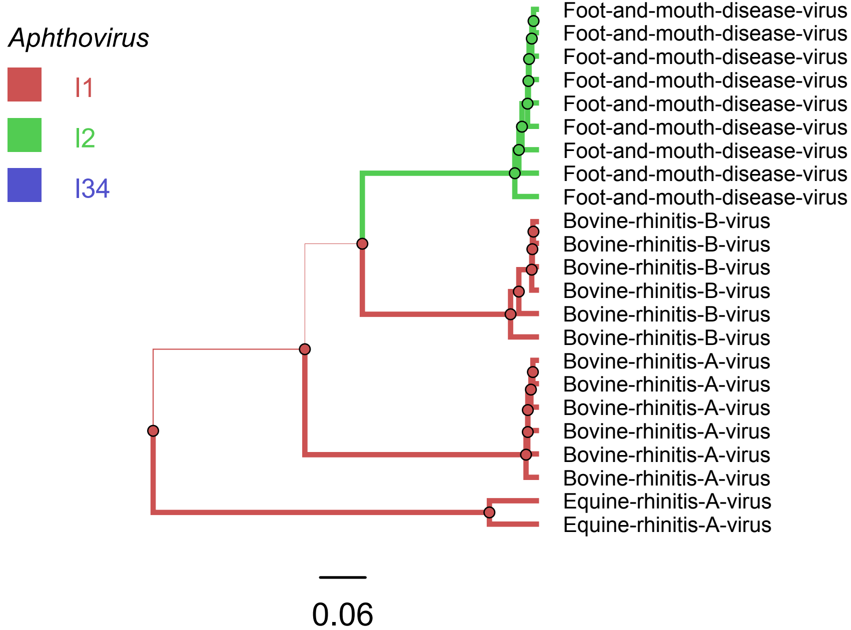
Aphthovirus
I1
I2
I34
Foot-and-mouth-disease-virus
Foot-and-mouth-disease-virus
Foot-and-mouth-disease-virus
Foot-and-mouth-disease-virus
Foot-and-mouth-disease-virus
Foot-and-mouth-disease-virus
Foot-and-mouth-disease-virus
Foot-and-mouth-disease-virus
Foot-and-mouth-disease-virus
Bovine-rhinitis-B-virus
Bovine-rhinitis-B-virus
Bovine-rhinitis-B-virus
Bovine-rhinitis-B-virus
Bovine-rhinitis-B-virus
Bovine-rhinitis-B-virus
Bovine-rhinitis-A-virus
Bovine-rhinitis-A-virus
Bovine-rhinitis-A-virus
Bovine-rhinitis-A-virus
Bovine-rhinitis-A-virus
Bovine-rhinitis-A-virus
Equine-rhinitis-A-virus
Equine-rhinitis-A-virus
0.06

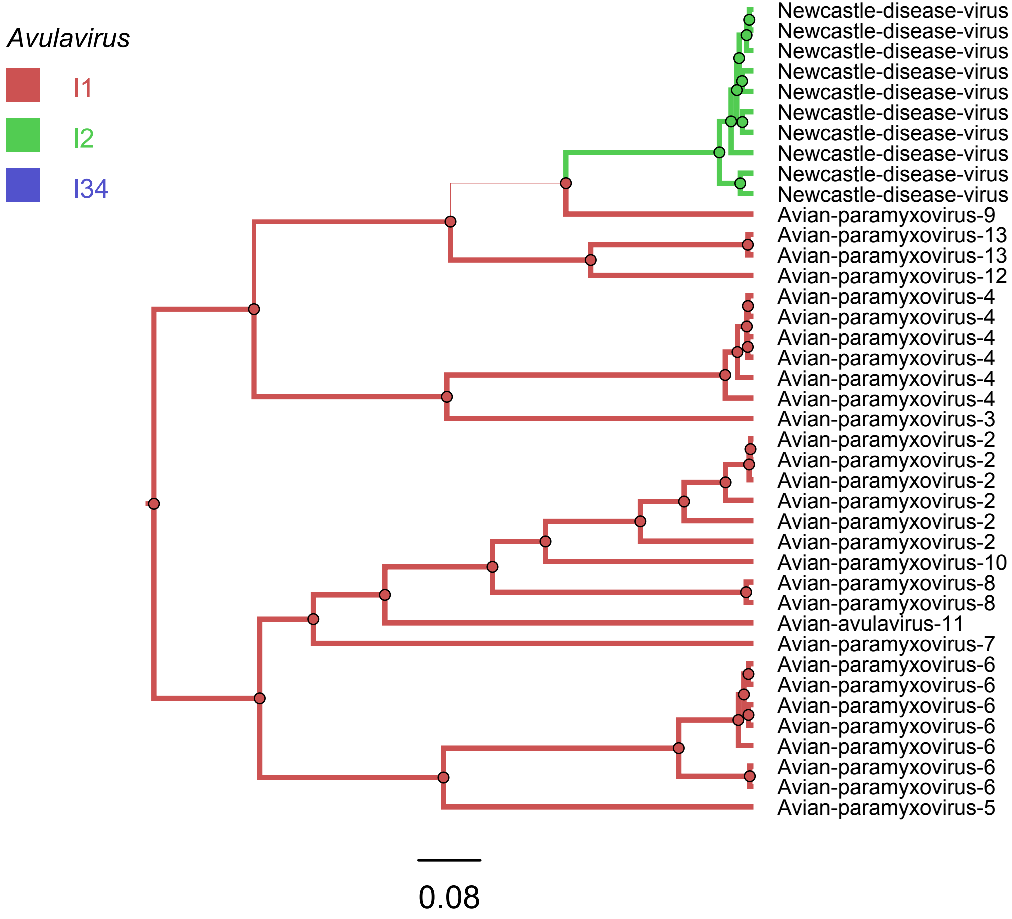

Avulavirus
l1
l2
l34
Newcastle-disease-virus
Newcastle-disease-virus
Newcastle-disease-virus
Newcastle-disease-virus
Newcastle-disease-virus
Newcastle-disease-virus
Newcastle-disease-virus
Newcastle-disease-virus
Newcastle-disease-virus
Newcastle-disease-virus
Avian-paramyxovirus-9
Avian-paramyxovirus-13
Avian-paramyxovirus-13
Avian-paramyxovirus-12
Avian-paramyxovirus-4
Avian-paramyxovirus-4
Avian-paramyxovirus-4
Avian-paramyxovirus-4
Avian-paramyxovirus-4
Avian-paramyxovirus-4
Avian-paramyxovirus-3
Avian-paramyxovirus-2
Avian-paramyxovirus-2
Avian-paramyxovirus-2
Avian-paramyxovirus-2
Avian-paramyxovirus-2
Avian-paramyxovirus-2
Avian-paramyxovirus-10
Avian-paramyxovirus-8
Avian-paramyxovirus-8
Avian-avulavirus-11
Avian-paramyxovirus-7
Avian-paramyxovirus-6
Avian-paramyxovirus-6
Avian-paramyxovirus-6
Avian-paramyxovirus-6
Avian-paramyxovirus-6
Avian-paramyxovirus-6
Avian-paramyxovirus-6
Avian-paramyxovirus-5
0.08

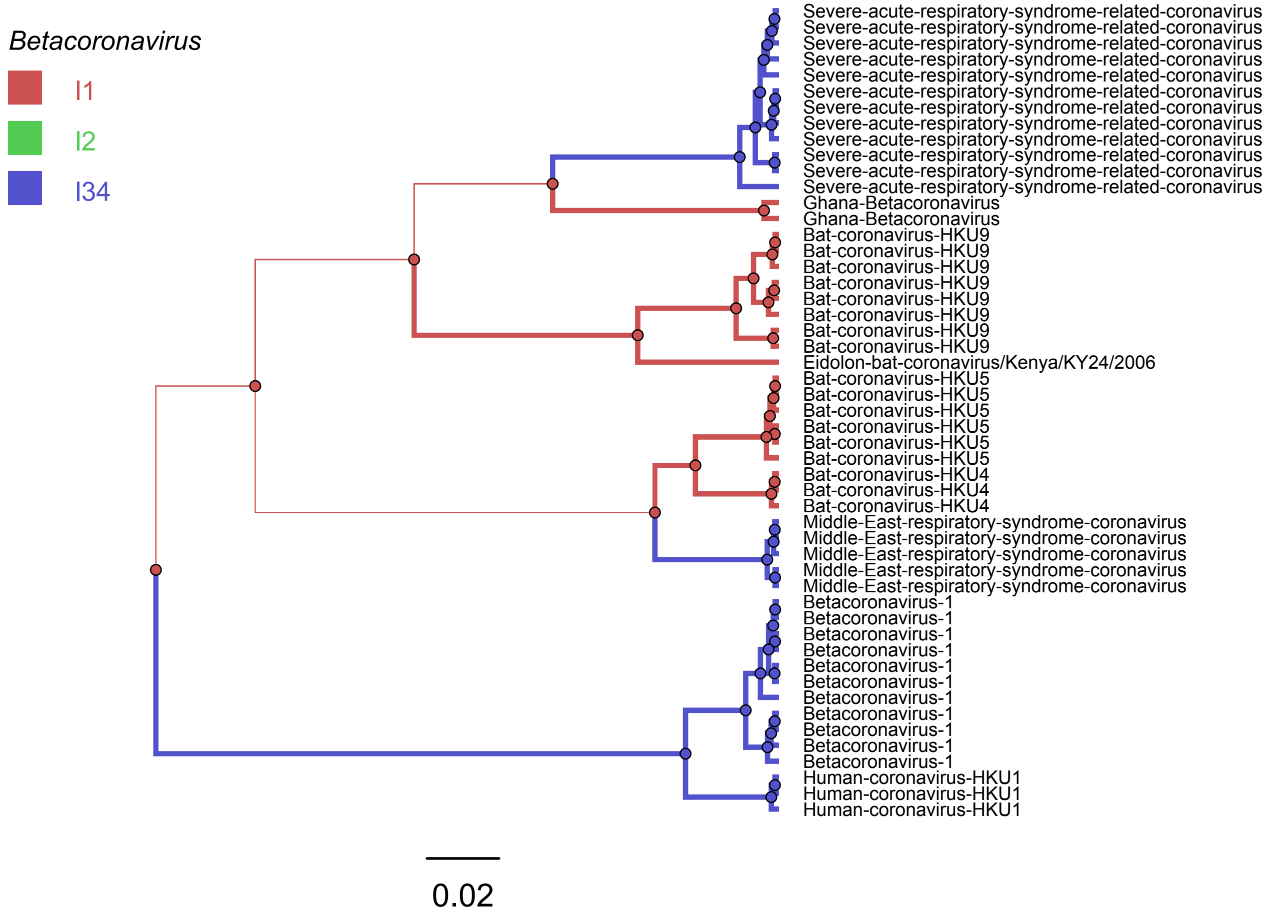

Betacoronavirus
I1
I2
I34
Severe-acute-respiratory-syndrome-related-coronavirus
Severe-acute-respiratory-syndrome-related-coronavirus
Severe-acute-respiratory-syndrome-related-coronavirus
Severe-acute-respiratory-syndrome-related-coronavirus
Severe-acute-respiratory-syndrome-related-coronavirus
Severe-acute-respiratory-syndrome-related-coronavirus
Severe-acute-respiratory-syndrome-related-coronavirus
Severe-acute-respiratory-syndrome-related-coronavirus
Severe-acute-respiratory-syndrome-related-coronavirus
Severe-acute-respiratory-syndrome-related-coronavirus
Severe-acute-respiratory-syndrome-related-coronavirus
Severe-acute-respiratory-syndrome-related-coronavirus
Ghana-Betacoronavirus
Ghana-Betacoronavirus
Bat-coronavirus-HKU9
Bat-coronavirus-HKU9
Bat-coronavirus-HKU9
Bat-coronavirus-HKU9
Bat-coronavirus-HKU9
Bat-coronavirus-HKU9
Bat-coronavirus-HKU9
Bat-coronavirus-HKU9
Eidolon-bat-coronavirus/Kenya/KY24/2006
Bat-coronavirus-HKU5
Bat-coronavirus-HKU5
Bat-coronavirus-HKU5
Bat-coronavirus-HKU5
Bat-coronavirus-HKU5
Bat-coronavirus-HKU5
Bat-coronavirus-HKU4
Bat-coronavirus-HKU4
Bat-coronavirus-HKU4
Middle-East-respiratory-syndrome-coronavirus
Middle-East-respiratory-syndrome-coronavirus
Middle-East-respiratory-syndrome-coronavirus
Middle-East-respiratory-syndrome-coronavirus
Middle-East-respiratory-syndrome-coronavirus
Betacoronavirus-1
Betacoronavirus-1
Betacoronavirus-1
Betacoronavirus-1
Betacoronavirus-1
Betacoronavirus-1
Betacoronavirus-1
Betacoronavirus-1
Betacoronavirus-1
Betacoronavirus-1
Betacoronavirus-1
Human-coronavirus-HKU1
Human-coronavirus-HKU1
Human-coronavirus-HKU1
0.02

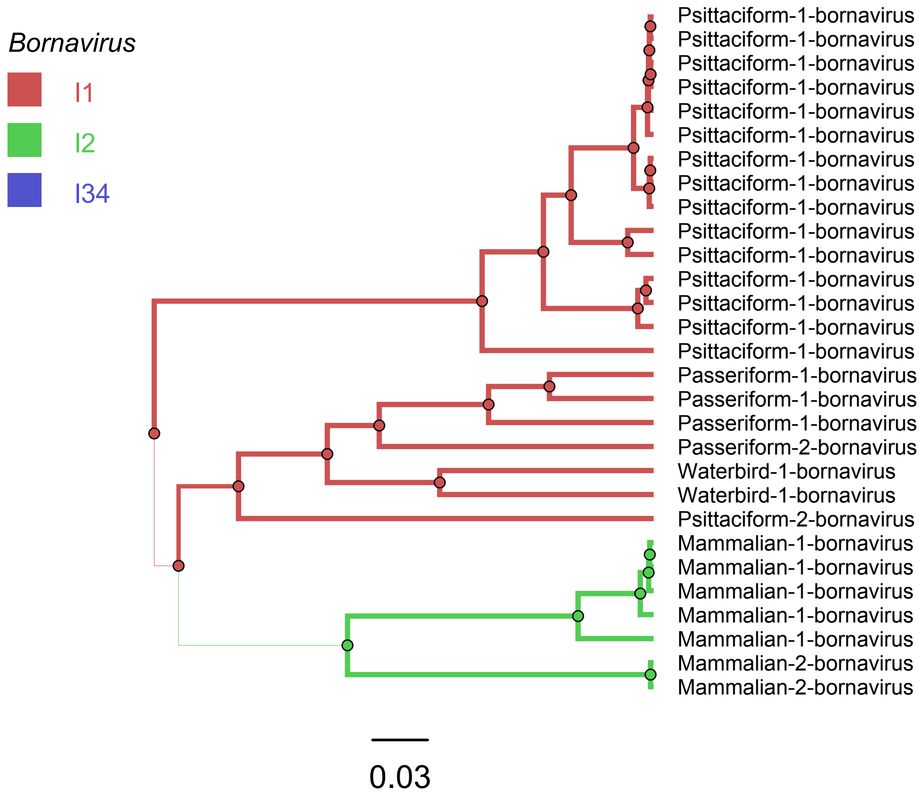
Bornavirus
I1
I2
I34
Psittaciform-1-bornavirus
Psittaciform-1-bornavirus
Psittaciform-1-bornavirus
Psittaciform-1-bornavirus
Psittaciform-1-bornavirus
Psittaciform-1-bornavirus
Psittaciform-1-bornavirus
Psittaciform-1-bornavirus
Psittaciform-1-bornavirus
Psittaciform-1-bornavirus
Psittaciform-1-bornavirus
Psittaciform-1-bornavirus
Psittaciform-1-bornavirus
Psittaciform-1-bornavirus
Psittaciform-1-bornavirus
Passeriform-1-bornavirus
Passeriform-1-bornavirus
Passeriform-1-bornavirus
Passeriform-2-bornavirus
Waterbird-1-bornavirus
Waterbird-1-bornavirus
Psittaciform-2-bornavirus
Mammalian-1-bornavirus
Mammalian-1-bornavirus
Mammalian-1-bornavirus
Mammalian-1-bornavirus
Mammalian-1-bornavirus
Mammalian-2-bornavirus
Mammalian-2-bornavirus
0.03

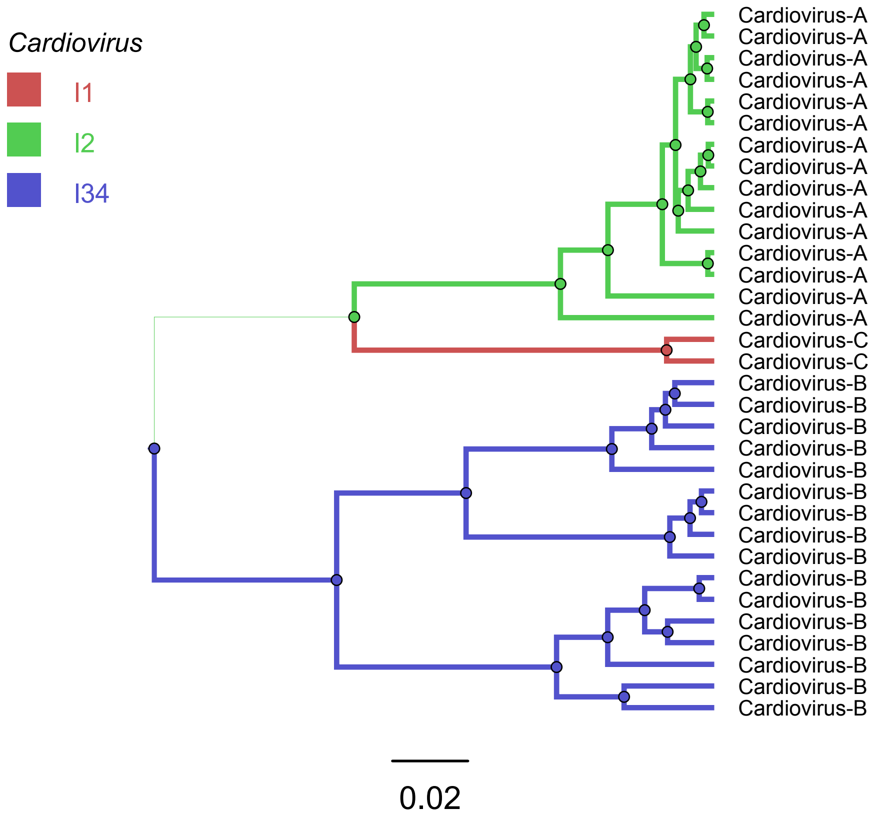

Cardiovirus
I1
I2
I34
Cardiovirus-A
Cardiovirus-A
Cardiovirus-A
Cardiovirus-A
Cardiovirus-A
Cardiovirus-A
Cardiovirus-A
Cardiovirus-A
Cardiovirus-A
Cardiovirus-A
Cardiovirus-A
Cardiovirus-A
Cardiovirus-A
Cardiovirus-A
Cardiovirus-A
Cardiovirus-C
Cardiovirus-C
Cardiovirus-B
Cardiovirus-B
Cardiovirus-B
Cardiovirus-B
Cardiovirus-B
Cardiovirus-B
Cardiovirus-B
Cardiovirus-B
Cardiovirus-B
Cardiovirus-B
Cardiovirus-B
Cardiovirus-B
Cardiovirus-B
Cardiovirus-B
Cardiovirus-B
Cardiovirus-B
0.02

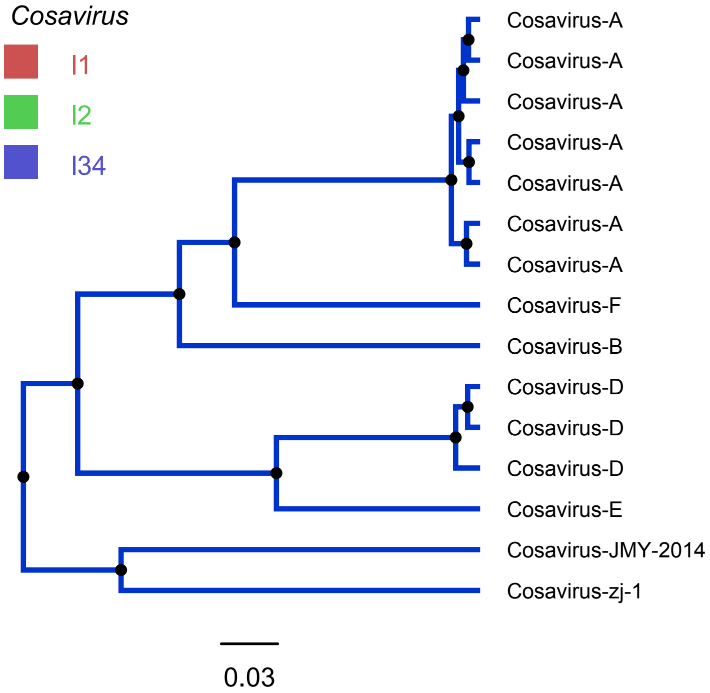

Cosavirus
I1
I2
I34
Cosavirus-A
Cosavirus-A
Cosavirus-A
Cosavirus-A
Cosavirus-A
Cosavirus-A
Cosavirus-A
Cosavirus-F
Cosavirus-B
Cosavirus-D
Cosavirus-D
Cosavirus-D
Cosavirus-E
Cosavirus-JMY-2014
Cosavirus-zj-1
0.03

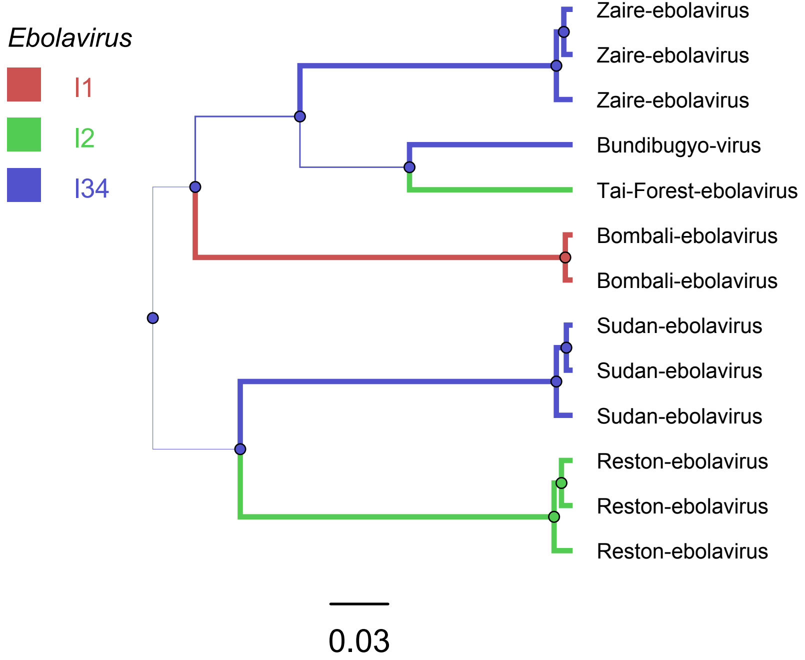
Ebolavirus
I1
I2
I34
Zaire-ebolavirus
Zaire-ebolavirus
Zaire-ebolavirus
Bundibugyo-virus
Tai-Forest-ebolavirus
Bombali-ebolavirus
Bombali-ebolavirus
Sudan-ebolavirus
Sudan-ebolavirus
Sudan-ebolavirus
Reston-ebolavirus
Reston-ebolavirus
Reston-ebolavirus
0.03

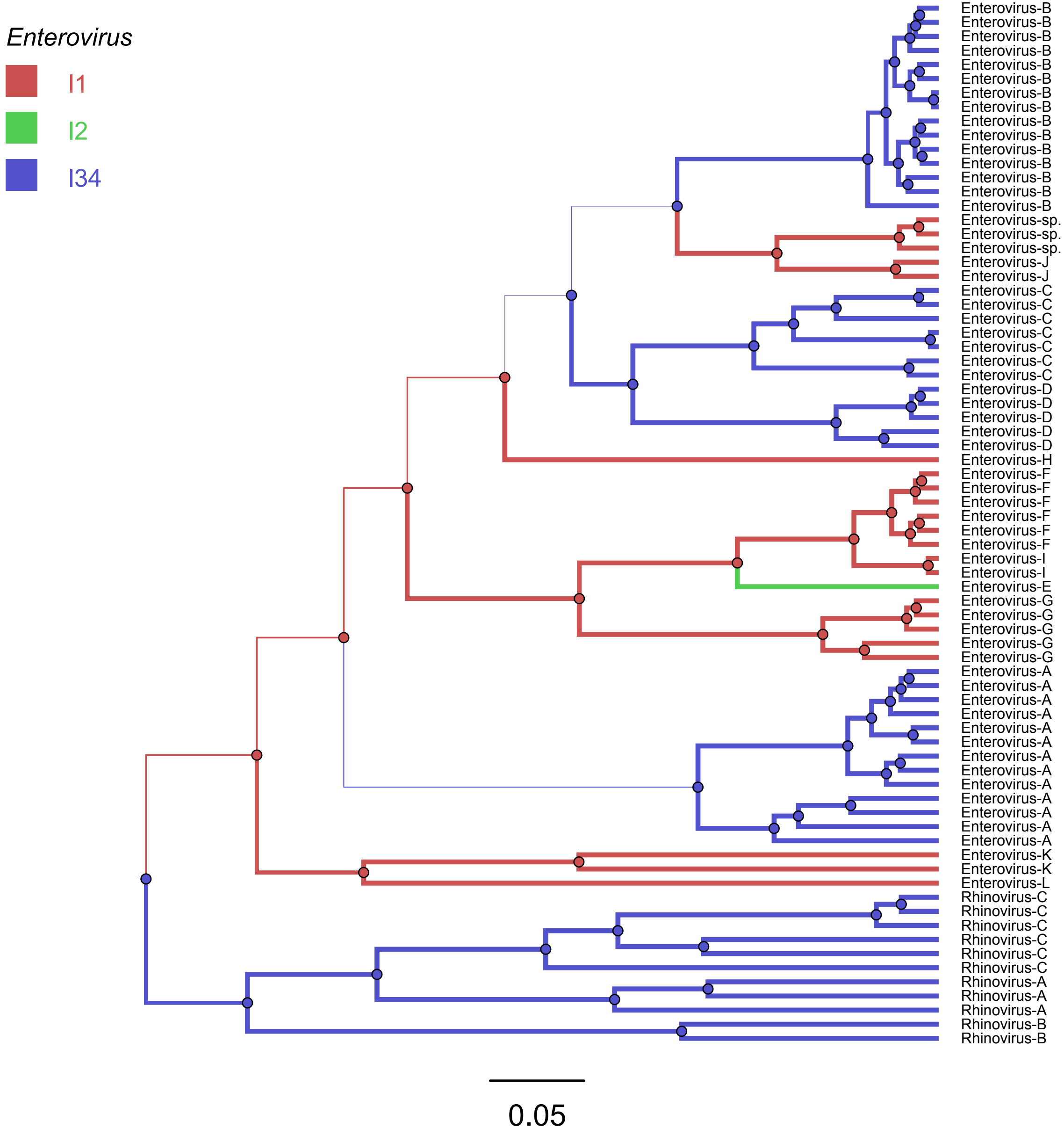
Enterovirus
I1
I2
I34
Enterovirus-B
Enterovirus-B
Enterovirus-B
Enterovirus-B
Enterovirus-B
Enterovirus-B
Enterovirus-B
Enterovirus-B
Enterovirus-B
Enterovirus-B
Enterovirus-B
Enterovirus-B
Enterovirus-B
Enterovirus-B
Enterovirus-B
Enterovirus-sp.
Enterovirus-sp.
Enterovirus-sp.
Enterovirus-J
Enterovirus-J
Enterovirus-C
Enterovirus-C
Enterovirus-C
Enterovirus-C
Enterovirus-C
Enterovirus-C
Enterovirus-C
Enterovirus-D
Enterovirus-D
Enterovirus-D
Enterovirus-D
Enterovirus-D
Enterovirus-H
Enterovirus-F
Enterovirus-F
Enterovirus-F
Enterovirus-F
Enterovirus-F
Enterovirus-F
Enterovirus-I
Enterovirus-I
Enterovirus-E
Enterovirus-G
Enterovirus-G
Enterovirus-G
Enterovirus-G
Enterovirus-G
Enterovirus-A
Enterovirus-A
Enterovirus-A
Enterovirus-A
Enterovirus-A
Enterovirus-A
Enterovirus-A
Enterovirus-A
Enterovirus-A
Enterovirus-A
Enterovirus-A
Enterovirus-A
Enterovirus-A
Enterovirus-K
Enterovirus-K
Enterovirus-L
Rhinovirus-C
Rhinovirus-C
Rhinovirus-C
Rhinovirus-C
Rhinovirus-C
Rhinovirus-C
Rhinovirus-A
Rhinovirus-A
Rhinovirus-A
Rhinovirus-B
Rhinovirus-B
0.05

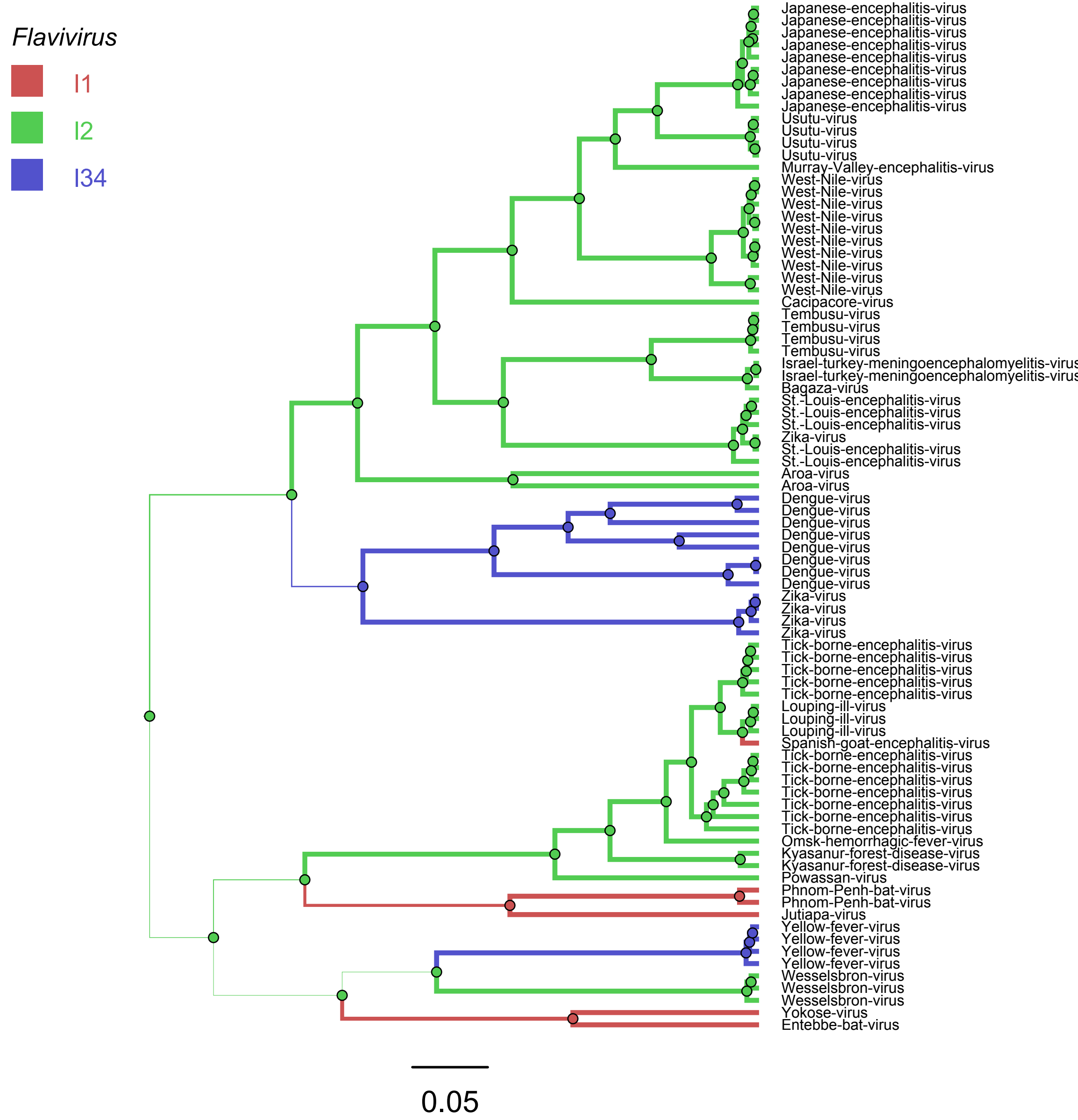
Flavivirus
I1
I2
I34
Japanese-encephalitis-virus
Japanese-encephalitis-virus
Japanese-encephalitis-virus
Japanese-encephalitis-virus
Japanese-encephalitis-virus
Japanese-encephalitis-virus
Japanese-encephalitis-virus
Japanese-encephalitis-virus
Japanese-encephalitis-virus
Usutu-virus
Usutu-virus
Usutu-virus
Usutu-virus
Murray-Valley-encephalitis-virus
West-Nile-virus
West-Nile-virus
West-Nile-virus
West-Nile-virus
West-Nile-virus
West-Nile-virus
West-Nile-virus
West-Nile-virus
West-Nile-virus
West-Nile-virus
Cacipacore-virus
Tembusu-virus
Tembusu-virus
Tembusu-virus
Tembusu-virus
Israel-turkey-meningoencephalomyelitis-virus
Israel-turkey-meningoencephalomyelitis-virus
Bagaza-virus
St.-Louis-encephalitis-virus
St.-Louis-encephalitis-virus
St.-Louis-encephalitis-virus
Zika-virus
St.-Louis-encephalitis-virus
St.-Louis-encephalitis-virus
Aroa-virus
Aroa-virus
Dengue-virus
Dengue-virus
Dengue-virus
Dengue-virus
Dengue-virus
Dengue-virus
Dengue-virus
Dengue-virus
Zika-virus
Zika-virus
Zika-virus
Zika-virus
Tick-borne-encephalitis-virus
Tick-borne-encephalitis-virus
Tick-borne-encephalitis-virus
Tick-borne-encephalitis-virus
Tick-borne-encephalitis-virus
Louping-ill-virus
Louping-ill-virus
Louping-ill-virus
Spanish-goat-encephalitis-virus
Tick-borne-encephalitis-virus
Tick-borne-encephalitis-virus
Tick-borne-encephalitis-virus
Tick-borne-encephalitis-virus
Tick-borne-encephalitis-virus
Tick-borne-encephalitis-virus
Tick-borne-encephalitis-virus
Omsk-hemorrhagic-fever-virus
Kyasanur-forest-disease-virus
Kyasanur-forest-disease-virus
Powassan-virus
Phnom-Penh-bat-virus
Phnom-Penh-bat-virus
Jutiapa-virus
Yellow-fever-virus
Yellow-fever-virus
Yellow-fever-virus
Yellow-fever-virus
Wesselsbron-virus
Wesselsbron-virus
Wesselsbron-virus
Yokose-virus
Entebbe-bat-virus
0.05

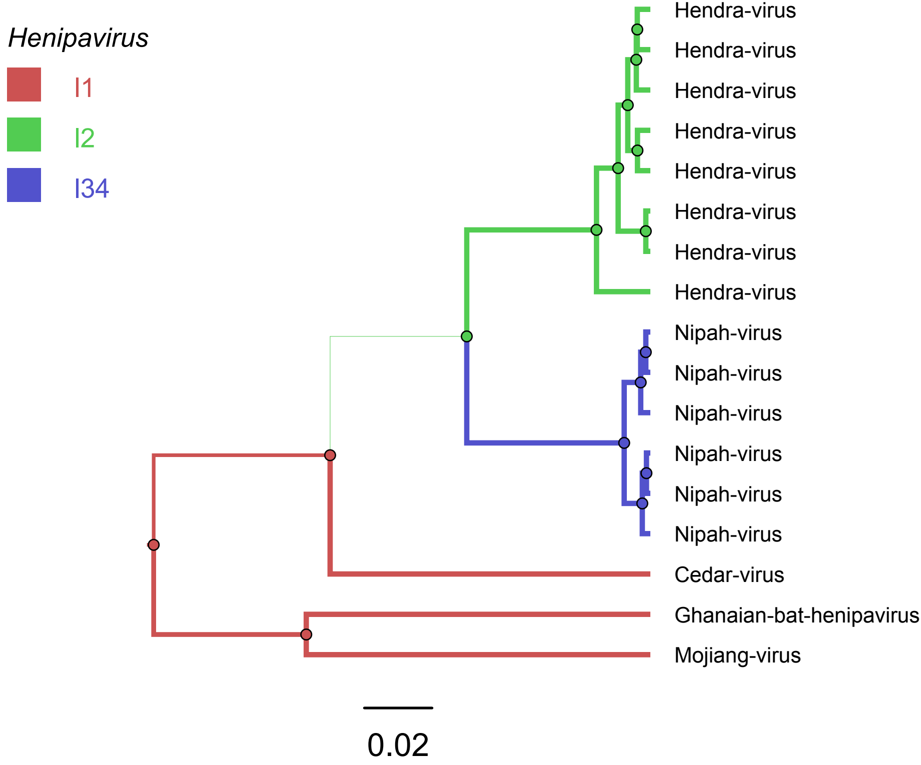
Henipavirus
I1
I2
I34
Hendra-virus
Hendra-virus
Hendra-virus
Hendra-virus
Hendra-virus
Hendra-virus
Hendra-virus
Hendra-virus
Nipah-virus
Nipah-virus
Nipah-virus
Nipah-virus
Nipah-virus
Nipah-virus
Cedar-virus
Ghanaian-bat-henipavirus
Mojiang-virus
0.02

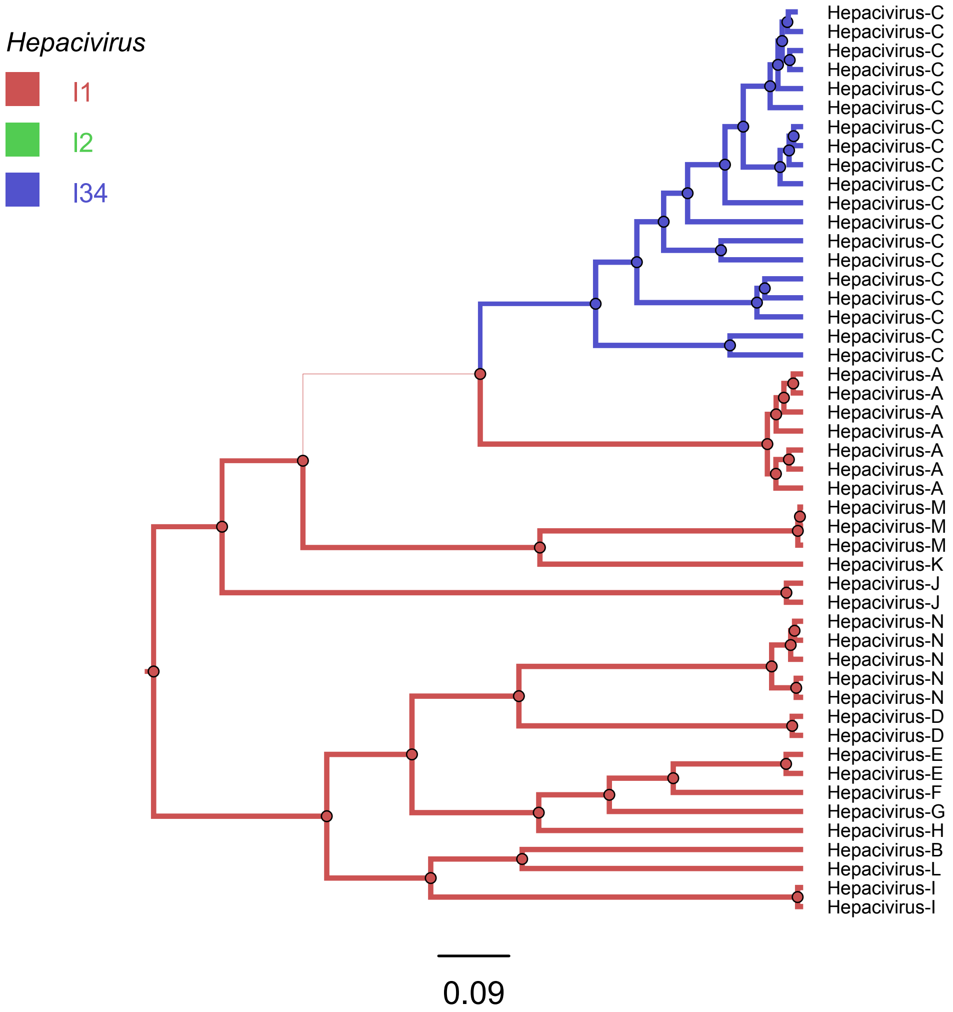
Hepacivirus
I1
I2
I34
Hepacivirus-C
Hepacivirus-C
Hepacivirus-C
Hepacivirus-C
Hepacivirus-C
Hepacivirus-C
Hepacivirus-C
Hepacivirus-C
Hepacivirus-C
Hepacivirus-C
Hepacivirus-C
Hepacivirus-C
Hepacivirus-C
Hepacivirus-C
Hepacivirus-C
Hepacivirus-C
Hepacivirus-C
Hepacivirus-C
Hepacivirus-C
Hepacivirus-A
Hepacivirus-A
Hepacivirus-A
Hepacivirus-A
Hepacivirus-A
Hepacivirus-A
Hepacivirus-A
Hepacivirus-M
Hepacivirus-M
Hepacivirus-M
Hepacivirus-K
Hepacivirus-J
Hepacivirus-J
Hepacivirus-N
Hepacivirus-N
Hepacivirus-N
Hepacivirus-N
Hepacivirus-N
Hepacivirus-D
Hepacivirus-D
Hepacivirus-E
Hepacivirus-E
Hepacivirus-F
Hepacivirus-G
Hepacivirus-H
Hepacivirus-B
Hepacivirus-L
Hepacivirus-I
Hepacivirus-I
0.09

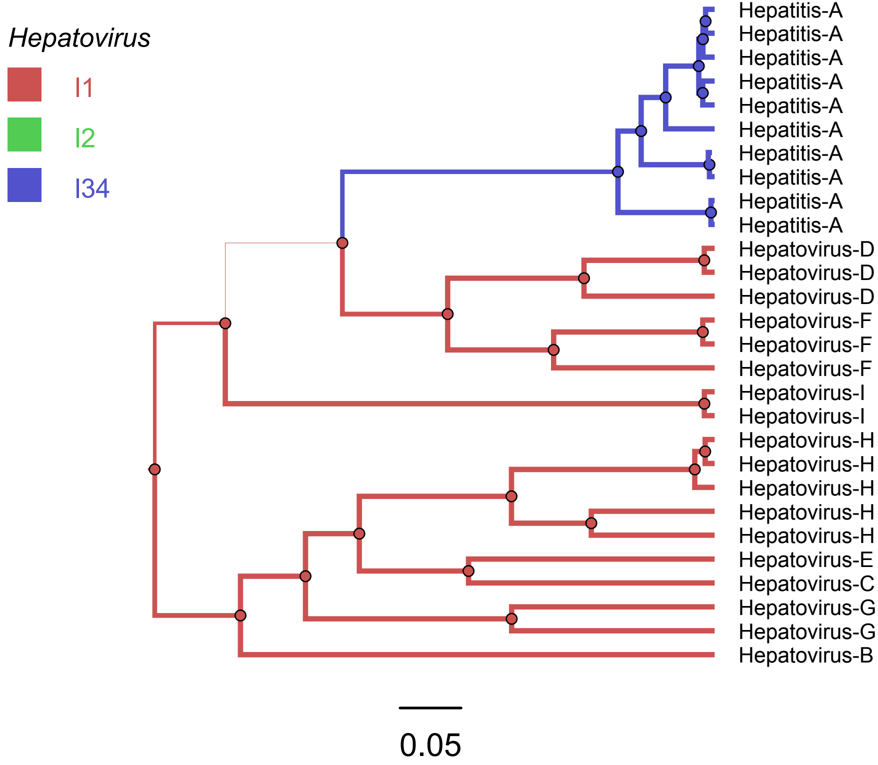
Hepatovirus
I1
I2
I34
Hepatitis-A
Hepatitis-A
Hepatitis-A
Hepatitis-A
Hepatitis-A
Hepatitis-A
Hepatitis-A
Hepatitis-A
Hepatitis-A
Hepatitis-A
Hepatovirus-D
Hepatovirus-D
Hepatovirus-D
Hepatovirus-F
Hepatovirus-F
Hepatovirus-F
Hepatovirus-I
Hepatovirus-I
Hepatovirus-H
Hepatovirus-H
Hepatovirus-H
Hepatovirus-H
Hepatovirus-H
Hepatovirus-E
Hepatovirus-C
Hepatovirus-G
Hepatovirus-G
Hepatovirus-B
0.05

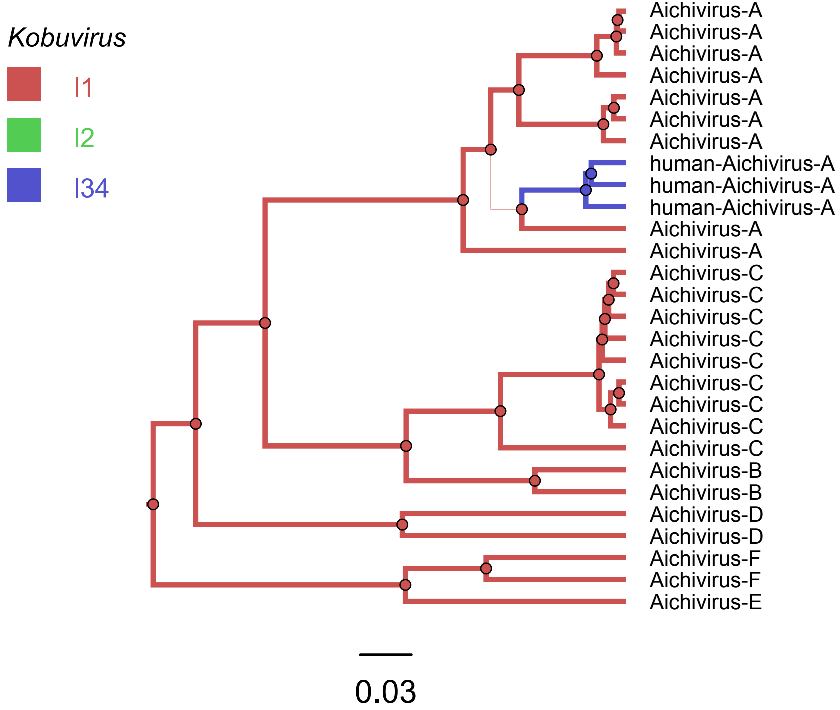
Kobuvirus
I1
I2
I34
Aichivirus-A
Aichivirus-A
Aichivirus-A
Aichivirus-A
Aichivirus-A
Aichivirus-A
Aichivirus-A
human-Aichivirus-A
human-Aichivirus-A
human-Aichivirus-A
Aichivirus-A
Aichivirus-A
Aichivirus-C
Aichivirus-C
Aichivirus-C
Aichivirus-C
Aichivirus-C
Aichivirus-C
Aichivirus-C
Aichivirus-C
Aichivirus-C
Aichivirus-B
Aichivirus-B
Aichivirus-D
Aichivirus-D
Aichivirus-F
Aichivirus-F
Aichivirus-E
0.03

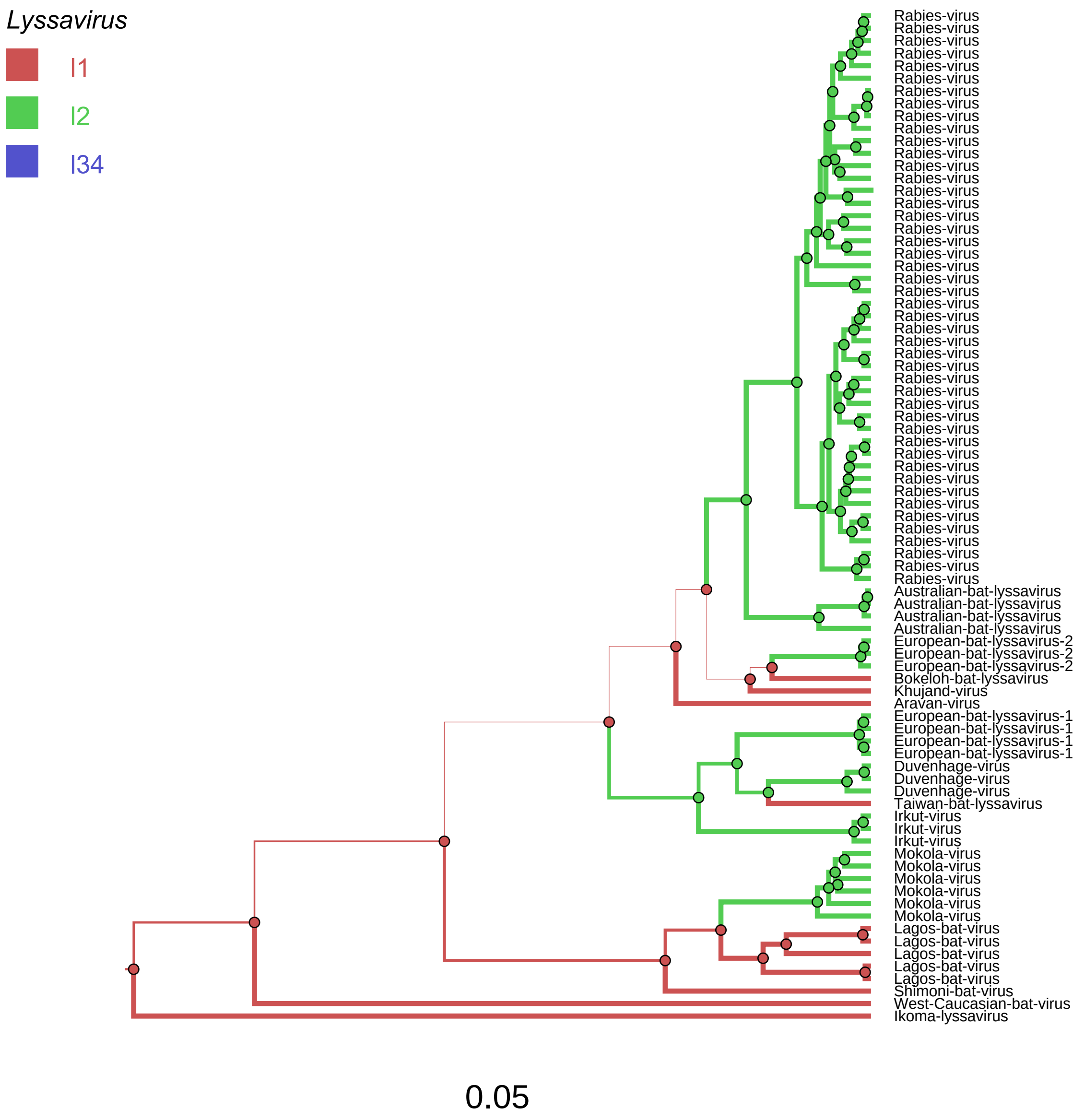

Lyssavirus
I1
I2
I34
Rabies-virus
Australian-bat-lyssavirus
European-bat-lyssavirus-2
Bokeloh-bat-lyssavirus
Khujand-virus
Aravan-virus
European-bat-lyssavirus-1
Duvenhage-virus
Taiwan-bat-lyssavirus
Irkut-virus
Mokola-virus
Lagos-bat-virus
Shimoni-bat-virus
West-Caucasian-bat-virus
Ikoma-lyssavirus
0.05

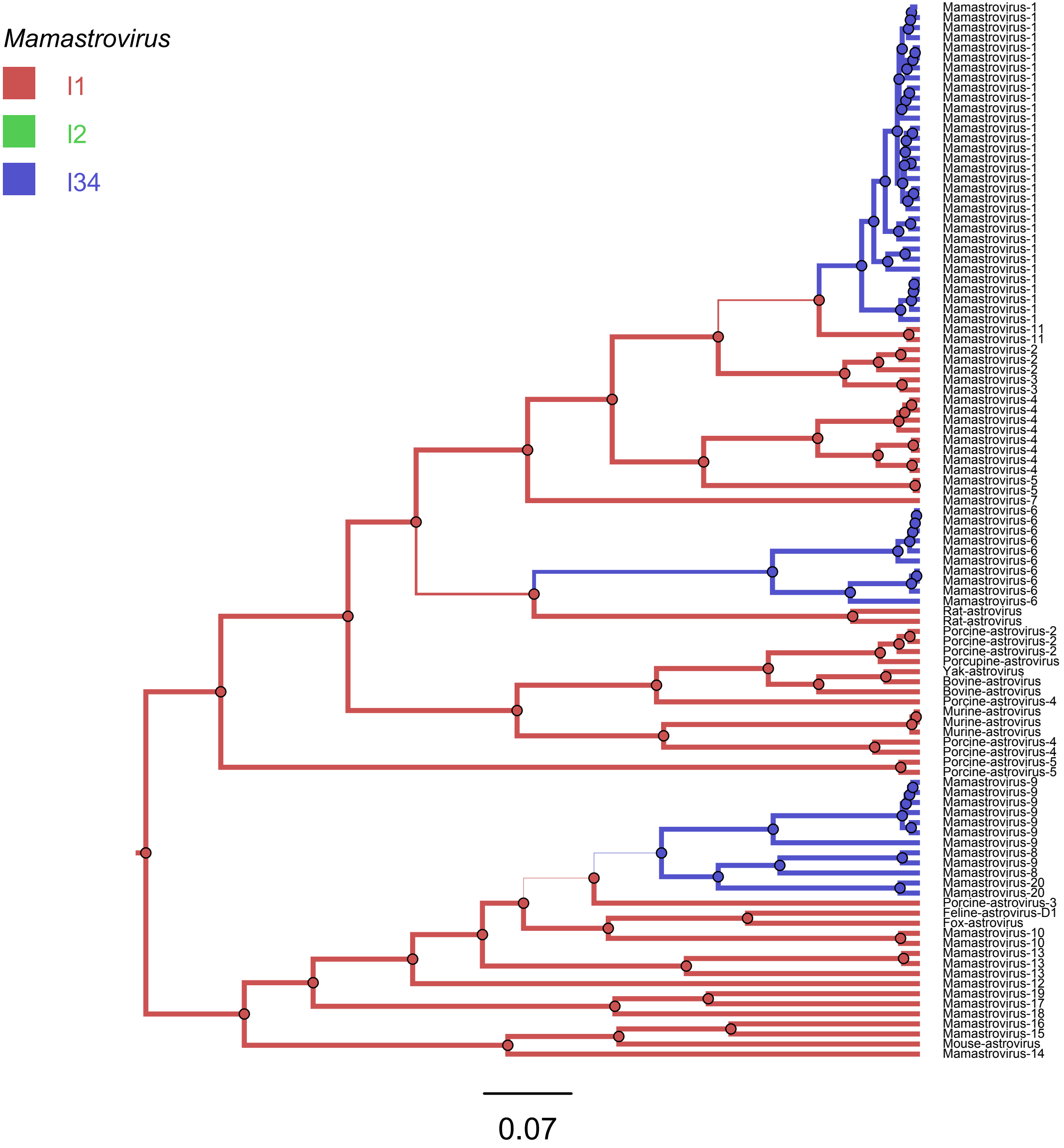

Mamastrovirus
I1
I2
I34
Mamastrovirus-1
Mamastrovirus-1
Mamastrovirus-1
Mamastrovirus-1
Mamastrovirus-1
Mamastrovirus-1
Mamastrovirus-1
Mamastrovirus-1
Mamastrovirus-1
Mamastrovirus-1
Mamastrovirus-1
Mamastrovirus-1
Mamastrovirus-1
Mamastrovirus-1
Mamastrovirus-1
Mamastrovirus-1
Mamastrovirus-1
Mamastrovirus-1
Mamastrovirus-1
Mamastrovirus-1
Mamastrovirus-1
Mamastrovirus-1
Mamastrovirus-1
Mamastrovirus-1
Mamastrovirus-1
Mamastrovirus-1
Mamastrovirus-1
Mamastrovirus-1
Mamastrovirus-1
Mamastrovirus-1
Mamastrovirus-1
Mamastrovirus-1
Mamastrovirus-11
Mamastrovirus-11
Mamastrovirus-2
Mamastrovirus-2
Mamastrovirus-2
Mamastrovirus-3
Mamastrovirus-3
Mamastrovirus-4
Mamastrovirus-4
Mamastrovirus-4
Mamastrovirus-4
Mamastrovirus-4
Mamastrovirus-4
Mamastrovirus-4
Mamastrovirus-4
Mamastrovirus-5
Mamastrovirus-5
Mamastrovirus-7
Mamastrovirus-6
Mamastrovirus-6
Mamastrovirus-6
Mamastrovirus-6
Mamastrovirus-6
Mamastrovirus-6
Mamastrovirus-6
Mamastrovirus-6
Mamastrovirus-6
Mamastrovirus-6
Rat-astrovirus
Rat-astrovirus
Porcine-astrovirus-2
Porcine-astrovirus-2
Porcine-astrovirus-2
Porcupine-astrovirus
Yak-astrovirus
Bovine-astrovirus
Bovine-astrovirus
Porcine-astrovirus-4
Murine-astrovirus
Murine-astrovirus
Murine-astrovirus
Porcine-astrovirus-4
Porcine-astrovirus-4
Porcine-astrovirus-5
Porcine-astrovirus-5
Mamastrovirus-9
Mamastrovirus-9
Mamastrovirus-9
Mamastrovirus-9
Mamastrovirus-9
Mamastrovirus-9
Mamastrovirus-9
Mamastrovirus-8
Mamastrovirus-9
Mamastrovirus-8
Mamastrovirus-20
Mamastrovirus-20
Porcine-astrovirus-3
Feline-astrovirus-D1
Fox-astrovirus
Mamastrovirus-10
Mamastrovirus-10
Mamastrovirus-13
Mamastrovirus-13
Mamastrovirus-13
Mamastrovirus-12
Mamastrovirus-19
Mamastrovirus-17
Mamastrovirus-18
Mamastrovirus-16
Mamastrovirus-15
Mouse-astrovirus
Mamastrovirus-14
0.07

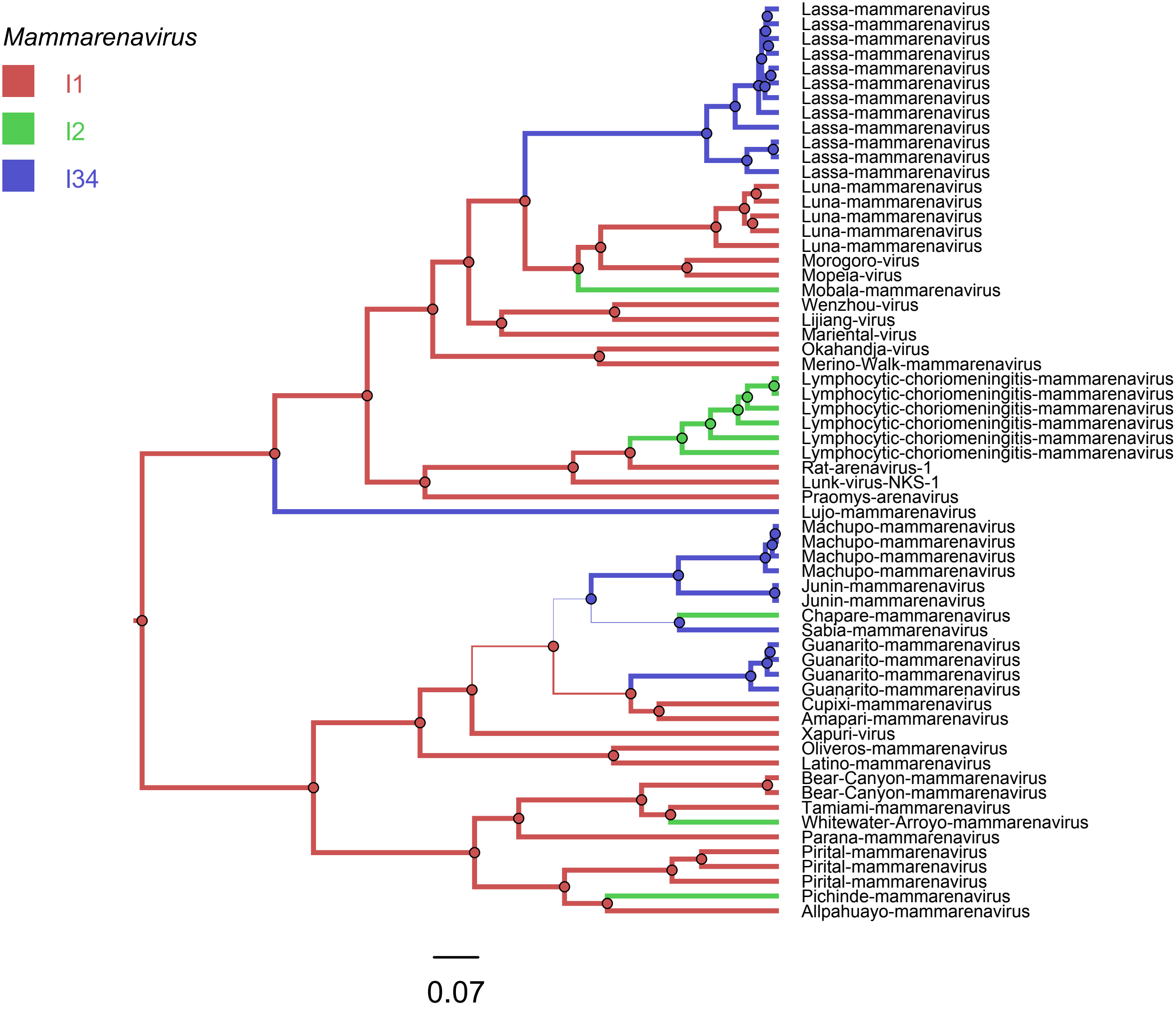

Mammarenavirus
I1
I2
I34
Lassa-mammarenavirus
Lassa-mammarenavirus
Lassa-mammarenavirus
Lassa-mammarenavirus
Lassa-mammarenavirus
Lassa-mammarenavirus
Lassa-mammarenavirus
Lassa-mammarenavirus
Lassa-mammarenavirus
Lassa-mammarenavirus
Lassa-mammarenavirus
Lassa-mammarenavirus
Luna-mammarenavirus
Luna-mammarenavirus
Luna-mammarenavirus
Luna-mammarenavirus
Luna-mammarenavirus
Morogoro-virus
Mopeia-virus
Mobala-mammarenavirus
Wenzhou-virus
Lijiang-virus
Mariental-virus
Okahandja-virus
Merino-Walk-mammarenavirus
Lymphocytic-choriomeningitis-mammarenavirus
Lymphocytic-choriomeningitis-mammarenavirus
Lymphocytic-choriomeningitis-mammarenavirus
Lymphocytic-choriomeningitis-mammarenavirus
Lymphocytic-choriomeningitis-mammarenavirus
Lymphocytic-choriomeningitis-mammarenavirus
Rat-arenavirus-1
Lunk-virus-NKS-1
Praomys-arenavirus
Lujo-mammarenavirus
Machupo-mammarenavirus
Machupo-mammarenavirus
Machupo-mammarenavirus
Machupo-mammarenavirus
Junin-mammarenavirus
Junin-mammarenavirus
Chapare-mammarenavirus
Sabia-mammarenavirus
Guanarito-mammarenavirus
Guanarito-mammarenavirus
Guanarito-mammarenavirus
Guanarito-mammarenavirus
Cupixi-mammarenavirus
Amapari-mammarenavirus
Xapuri-virus
Oliveros-mammarenavirus
Latino-mammarenavirus
Bear-Canyon-mammarenavirus
Bear-Canyon-mammarenavirus
Tamiami-mammarenavirus
Whitewater-Arroyo-mammarenavirus
Parana-mammarenavirus
Pirital-mammarenavirus
Pirital-mammarenavirus
Pirital-mammarenavirus
Pichinde-mammarenavirus
Allpahuayo-mammarenavirus
0.07

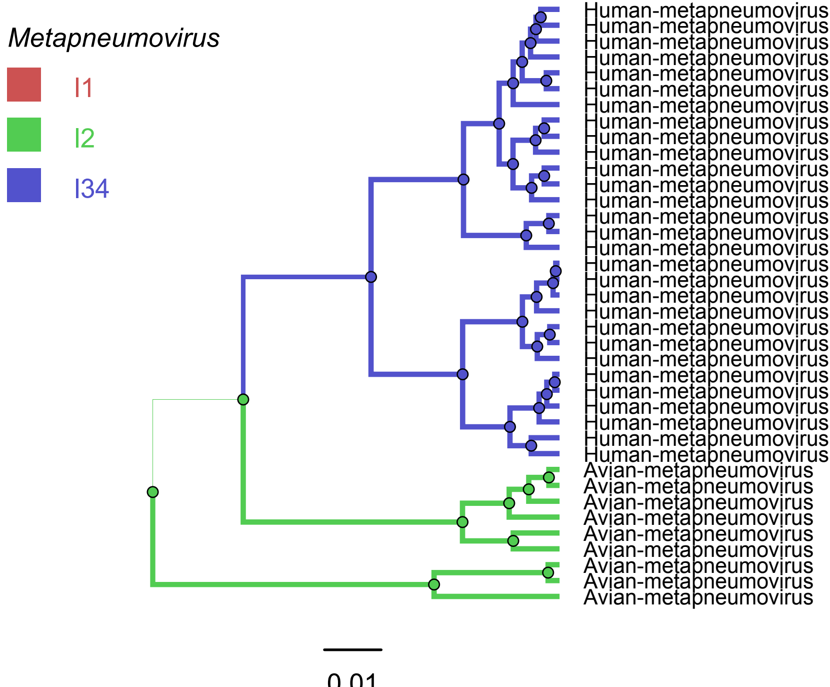
Metapneumovirus
I1
I2
I34
Human-metapneumovirus
Human-metapneumovirus
Human-metapneumovirus
Human-metapneumovirus
Human-metapneumovirus
Human-metapneumovirus
Human-metapneumovirus
Human-metapneumovirus
Human-metapneumovirus
Human-metapneumovirus
Human-metapneumovirus
Human-metapneumovirus
Human-metapneumovirus
Human-metapneumovirus
Human-metapneumovirus
Human-metapneumovirus
Human-metapneumovirus
Human-metapneumovirus
Human-metapneumovirus
Human-metapneumovirus
Human-metapneumovirus
Human-metapneumovirus
Human-metapneumovirus
Human-metapneumovirus
Human-metapneumovirus
Human-metapneumovirus
Human-metapneumovirus
Human-metapneumovirus
Human-metapneumovirus
Avian-metapneumovirus
Avian-metapneumovirus
Avian-metapneumovirus
Avian-metapneumovirus
Avian-metapneumovirus
Avian-metapneumovirus
Avian-metapneumovirus
Avian-metapneumovirus
Avian-metapneumovirus
0.01

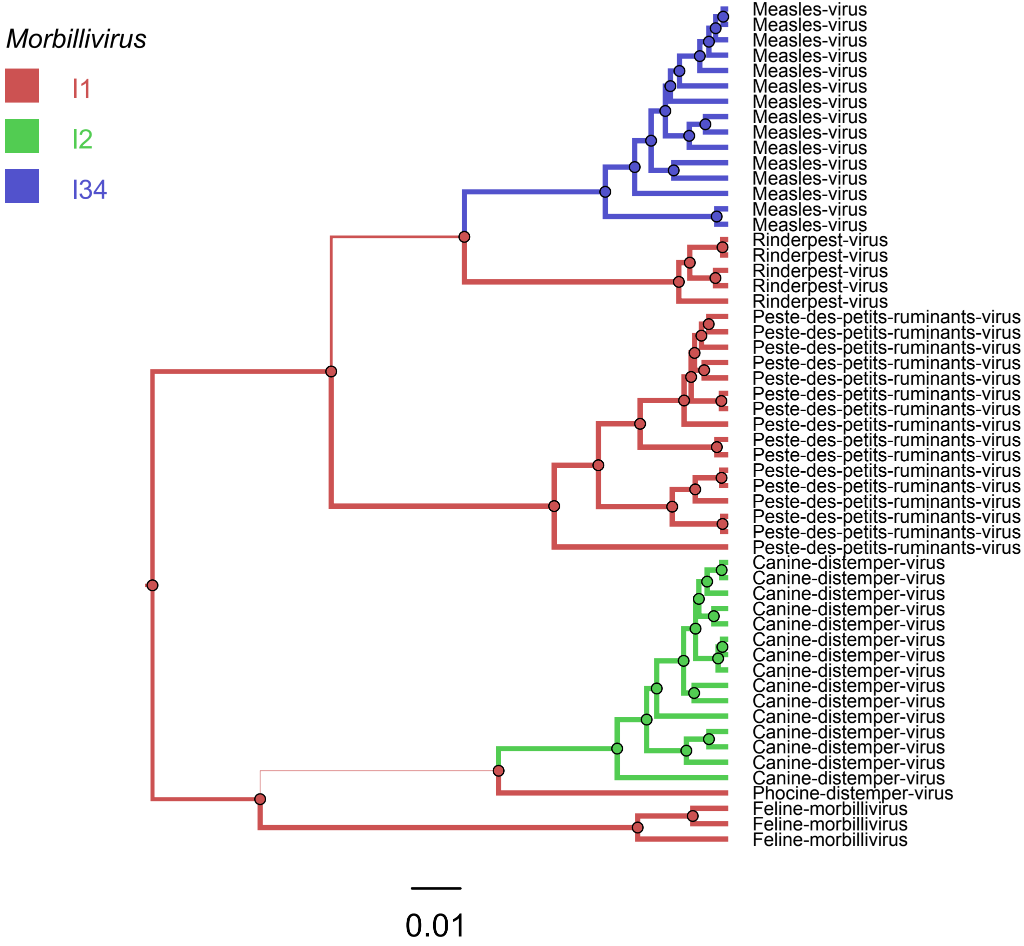
Morbillivirus
l1
l2
l34
Measles-virus
Measles-virus
Measles-virus
Measles-virus
Measles-virus
Measles-virus
Measles-virus
Measles-virus
Measles-virus
Measles-virus
Measles-virus
Measles-virus
Measles-virus
Measles-virus
Measles-virus
Rinderpest-virus
Rinderpest-virus
Rinderpest-virus
Rinderpest-virus
Rinderpest-virus
Peste-des-petits-ruminants-virus
Peste-des-petits-ruminants-virus
Peste-des-petits-ruminants-virus
Peste-des-petits-ruminants-virus
Peste-des-petits-ruminants-virus
Peste-des-petits-ruminants-virus
Peste-des-petits-ruminants-virus
Peste-des-petits-ruminants-virus
Peste-des-petits-ruminants-virus
Peste-des-petits-ruminants-virus
Peste-des-petits-ruminants-virus
Peste-des-petits-ruminants-virus
Peste-des-petits-ruminants-virus
Peste-des-petits-ruminants-virus
Peste-des-petits-ruminants-virus
Peste-des-petits-ruminants-virus
Canine-distemper-virus
Canine-distemper-virus
Canine-distemper-virus
Canine-distemper-virus
Canine-distemper-virus
Canine-distemper-virus
Canine-distemper-virus
Canine-distemper-virus
Canine-distemper-virus
Canine-distemper-virus
Canine-distemper-virus
Canine-distemper-virus
Canine-distemper-virus
Canine-distemper-virus
Canine-distemper-virus
Phocine-distemper-virus
Feline-morbillivirus
Feline-morbillivirus
Feline-morbillivirus
0.01

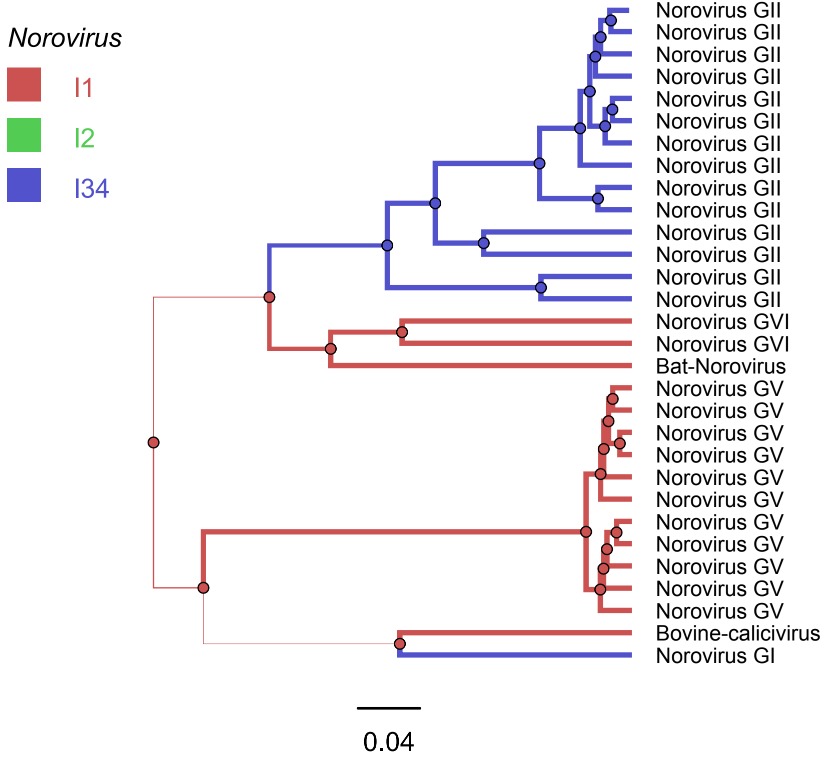
Norovirus
I1
I2
I34
Norovirus GII
Norovirus GII
Norovirus GII
Norovirus GII
Norovirus GII
Norovirus GII
Norovirus GII
Norovirus GII
Norovirus GII
Norovirus GII
Norovirus GII
Norovirus GII
Norovirus GII
Norovirus GII
Norovirus GVI
Norovirus GVI
Bat-Norovirus
Norovirus GV
Norovirus GV
Norovirus GV
Norovirus GV
Norovirus GV
Norovirus GV
Norovirus GV
Norovirus GV
Norovirus GV
Norovirus GV
Norovirus GV
Bovine-calicivirus
Norovirus GI
0.04

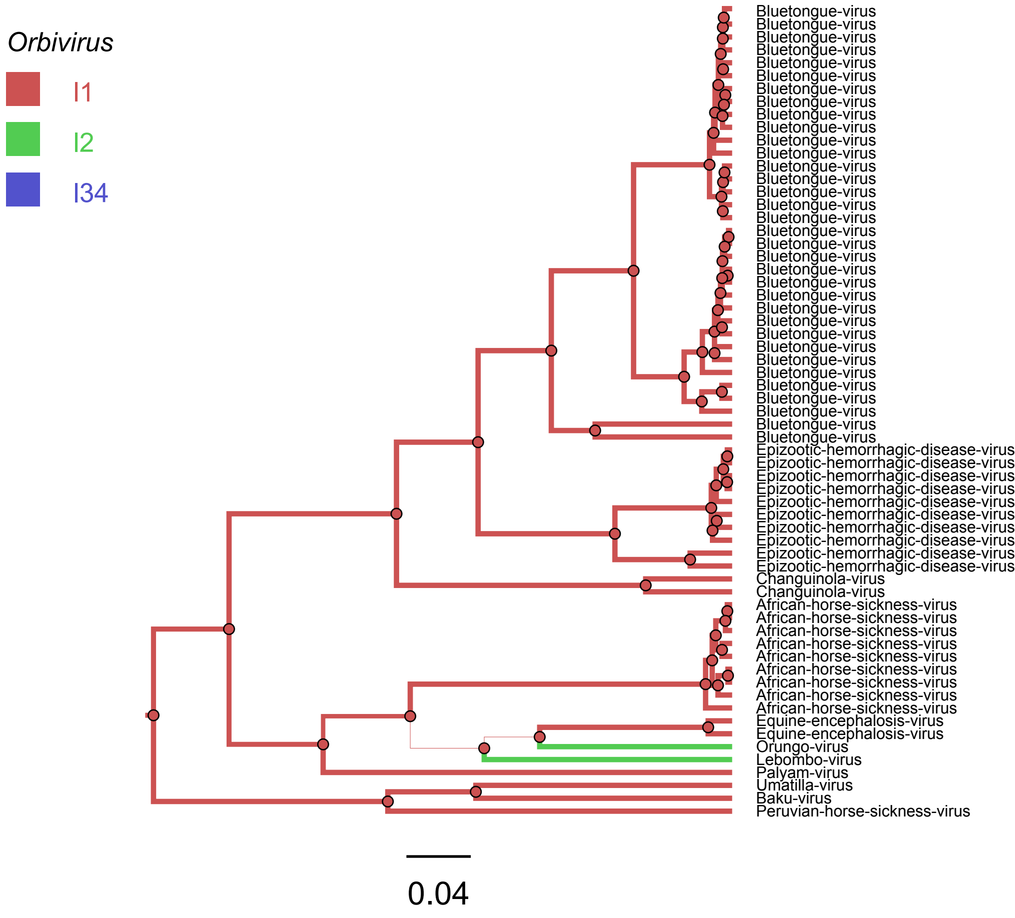

Orbivirus
I1
I2
I34
Bluetongue-virus
Epizootic-hemorrhagic-disease-virus
Changuinola-virus
African-horse-sickness-virus
Equine-encephalosis-virus
Orungo-virus
Lebombo-virus
Palyam-virus
Umatilla-virus
Baku-virus
Peruvian-horse-sickness-virus
0.04

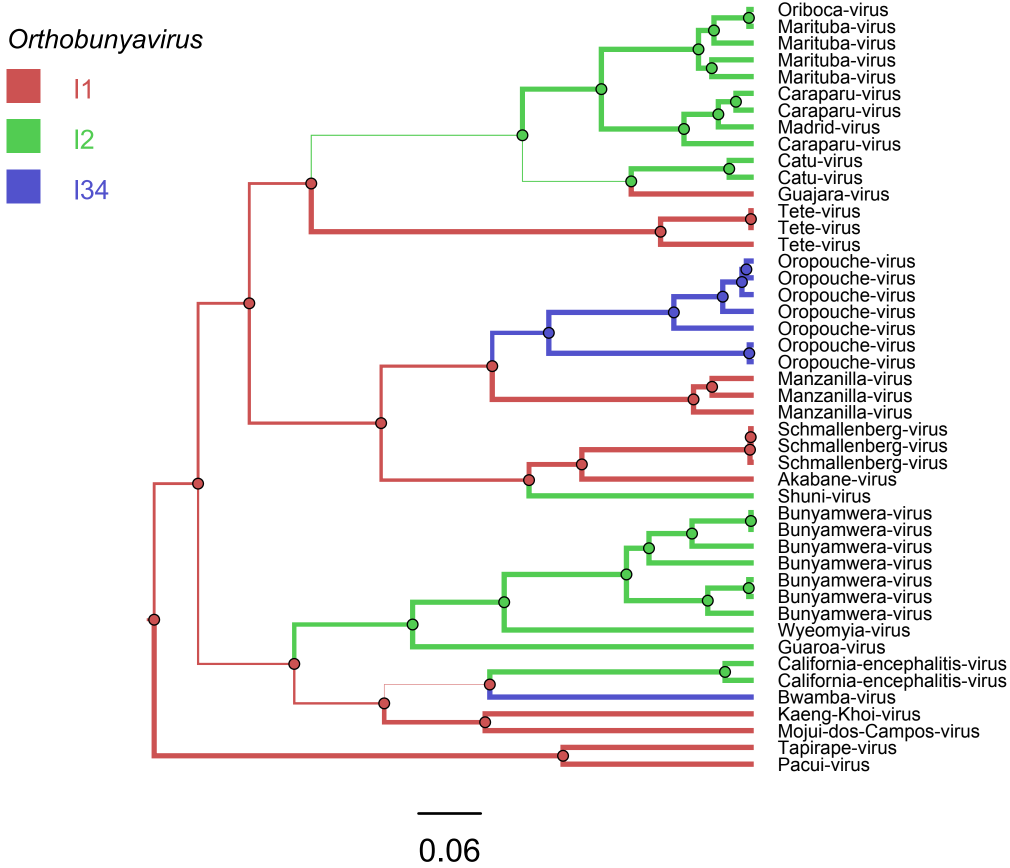
Orthobunyavirus
I1
I2
I34
Oriboca-virus
Marituba-virus
Marituba-virus
Marituba-virus
Marituba-virus
Caraparu-virus
Caraparu-virus
Madrid-virus
Caraparu-virus
Catu-virus
Catu-virus
Guajara-virus
Tete-virus
Tete-virus
Tete-virus
Oropouche-virus
Oropouche-virus
Oropouche-virus
Oropouche-virus
Oropouche-virus
Oropouche-virus
Oropouche-virus
Manzanilla-virus
Manzanilla-virus
Manzanilla-virus
Schmallenberg-virus
Schmallenberg-virus
Schmallenberg-virus
Akabane-virus
Shuni-virus
Bunyamwera-virus
Bunyamwera-virus
Bunyamwera-virus
Bunyamwera-virus
Bunyamwera-virus
Bunyamwera-virus
Bunyamwera-virus
Wyeomyia-virus
Guaroa-virus
California-encephalitis-virus
California-encephalitis-virus
Bwamba-virus
Kaeng-Khoi-virus
Mojui-dos-Campos-virus
Tapirape-virus
Pacui-virus
0.06

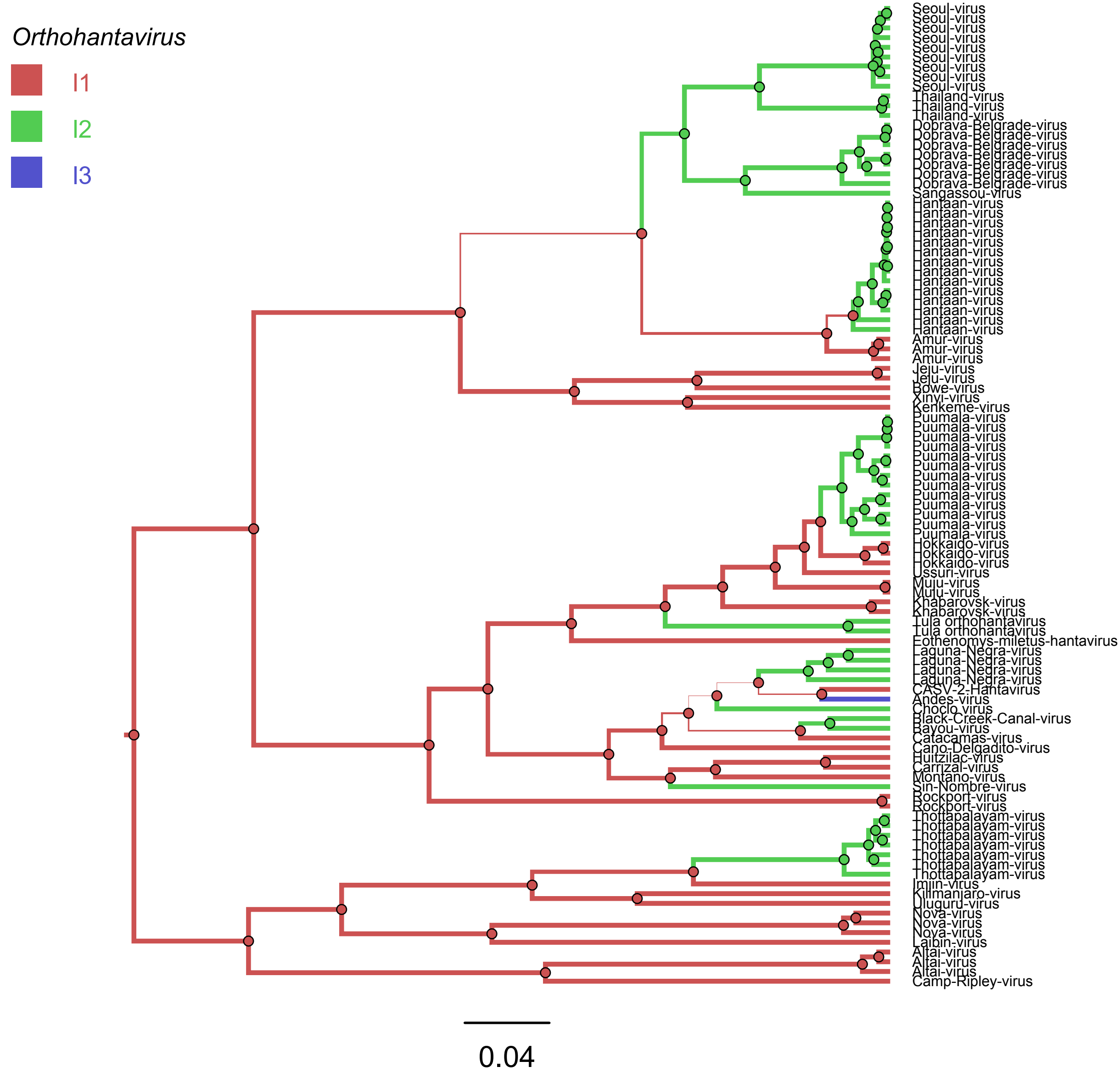

Orthohantavirus
I1
I2
I3
Seoul-virus
Thailand-virus
Dobrava-Belgrade-virus
Sangassou-virus
Hantaan-virus
Amur-virus
Jeju-virus
Bowe-virus
Xinyi-virus
Kenkeme-virus
Puumala-virus
Hokkaido-virus
Ussuri-virus
Muju-virus
Khabarovsk-virus
Tula orthohantavirus
Eothenomys-miletus-hantavirus
Laguna-Negra-virus
CASV-2-Hantavirus
Andes-virus
Choclo virus
Black-Creek-Canal-virus
Bayou-virus
Catacamas-virus
Cano-Delgadito-virus
Huitzilac-virus
Carrizal-virus
Montano-virus
Sin-Nombre-virus
Rockport-virus
Thottapalayam-virus
Imjin-virus
Kilimanjaro-virus
Uluguru-virus
Nova-virus
Laibin-virus
Altai-virus
Camp-Ripley-virus
0.04

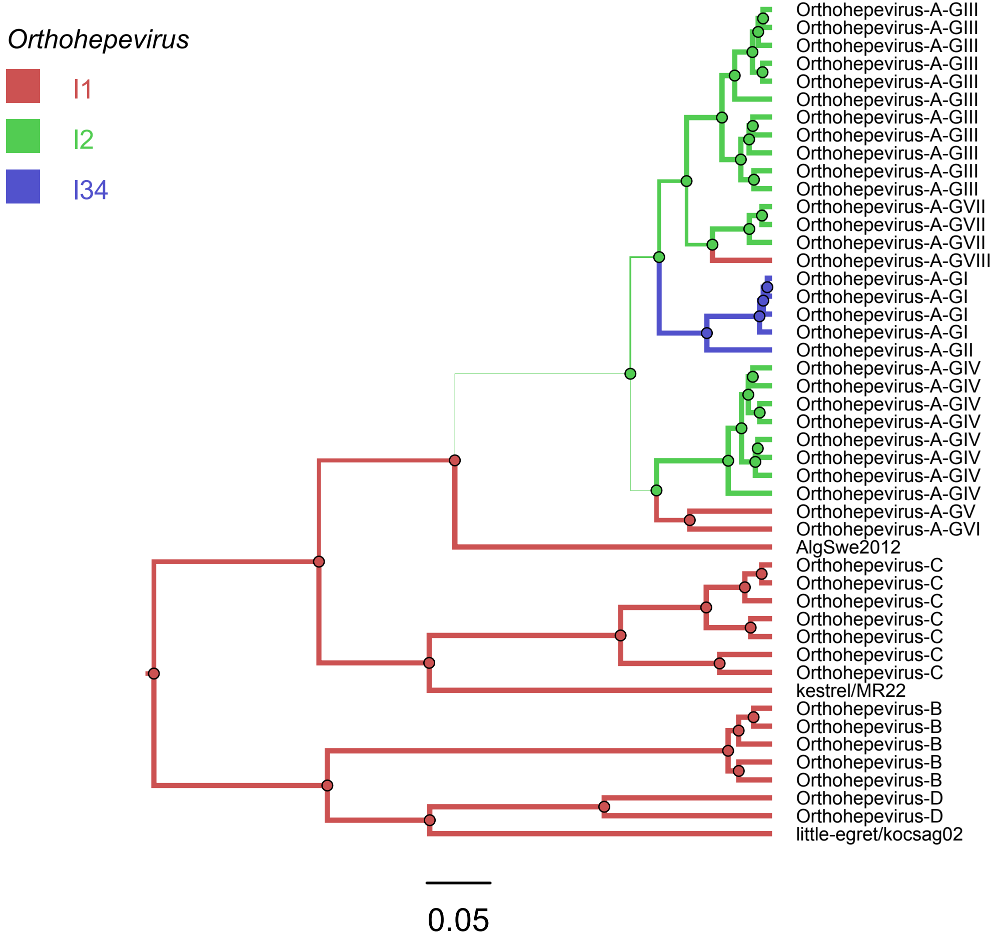

Orthohepevirus
I1
I2
I34
Orthohepevirus-A-GIII
Orthohepevirus-A-GIII
Orthohepevirus-A-GIII
Orthohepevirus-A-GIII
Orthohepevirus-A-GIII
Orthohepevirus-A-GIII
Orthohepevirus-A-GIII
Orthohepevirus-A-GIII
Orthohepevirus-A-GIII
Orthohepevirus-A-GIII
Orthohepevirus-A-GIII
Orthohepevirus-A-GVII
Orthohepevirus-A-GVII
Orthohepevirus-A-GVII
Orthohepevirus-A-GVIII
Orthohepevirus-A-GI
Orthohepevirus-A-GI
Orthohepevirus-A-GI
Orthohepevirus-A-GI
Orthohepevirus-A-GII
Orthohepevirus-A-GIV
Orthohepevirus-A-GIV
Orthohepevirus-A-GIV
Orthohepevirus-A-GIV
Orthohepevirus-A-GIV
Orthohepevirus-A-GIV
Orthohepevirus-A-GIV
Orthohepevirus-A-GIV
Orthohepevirus-A-GV
Orthohepevirus-A-GVI
AlgSwe2012
Orthohepevirus-C
Orthohepevirus-C
Orthohepevirus-C
Orthohepevirus-C
Orthohepevirus-C
Orthohepevirus-C
Orthohepevirus-C
kestrel/MR22
Orthohepevirus-B
Orthohepevirus-B
Orthohepevirus-B
Orthohepevirus-B
Orthohepevirus-B
Orthohepevirus-D
Orthohepevirus-D
little-egret/kocsag02
0.05

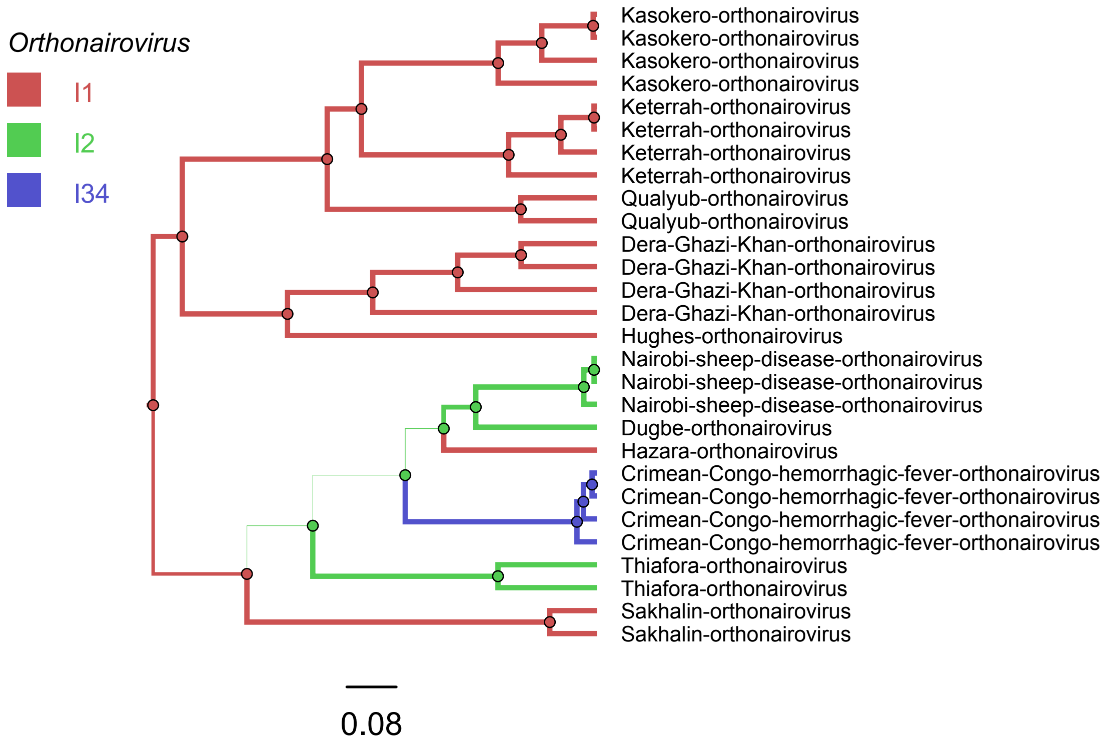
Orthonairovirus
I1
I2
I34
Kasokero-orthonairovirus
Kasokero-orthonairovirus
Kasokero-orthonairovirus
Kasokero-orthonairovirus
Keterrah-orthonairovirus
Keterrah-orthonairovirus
Keterrah-orthonairovirus
Keterrah-orthonairovirus
Qualyub-orthonairovirus
Qualyub-orthonairovirus
Dera-Ghazi-Khan-orthonairovirus
Dera-Ghazi-Khan-orthonairovirus
Dera-Ghazi-Khan-orthonairovirus
Dera-Ghazi-Khan-orthonairovirus
Hughes-orthonairovirus
Nairobi-sheep-disease-orthonairovirus
Nairobi-sheep-disease-orthonairovirus
Nairobi-sheep-disease-orthonairovirus
Dugbe-orthonairovirus
Hazara-orthonairovirus
Crimean-Congo-hemorrhagic-fever-orthonairovirus
Crimean-Congo-hemorrhagic-fever-orthonairovirus
Crimean-Congo-hemorrhagic-fever-orthonairovirus
Crimean-Congo-hemorrhagic-fever-orthonairovirus
Thiafora-orthonairovirus
Thiafora-orthonairovirus
Sakhalin-orthonairovirus
Sakhalin-orthonairovirus
0.08

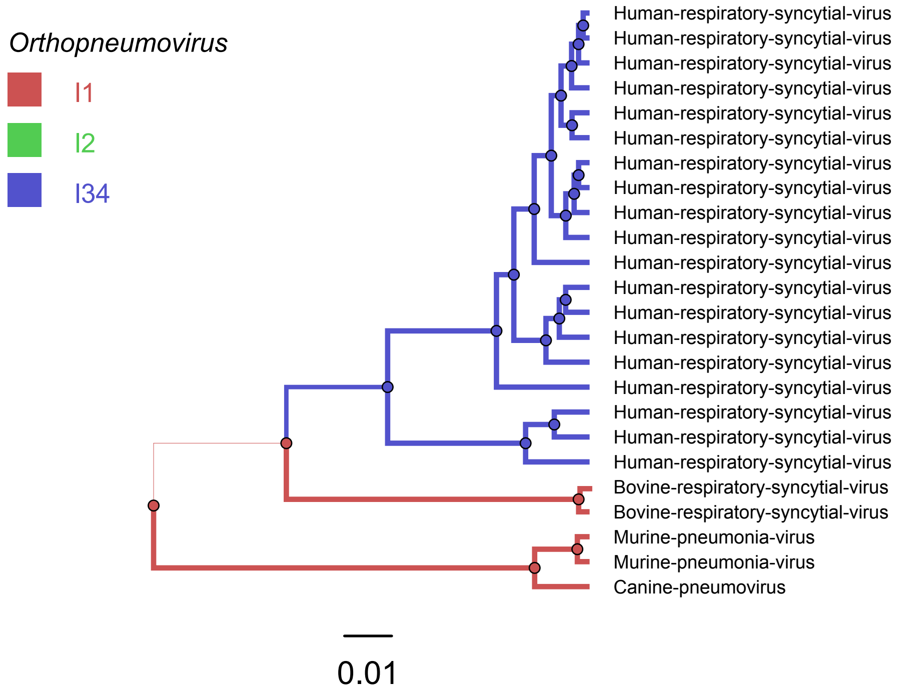

Orthopneumovirus
I1
I2
I34
Human-respiratory-syncytial-virus
Human-respiratory-syncytial-virus
Human-respiratory-syncytial-virus
Human-respiratory-syncytial-virus
Human-respiratory-syncytial-virus
Human-respiratory-syncytial-virus
Human-respiratory-syncytial-virus
Human-respiratory-syncytial-virus
Human-respiratory-syncytial-virus
Human-respiratory-syncytial-virus
Human-respiratory-syncytial-virus
Human-respiratory-syncytial-virus
Human-respiratory-syncytial-virus
Human-respiratory-syncytial-virus
Human-respiratory-syncytial-virus
Human-respiratory-syncytial-virus
Human-respiratory-syncytial-virus
Human-respiratory-syncytial-virus
Human-respiratory-syncytial-virus
Bovine-respiratory-syncytial-virus
Bovine-respiratory-syncytial-virus
Murine-pneumonia-virus
Murine-pneumonia-virus
Canine-pneumovirus
0.01

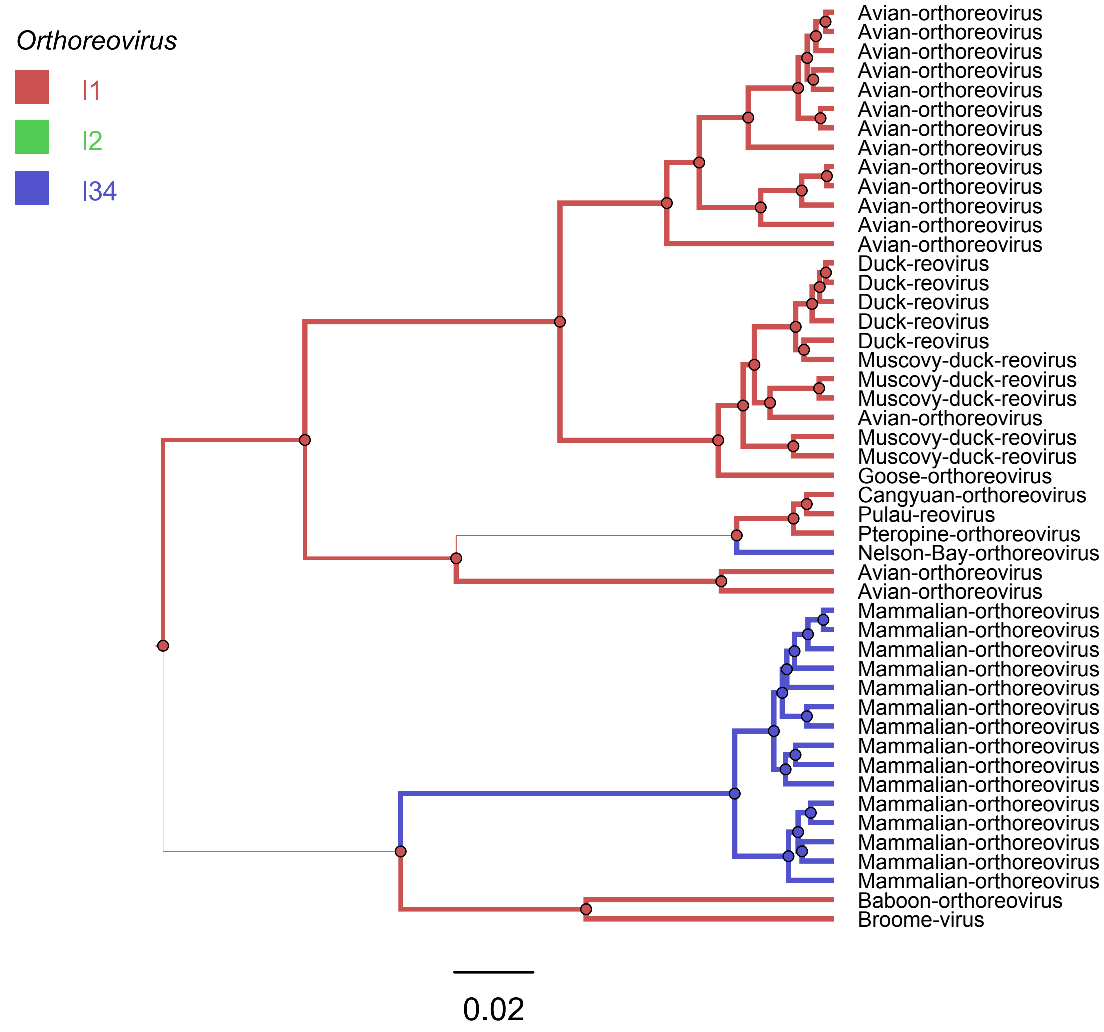

Orthoreovirus
I1
I2
I34
Avian-orthoreovirus
Avian-orthoreovirus
Avian-orthoreovirus
Avian-orthoreovirus
Avian-orthoreovirus
Avian-orthoreovirus
Avian-orthoreovirus
Avian-orthoreovirus
Avian-orthoreovirus
Avian-orthoreovirus
Avian-orthoreovirus
Avian-orthoreovirus
Avian-orthoreovirus
Duck-reovirus
Duck-reovirus
Duck-reovirus
Duck-reovirus
Duck-reovirus
Muscovy-duck-reovirus
Muscovy-duck-reovirus
Muscovy-duck-reovirus
Avian-orthoreovirus
Muscovy-duck-reovirus
Muscovy-duck-reovirus
Goose-orthoreovirus
Cangyuan-orthoreovirus
Pulau-reovirus
Pteropine-orthoreovirus
Nelson-Bay-orthoreovirus
Avian-orthoreovirus
Avian-orthoreovirus
Mammalian-orthoreovirus
Mammalian-orthoreovirus
Mammalian-orthoreovirus
Mammalian-orthoreovirus
Mammalian-orthoreovirus
Mammalian-orthoreovirus
Mammalian-orthoreovirus
Mammalian-orthoreovirus
Mammalian-orthoreovirus
Mammalian-orthoreovirus
Mammalian-orthoreovirus
Mammalian-orthoreovirus
Mammalian-orthoreovirus
Mammalian-orthoreovirus
Mammalian-orthoreovirus
Baboon-orthoreovirus
Broome-virus
0.02

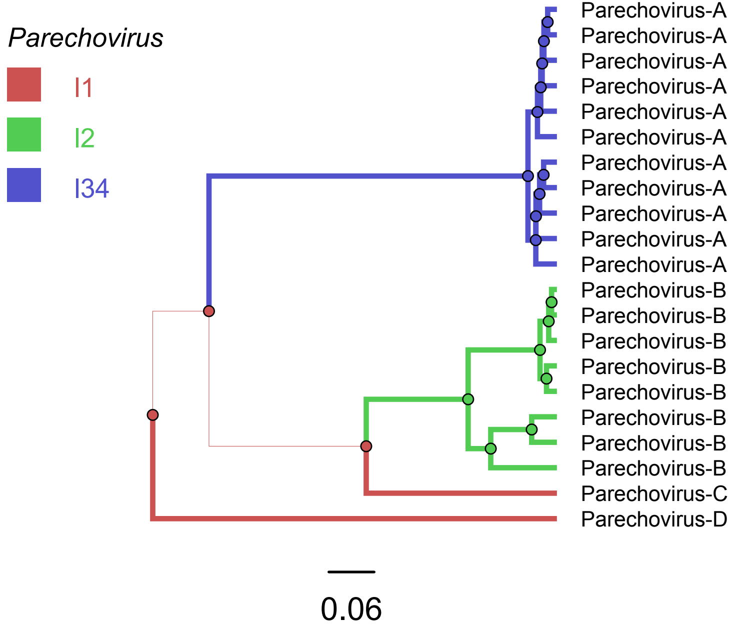
Parechovirus
I1
I2
I34
Parechovirus-A
Parechovirus-A
Parechovirus-A
Parechovirus-A
Parechovirus-A
Parechovirus-A
Parechovirus-A
Parechovirus-A
Parechovirus-A
Parechovirus-A
Parechovirus-A
Parechovirus-B
Parechovirus-B
Parechovirus-B
Parechovirus-B
Parechovirus-B
Parechovirus-B
Parechovirus-B
Parechovirus-B
Parechovirus-C
Parechovirus-D
0.06

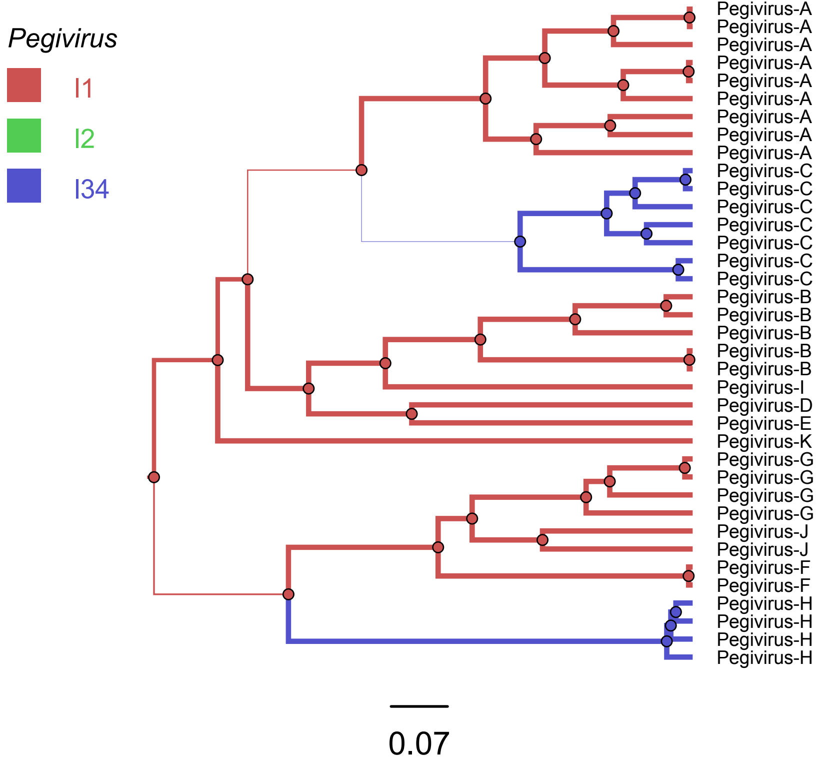

Pegivirus
I1
I2
I34
Pegivirus-A
Pegivirus-A
Pegivirus-A
Pegivirus-A
Pegivirus-A
Pegivirus-A
Pegivirus-A
Pegivirus-A
Pegivirus-A
Pegivirus-C
Pegivirus-C
Pegivirus-C
Pegivirus-C
Pegivirus-C
Pegivirus-C
Pegivirus-C
Pegivirus-B
Pegivirus-B
Pegivirus-B
Pegivirus-B
Pegivirus-B
Pegivirus-I
Pegivirus-D
Pegivirus-E
Pegivirus-K
Pegivirus-G
Pegivirus-G
Pegivirus-G
Pegivirus-G
Pegivirus-J
Pegivirus-J
Pegivirus-F
Pegivirus-F
Pegivirus-H
Pegivirus-H
Pegivirus-H
Pegivirus-H
0.07

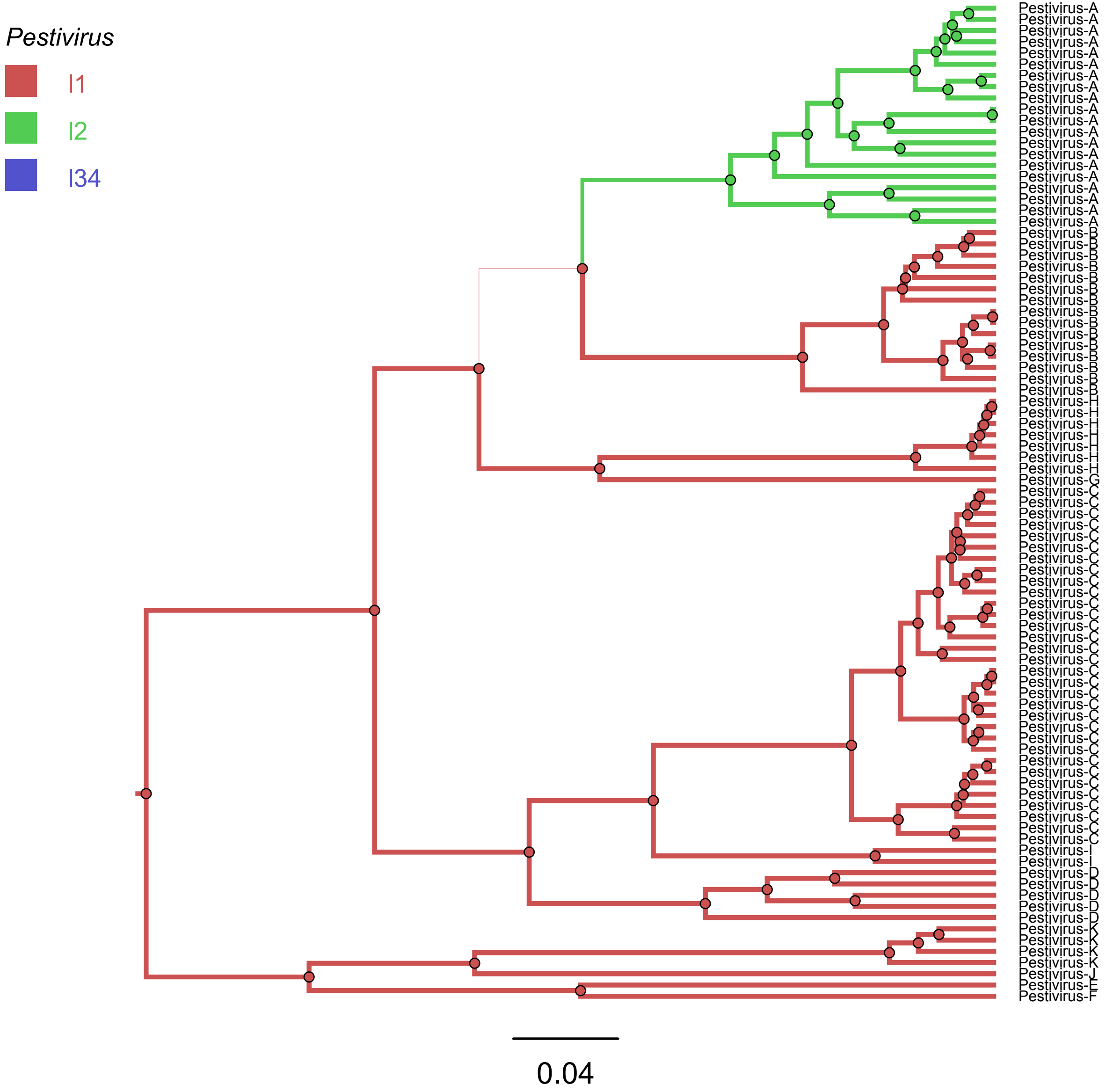

Pestivirus
I1
I2
I34
Pestivirus-A
Pestivirus-A
Pestivirus-A
Pestivirus-A
Pestivirus-A
Pestivirus-A
Pestivirus-A
Pestivirus-A
Pestivirus-A
Pestivirus-A
Pestivirus-A
Pestivirus-A
Pestivirus-A
Pestivirus-A
Pestivirus-A
Pestivirus-A
Pestivirus-A
Pestivirus-A
Pestivirus-A
Pestivirus-A
Pestivirus-B
Pestivirus-B
Pestivirus-B
Pestivirus-B
Pestivirus-B
Pestivirus-B
Pestivirus-B
Pestivirus-B
Pestivirus-B
Pestivirus-B
Pestivirus-B
Pestivirus-B
Pestivirus-B
Pestivirus-B
Pestivirus-B
Pestivirus-H
Pestivirus-H
Pestivirus-H
Pestivirus-H
Pestivirus-H
Pestivirus-H
Pestivirus-H
Pestivirus-G
Pestivirus-C
Pestivirus-C
Pestivirus-C
Pestivirus-C
Pestivirus-C
Pestivirus-C
Pestivirus-C
Pestivirus-C
Pestivirus-C
Pestivirus-C
Pestivirus-C
Pestivirus-C
Pestivirus-C
Pestivirus-C
Pestivirus-C
Pestivirus-C
Pestivirus-C
Pestivirus-C
Pestivirus-C
Pestivirus-C
Pestivirus-C
Pestivirus-C
Pestivirus-C
Pestivirus-C
Pestivirus-C
Pestivirus-C
Pestivirus-C
Pestivirus-C
Pestivirus-C
Pestivirus-C
Pestivirus-C
Pestivirus-C
Pestivirus-I
Pestivirus-I
Pestivirus-D
Pestivirus-D
Pestivirus-D
Pestivirus-D
Pestivirus-D
Pestivirus-K
Pestivirus-K
Pestivirus-K
Pestivirus-K
Pestivirus-J
Pestivirus-E
Pestivirus-F
0.04

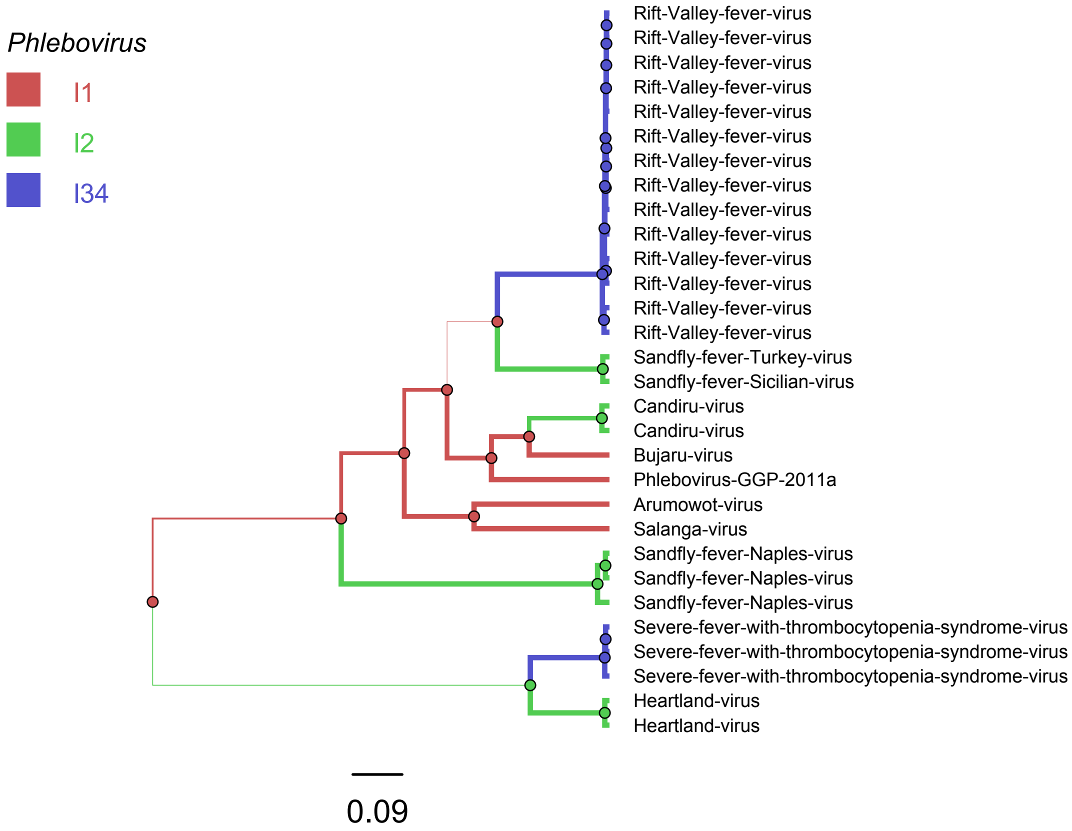
Phlebovirus
I1
I2
I34
Rift-Valley-fever-virus
Rift-Valley-fever-virus
Rift-Valley-fever-virus
Rift-Valley-fever-virus
Rift-Valley-fever-virus
Rift-Valley-fever-virus
Rift-Valley-fever-virus
Rift-Valley-fever-virus
Rift-Valley-fever-virus
Rift-Valley-fever-virus
Rift-Valley-fever-virus
Rift-Valley-fever-virus
Rift-Valley-fever-virus
Rift-Valley-fever-virus
Sandfly-fever-Turkey-virus
Sandfly-fever-Sicilian-virus
Candiru-virus
Candiru-virus
Bujaru-virus
Phlebovirus-GGP-2011a
Arumowot-virus
Salanga-virus
Sandfly-fever-Naples-virus
Sandfly-fever-Naples-virus
Sandfly-fever-Naples-virus
Severe-fever-with-thrombocytopenia-syndrome-virus
Severe-fever-with-thrombocytopenia-syndrome-virus
Severe-fever-with-thrombocytopenia-syndrome-virus
Heartland-virus
Heartland-virus
0.09

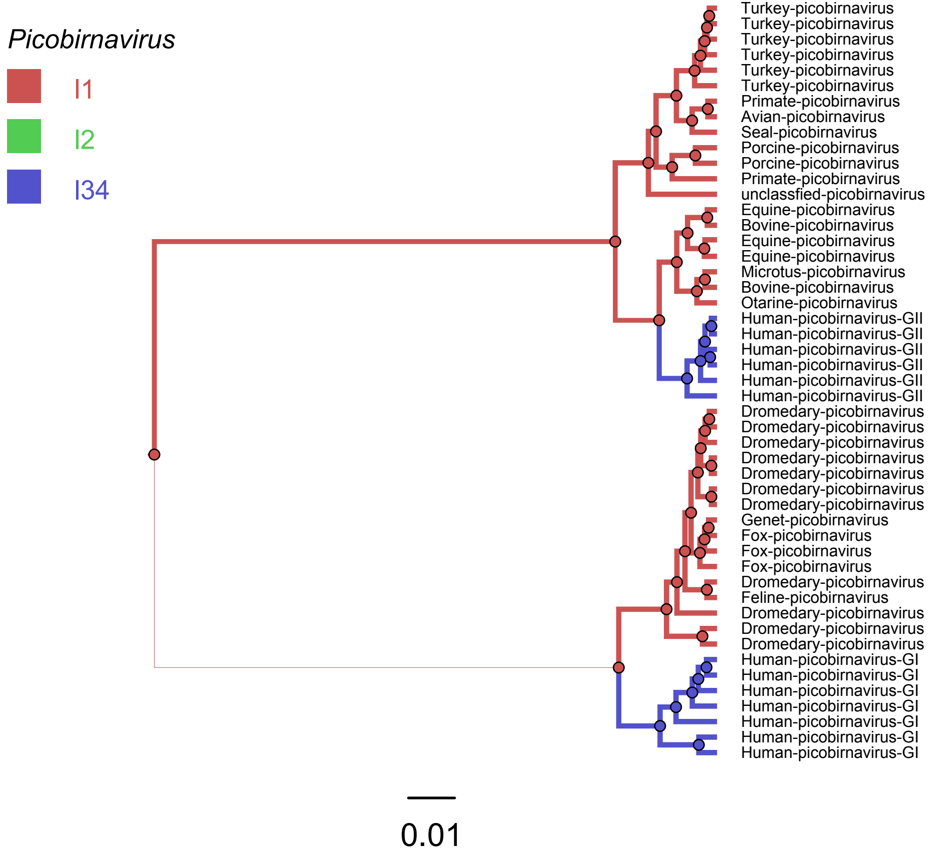

Picobirnavirus
I1
I2
I34
Turkey-picobirnavirus
Turkey-picobirnavirus
Turkey-picobirnavirus
Turkey-picobirnavirus
Turkey-picobirnavirus
Turkey-picobirnavirus
Primate-picobirnavirus
Avian-picobirnavirus
Seal-picobirnavirus
Porcine-picobirnavirus
Porcine-picobirnavirus
Primate-picobirnavirus
unclassfied-picobirnavirus
Equine-picobirnavirus
Bovine-picobirnavirus
Equine-picobirnavirus
Equine-picobirnavirus
Microtus-picobirnavirus
Bovine-picobirnavirus
Otarine-picobirnavirus
Human-picobirnavirus-GII
Human-picobirnavirus-GII
Human-picobirnavirus-GII
Human-picobirnavirus-GII
Human-picobirnavirus-GII
Human-picobirnavirus-GII
Dromedary-picobirnavirus
Dromedary-picobirnavirus
Dromedary-picobirnavirus
Dromedary-picobirnavirus
Dromedary-picobirnavirus
Dromedary-picobirnavirus
Dromedary-picobirnavirus
Genet-picobirnavirus
Fox-picobirnavirus
Fox-picobirnavirus
Fox-picobirnavirus
Dromedary-picobirnavirus
Feline-picobirnavirus
Dromedary-picobirnavirus
Dromedary-picobirnavirus
Dromedary-picobirnavirus
Human-picobirnavirus-GI
Human-picobirnavirus-GI
Human-picobirnavirus-GI
Human-picobirnavirus-GI
Human-picobirnavirus-GI
Human-picobirnavirus-GI
Human-picobirnavirus-GI
0.01

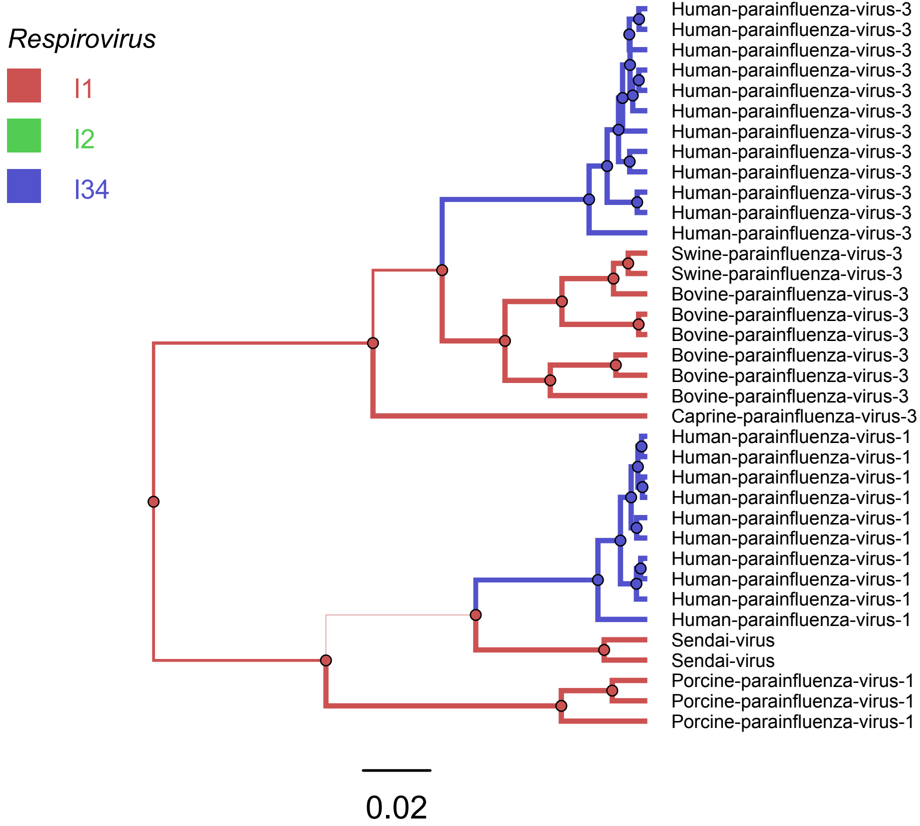

Respirovirus
I1
I2
I34
Human-parainfluenza-virus-3
Human-parainfluenza-virus-3
Human-parainfluenza-virus-3
Human-parainfluenza-virus-3
Human-parainfluenza-virus-3
Human-parainfluenza-virus-3
Human-parainfluenza-virus-3
Human-parainfluenza-virus-3
Human-parainfluenza-virus-3
Human-parainfluenza-virus-3
Human-parainfluenza-virus-3
Human-parainfluenza-virus-3
Swine-parainfluenza-virus-3
Swine-parainfluenza-virus-3
Bovine-parainfluenza-virus-3
Bovine-parainfluenza-virus-3
Bovine-parainfluenza-virus-3
Bovine-parainfluenza-virus-3
Bovine-parainfluenza-virus-3
Bovine-parainfluenza-virus-3
Caprine-parainfluenza-virus-3
Human-parainfluenza-virus-1
Human-parainfluenza-virus-1
Human-parainfluenza-virus-1
Human-parainfluenza-virus-1
Human-parainfluenza-virus-1
Human-parainfluenza-virus-1
Human-parainfluenza-virus-1
Human-parainfluenza-virus-1
Human-parainfluenza-virus-1
Human-parainfluenza-virus-1
Sendai-virus
Sendai-virus
Porcine-parainfluenza-virus-1
Porcine-parainfluenza-virus-1
Porcine-parainfluenza-virus-1
0.02

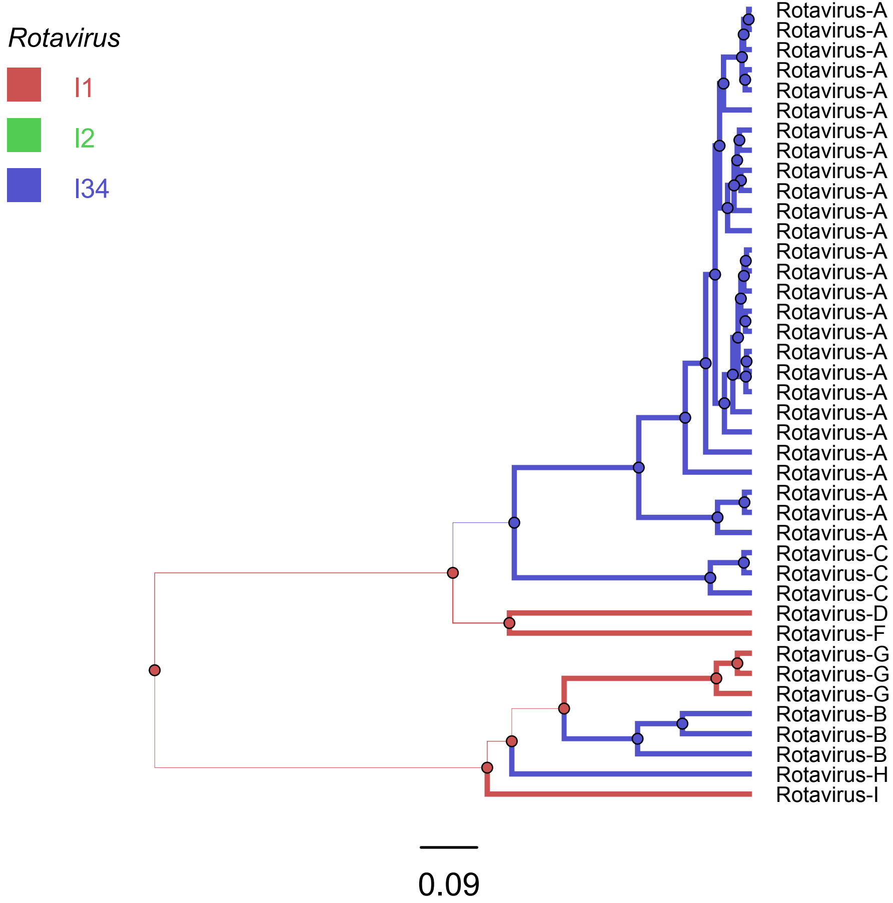
Rotavirus
I1
I2
I34
Rotavirus-A
Rotavirus-A
Rotavirus-A
Rotavirus-A
Rotavirus-A
Rotavirus-A
Rotavirus-A
Rotavirus-A
Rotavirus-A
Rotavirus-A
Rotavirus-A
Rotavirus-A
Rotavirus-A
Rotavirus-A
Rotavirus-A
Rotavirus-A
Rotavirus-A
Rotavirus-A
Rotavirus-A
Rotavirus-A
Rotavirus-A
Rotavirus-A
Rotavirus-A
Rotavirus-A
Rotavirus-A
Rotavirus-A
Rotavirus-A
Rotavirus-A
Rotavirus-C
Rotavirus-C
Rotavirus-C
Rotavirus-D
Rotavirus-F
Rotavirus-G
Rotavirus-G
Rotavirus-G
Rotavirus-B
Rotavirus-B
Rotavirus-B
Rotavirus-H
Rotavirus-I
0.09

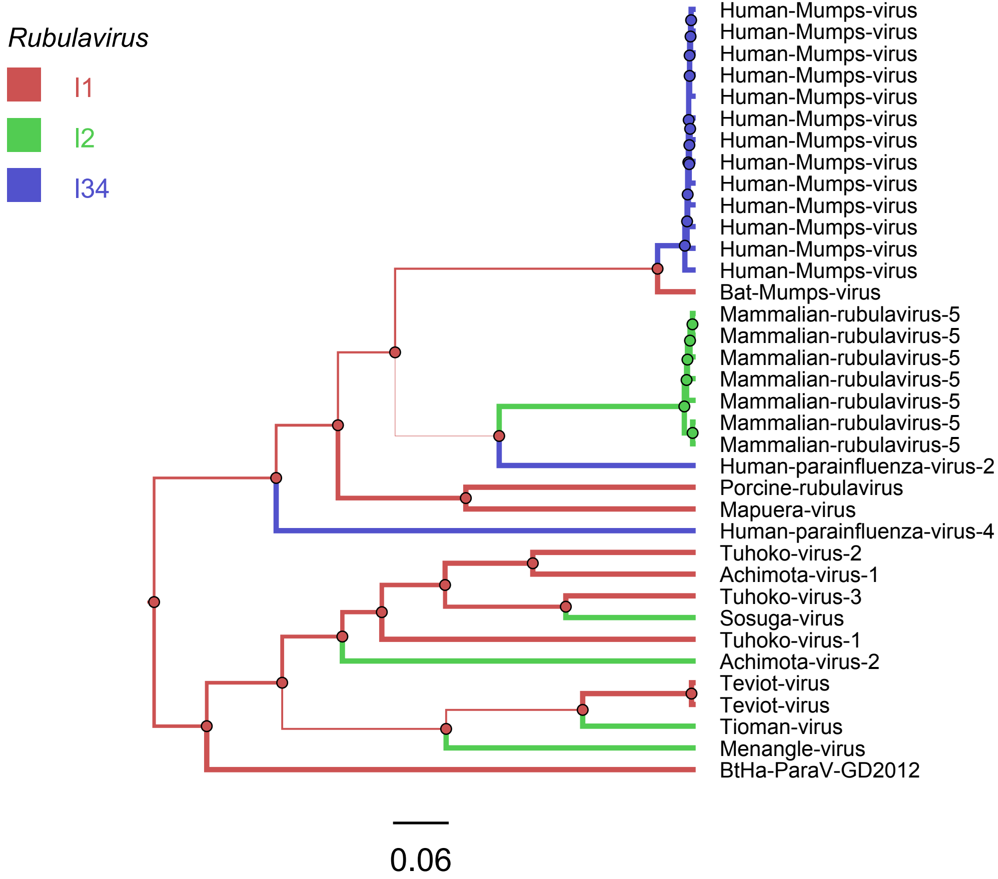
Rubulavirus
I1
I2
I34
Human-Mumps-virus
Human-Mumps-virus
Human-Mumps-virus
Human-Mumps-virus
Human-Mumps-virus
Human-Mumps-virus
Human-Mumps-virus
Human-Mumps-virus
Human-Mumps-virus
Human-Mumps-virus
Human-Mumps-virus
Human-Mumps-virus
Human-Mumps-virus
Bat-Mumps-virus
Mammalian-rubulavirus-5
Mammalian-rubulavirus-5
Mammalian-rubulavirus-5
Mammalian-rubulavirus-5
Mammalian-rubulavirus-5
Mammalian-rubulavirus-5
Mammalian-rubulavirus-5
Human-parainfluenza-virus-2
Porcine-rubulavirus
Mapuera-virus
Human-parainfluenza-virus-4
Tuhoko-virus-2
Achimota-virus-1
Tuhoko-virus-3
Sosuga-virus
Tuhoko-virus-1
Achimota-virus-2
Teviot-virus
Teviot-virus
Tioman-virus
Menangle-virus
BtHa-ParaV-GD2012
0.06

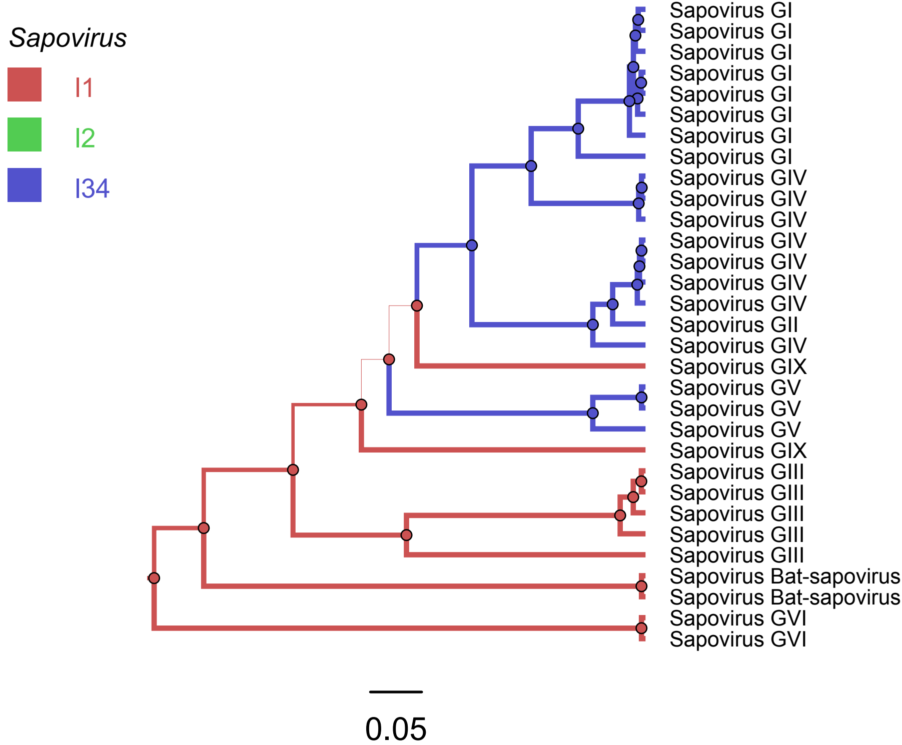
Sapovirus
I1
I2
I34
Sapovirus GI
Sapovirus GI
Sapovirus GI
Sapovirus GI
Sapovirus GI
Sapovirus GI
Sapovirus GI
Sapovirus GI
Sapovirus GIV
Sapovirus GIV
Sapovirus GIV
Sapovirus GIV
Sapovirus GIV
Sapovirus GIV
Sapovirus GIV
Sapovirus GII
Sapovirus GIV
Sapovirus GIX
Sapovirus GV
Sapovirus GV
Sapovirus GV
Sapovirus GIX
Sapovirus GIII
Sapovirus GIII
Sapovirus GIII
Sapovirus GIII
Sapovirus GIII
Sapovirus Bat-sapovirus
Sapovirus Bat-sapovirus
Sapovirus GVI
Sapovirus GVI
0.05

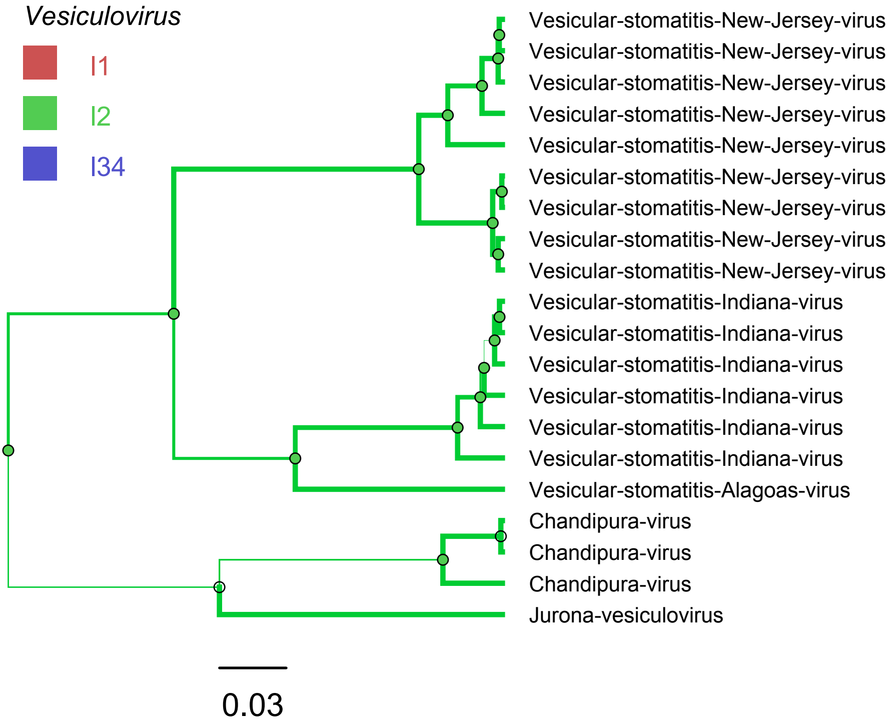
Vesiculovirus
I1
I2
I34
Vesicular-stomatitis-New-Jersey-virus
Vesicular-stomatitis-New-Jersey-virus
Vesicular-stomatitis-New-Jersey-virus
Vesicular-stomatitis-New-Jersey-virus
Vesicular-stomatitis-New-Jersey-virus
Vesicular-stomatitis-New-Jersey-virus
Vesicular-stomatitis-New-Jersey-virus
Vesicular-stomatitis-New-Jersey-virus
Vesicular-stomatitis-New-Jersey-virus
Vesicular-stomatitis-Indiana-virus
Vesicular-stomatitis-Indiana-virus
Vesicular-stomatitis-Indiana-virus
Vesicular-stomatitis-Indiana-virus
Vesicular-stomatitis-Indiana-virus
Vesicular-stomatitis-Indiana-virus
Vesicular-stomatitis-Alagoas-virus
Chandipura-virus
Chandipura-virus
Chandipura-virus
Jurona-vesiculovirus
0.03

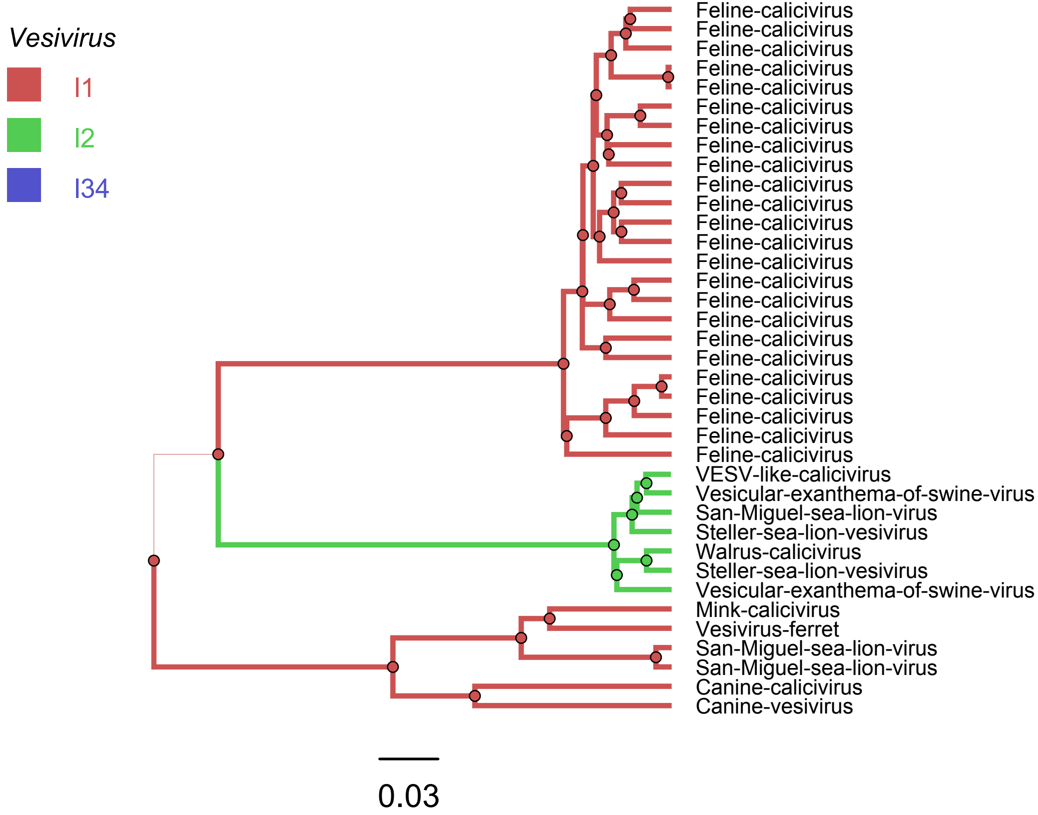
Vesivirus
I1
I2
I34
Feline-calicivirus
Feline-calicivirus
Feline-calicivirus
Feline-calicivirus
Feline-calicivirus
Feline-calicivirus
Feline-calicivirus
Feline-calicivirus
Feline-calicivirus
Feline-calicivirus
Feline-calicivirus
Feline-calicivirus
Feline-calicivirus
Feline-calicivirus
Feline-calicivirus
Feline-calicivirus
Feline-calicivirus
Feline-calicivirus
Feline-calicivirus
Feline-calicivirus
Feline-calicivirus
Feline-calicivirus
Feline-calicivirus
Feline-calicivirus
VESV-like-calicivirus
Vesicular-exanthema-of-swine-virus
San-Miguel-sea-lion-virus
Steller-sea-lion-vesivirus
Walrus-calicivirus
Steller-sea-lion-vesivirus
Vesicular-exanthema-of-swine-virus
Mink-calicivirus
Vesivirus-ferret
San-Miguel-sea-lion-virus
San-Miguel-sea-lion-virus
Canine-calicivirus
Canine-vesivirus
0.03
